# Purging of highly deleterious mutations through severe bottlenecks in Alpine ibex

**DOI:** 10.1101/605147

**Authors:** Christine Grossen, Frederic Guillaume, Lukas F. Keller, Daniel Croll

**Author notes:** Author contributions:* Conception and design of study: CG, LFK, DC Acquisition and analysis of data: CG Interpretation of data: CG, FG, LFK, DC Funding: CG, LFK Wrote the manuscript with input from the other authors: CG, DC.

## Abstract

Human activity caused dramatic population declines in many wild species. The resulting bottlenecks have a profound impact on the genetic makeup of a species with unknown consequences for health. A key genetic factor for species survival is the evolution of deleterious mutation load, but how bottleneck strength and mutation load interact lacks empirical evidence. Here, we take advantage of the exceptionally well-characterized population bottlenecks of the once nearly extinct Alpine ibex. The species survived one of the most dramatic bottlenecks known for successfully restored species. We analyze 60 complete genomes of six ibex species and the domestic goat. We show that historic bottlenecks rather than the current conservation status predict levels of genome-wide variation. By retracing the recolonization of the Alps by Alpine ibex, we find genomic evidence of concurrent purging of highly deleterious mutations but accumulation of mildly deleterious mutations. This demonstrates how human-induced severe bottlenecks caused both relaxed selection and purging, thus reshaping the landscape of deleterious mutation load. Our findings also highlight that even populations of ~1000 individuals can accumulate mildly deleterious mutations. Hence, conservation efforts should focus on preventing population declines below such levels to ensure long-term survival of species.

## Main text

Climate change and pressure from human activities such as hunting caused profound changes in the population and demographic structure of many species ^1^. This is because extinction events and subsequent recolonization severely alter the genetic makeup ^2^. The demographic changes have important consequences for wildlife management and the conservation of endangered species ^2^ including raising the risk of genetic disorders ^e.g 3–9^. However, nearly all plant and animal populations including humans suffered from temporary reductions in population size – so-called bottlenecks. Bottlenecks increase genetic drift and inbreeding, which leads to a loss of genetic variation, reduces the efficacy of natural selection, and increases the expression of deleterious recessive mutations ^10–12^. The expression of recessive mutations under inbreeding creates the potential for selection to act against these mutations. This process known as purging reduces the frequency of deleterious mutations depending on the degree of dominance and the magnitude of the deleterious effects ^13^. Because purging depends on levels of inbreeding, bottlenecks tend to purge highly deleterious, recessive mutations unless population sizes are extremely low ^13–15^. Bottlenecks also increase genetic drift and reduce the efficacy of selection ^16^. This allows mildly deleterious mutations to drift to substantially higher frequencies ^4,6,17^. Hence, bottlenecks generate complex dynamics of deleterious mutation frequencies due to the independent effects of purging and reduced selection efficacy ^13–15,18,19^.

A major gap in our understanding is how reduced selection efficacy and purging jointly determine the mutation load in wild populations. Theoretical predictions are well established ^13,15,18,20^ but empirical evidence is conflicting ^9,21,22^ including for humans (see e.g. ^23–31^. Previous research used changes in fitness to infer possible purging events, ^7,20,32–34^, but changes in fitness can result from causes unrelated to purging ^12,35^. Direct evidence for purging exists only for isolated mountain gorilla populations that split off larger lowland populations ~20’000 years ago ^36^. However, it remains unknown how recent, dramatic bottleneck events on the scale caused by human activity impacts levels of deleterious mutations in the wild. Here, we take advantage of exceptionally well characterized repeated bottlenecks during the reintroduction of the once near-extinct Alpine ibex to retrace the fate of deleterious mutations. Alpine ibex were reduced to ~100 individuals in the 19th century in a single population in the Gran Paradiso region of Northern Italy ^37^. In less than a century, a census size of ca. 50’000 individuals has been re-established across the Alps. Thus, the population bottleneck of Alpine ibex is among the most dramatic recorded for any successfully restored species. Most extant populations experienced two to four, well-recorded bottlenecks leaving strong footprints of low genetic diversity ^38,39^.

We analyzed 60 genomes covering seven species including the Alpine ibex (*C. ibex*), five additional wild goats and the domestic goat. Siberian ibex have the largest population size (~200’000), followed by Alpine and Iberian ibex (both ~50’000), Bezoar (~25’000), Markhor (~6000) and Nubian ibex (~2500) ^40^. We found that Alpine ibex together with Iberian ibex and Markhor (*C. falconeri*) have exceptionally low genome-wide variation compared to closely related species (Figure 1, Figure S1). All three species either underwent severe bottlenecks in the past century or are currently threatened (Figure 1C, Figure S2, Tables S1 and S2). Historic records indicate that Alpine ibex suffered a bottleneck of ~100 individuals at the end of the 19th century and Iberian ibex a bottleneck of ~1000 individuals ^41^ (Table S1). In contrast, genome-wide diversity was highest in Siberian ibex (*C. sibirica*), which have large and relatively well-connected populations ^42^. The genomes of some Alpine ibex, the Markhor and some domestic goat individuals contained more than 20% genome-wide runs of homozygosity (ROH) (Figure 1D, Figure S3-S5, Table S3) ^43^. Overall, there was clear genomic evidence that the near extinction and recovery of the Alpine ibex resulted in substantial genetic drift and inbreeding.

**Figure 1:**
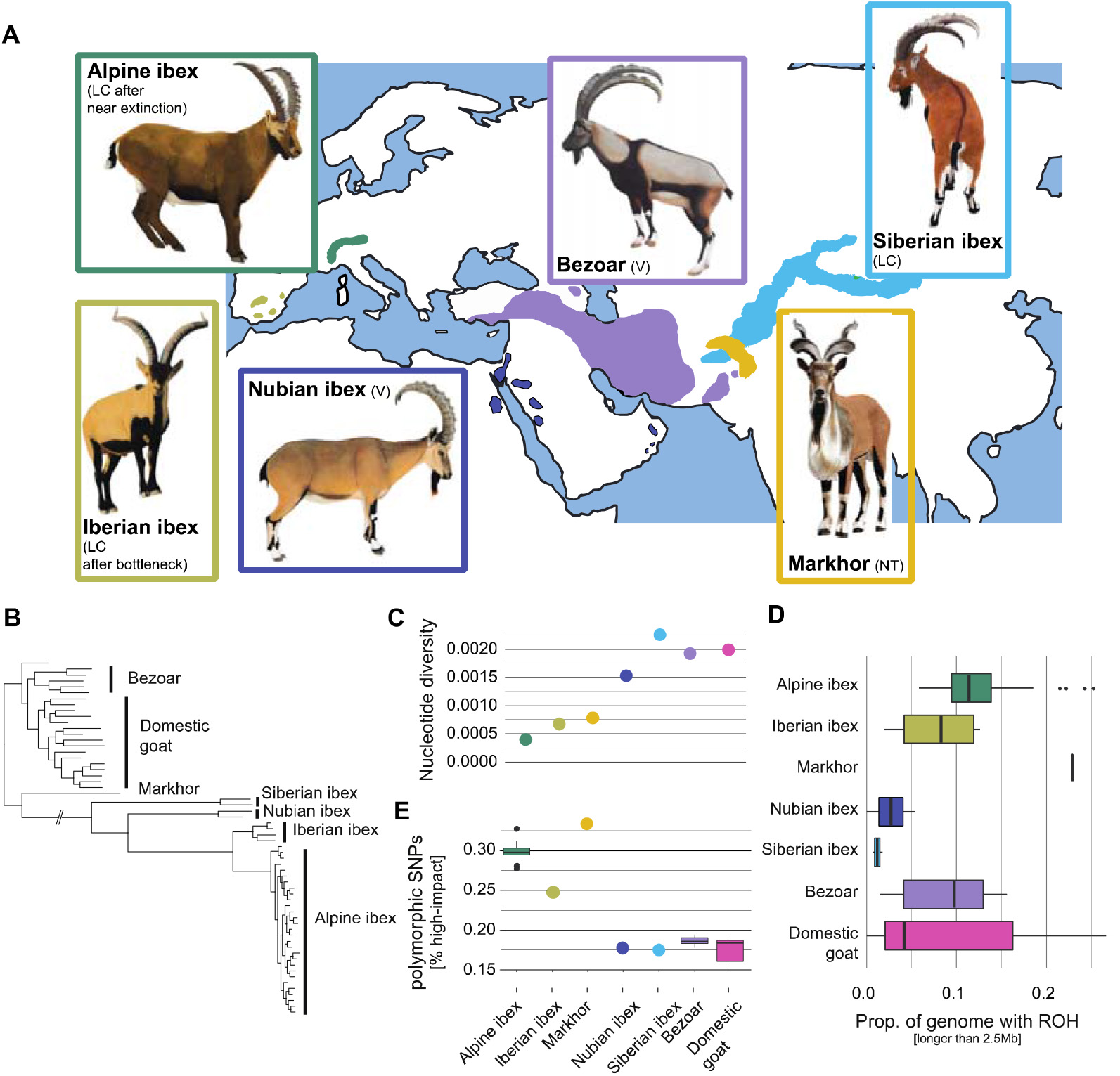
Genetic diversity and deleterious mutation load of ibex species. A) Geographical distribution and IUCN conservation status of ibex and wild goat species (LC: Least concern, V: Vulnerable, NT: Near threatened ^40^). Sample sizes: Alpine ibex (*C. ibex*): N=29, Iberian ibex (*C. pyrenaica*): N=4, Bezoar (*C. aegagrus*): N=6, Siberian ibex (*C. sibirica*): N=2, Markhor (*C. falconeri*): N=1, Nubian ibex (*C. nubiana*): N=2. B) Maximum likelihood phylogenetic analyses, C) nucleotide diversity, D) proportion of the genome with runs of homozygosity (ROH) longer than 2.5 Mb and E) percentage of polymorphic sites within species that segregate highly deleterious mutations. Confidence intervals are based on downsampling to four individuals matching the sample size of Iberian ibex (100 replicates).

We analyzed all *Capra* genomes for evidence of segregating deleterious mutations (Figure S6). We restricted our analyses to coding sequences with evidence for transcriptional activity in Alpine ibex organs and high evolutionary conservation (*i.e.* GERP, see methods) ^44^ yielding a total of 370’853 SNPs (Table S4). We found that across all seven *Capra* species 0.17% of these SNPs carried a highly deleterious variant with the majority of the highly deleterious variants incurring a stop-gain mutation, a further 19.1% carried a moderate impact variant and 33.1% carried a low impact variant (Table S4). We used seven additional mutation impact scoring approaches (see methods). The proportion of highly deleterious variants segregating within species was inversely correlated with nucleotide diversity (Figure 1C and E, Figure S7A-D; Pearson, df = 5, *r* = −0.86, *p* = 0.012). Hence, species with the smallest populations or the most severe population size reductions (*i.e.* Alpine ibex, Iberian ibex and Markhor) showed an accumulation of deleterious mutations relative to closely related species.

Both Alpine and Iberian ibex experienced severe bottlenecks due to overhunting and habitat fragmentation. We first analyzed evidence for purifying selection using allele frequency spectra. We focused only on derived sites that were polymorphic in at least one of the two sister species (Figure 2A, S8). We found that frequency distributions of high and moderate impact mutations in Alpine ibex were downwards shifted compared to modifier (*i.e.* neutral) mutations, which strongly suggests purifying selection against highly deleterious mutations (Figure 2B, D). Short indels (≤ 10 bp) in coding sequences revealed a similar downward shift (Figure S9). We found no comparable frequency shifts in Iberian ibex (Figure 2B). This is consistent with purifying selection acting more efficiently against highly deleterious mutations in Alpine ibex compared to Iberian ibex. We repeated the analyses of the site frequency spectra also for three alternative scoring systems (GERP, phyloP and phastCons) reporting phylogenetic conservation. We found for all three scores an excess in Alpine ibex of the most deleterious mutation category (Figure S10).

**Figure 2:**
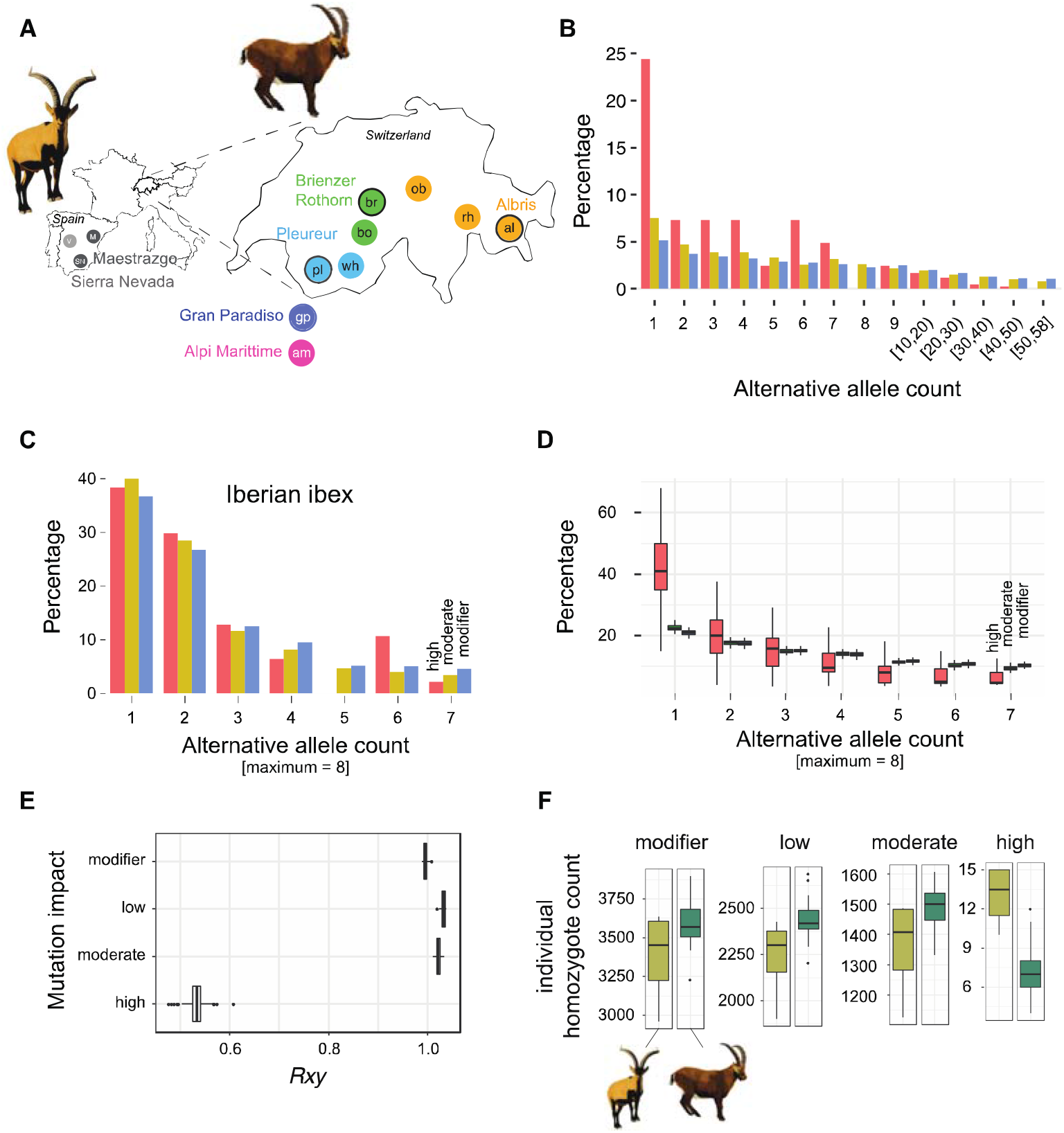
Deleterious mutations segregating in Alpine and Iberian ibex. A) Population sampling locations of Iberian ibex (left, grey circles) and Alpine ibex (right, colored circles). Each filled circle represents a population. Circles with a black outline indicate the first three reintroduced populations in Switzerland that were used for all subsequent population reintroductions of Alpine ibex. Colors associate founder and descendant populations (see also Figure 3A). (B) Site frequency spectrum for neutral (modifier), mildly (moderate impact) and highly deleterious (high impact) mutations for Alpine ibex. Alternative allele counts >10 are shown as mean counts per interval. (C) Site frequency spectrum for Iberian ibex. (D) Site frequency spectrum for Alpine ibex downsampled to four individuals matching the sample size of Iberian ibex (100 replicates). See Figure S11 for additional downsampling and jack-knifing analyses of data presented in B-D. E) *Rxy* analysis contrasting Iberian with Alpine ibex across the spectrum of impact categories. *Rxy* < 1 indicates a relative frequency deficit of the corresponding category in Alpine ibex compared to Iberian ibex. *Rxy* distributions are based on jack-knifing across chromosomes. All pairwise comparisons were significant except low vs. moderate (Tukey test, *p* < 0.0001). F) Individual homozygote counts per impact category for Iberian (light green) and Alpine ibex (dark green).

To test whether Alpine ibex showed evidence for purging of deleterious mutations, we calculated the relative number of derived alleles *Rxy* ^29^ for the different categories of mutations (Figure 2D). We used a random set of intergenic SNPs for standardization, which makes *Rxy* robust against sampling effects and population substructure ^29^. Low and moderate impact mutations (*i.e.* mildly deleterious mutations) showed a minor excess in Alpine ibex compared to Iberian ibex, indicating a higher load in Alpine ibex. In contrast, we found that highly deleterious mutations were strongly reduced in Alpine ibex compared to Iberian ibex (Tukey test, *p* < 0.0001, Figure 2E). Strikingly, the proportion of SNPs across the genome segregating a highly deleterious mutation is higher in Alpine ibex (Figure 1E), but *Rxy* shows that highly deleterious mutations have a pronounced downwards allele frequency shift in Alpine ibex compared to Iberian ibex. Furthermore, the number of homozygous sites with highly deleterious mutations per individual was significantly lower in Alpine ibex than Iberian ibex (*t* test, *p* = 0.015, Figure 2E). Individual allele counts at highly deleterious sites were also significantly lower in Alpine ibex compared to Iberian ibex (*t* test, *p* = 0.003). We assessed the robustness of the *Rxy* analyses using four additional mutation scoring methods (*i.e.* SIFT, REVEL, CADD and VEST3). We found that the highest impact category had a consistent deficit in Alpine ibex compared to Iberian ibex (Figure S12).. Together, this shows that highly deleterious mutations were substantially purged in Alpine ibex. We also found evidence for the accumulation of mildly deleterious mutations through genetic drift in Alpine ibex.

Consistent with the fact that all extant Alpine ibex originate from the Gran Paradiso, this population occupies the center of a principal component analysis (Figure 3A-B, Figure S13A-B; ^39^). The first populations re-established in the Alps were clearly differentiated from the Gran Paradiso source population and showed reduced nucleotide diversity due to reintroduction bottlenecks ^38^ (Figure 3A, C). These initial three reintroduced populations were used to establish additional populations raising the total number of experienced bottlenecks to 3 or 4. The additional bottlenecks led to further loss of nucleotide diversity and genetic drift, as indicated by the increasing spread in the principal component analysis (Figure 3A-C). An exceptional case constitutes the Alpi Marittime population, which was established through the translocation of 25 Gran Paradiso individuals of which only six successfully reproduced ^45^. As expected from such an extreme bottleneck, Alpi Marittime showed strong genetic differentiation from all other Alpine ibex populations and highly reduced nucleotide diversity (Figure 3B-C; ^46^). To compare the strength of drift experienced by different populations, we estimated long-term effective population sizes. For this, we used detailed demographic records spanning the near century since the populations were established ^47,48^. We found that nucleotide diversity decreased with smaller long-term population size (Spearman’s rank correlation, rho = 0.93, *p* = 0.007, Figure 3C). We found the same trend for the individual number of heterozygous sites per kilobase (Figure S14). Populations with the lowest harmonic mean population sizes also showed the highest levels of inbreeding. Genomes from the Gran Paradiso source population generally showed the lowest proportions of the genome affected by ROH, while reintroduced populations of lowest effective population size had the highest proportions of the genome affected by ROH (Pearson, df = 19, *r* = −0.70, *p* = 0.0004, Figure 3D and Figure S3).

**Figure 3:**
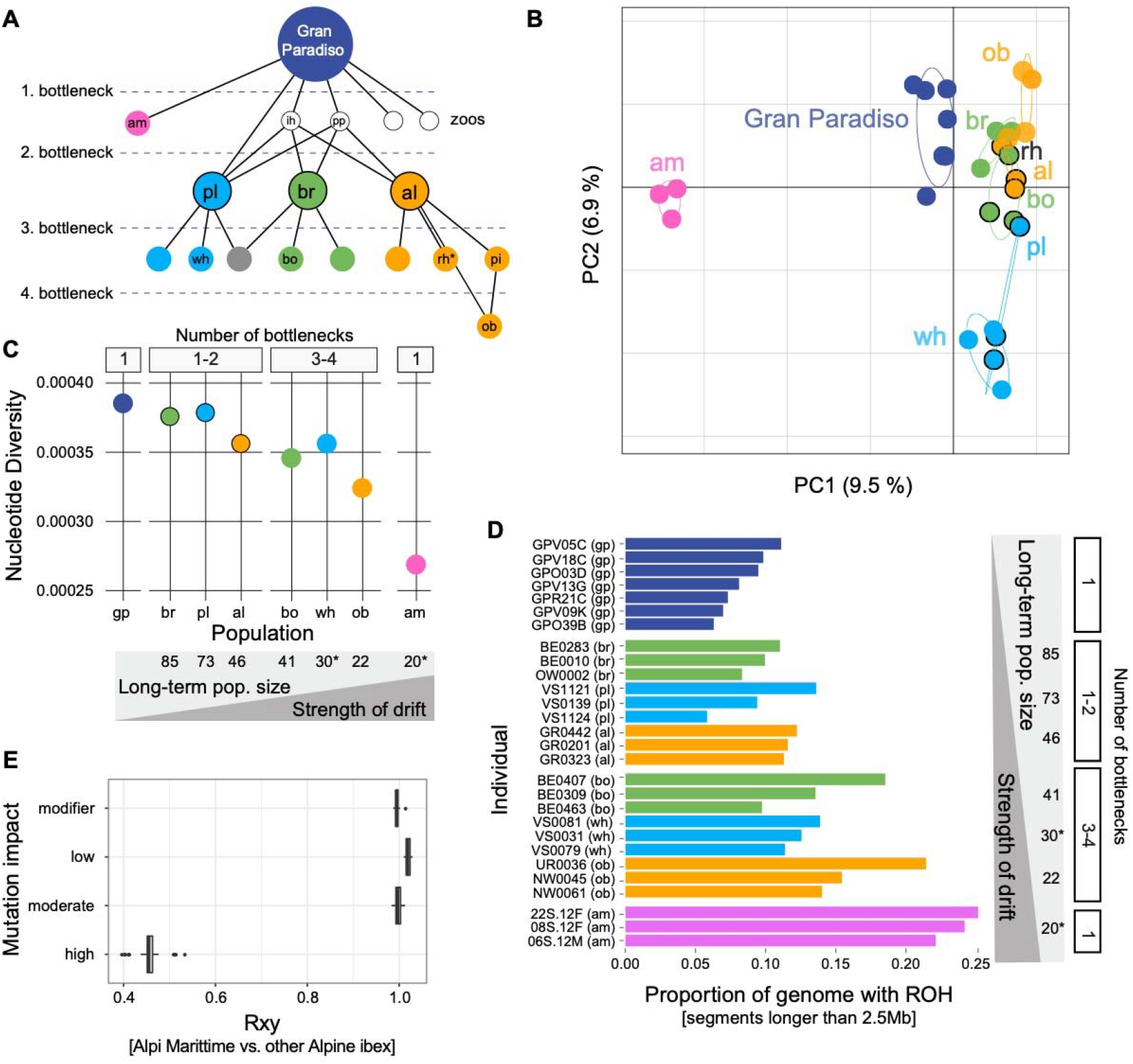
Genomic consequences of the Alpine ibex recolonization. A) The recolonization history and population pedigree of Alpine ibex. Locations include also zoos and the population Pilatus (pi), which was not sampled for this study. am: Alpi Marittime, gp: Gran Paradiso; ih: Zoo Interlaken Harder; al: Albris; bo: Bire O◻schinen; br: Brienzer Rothorn; ob: Oberbauenstock; pl: Pleureur; rh: Rheinwald; wh: Weisshorn; pi: Pilatus; pp: Wildpark Peter and Paul. The grey circle represents a population that was founded from more than one population. Figure elements were modified from Biebach and Keller (2009) with permission. B) Principal component analysis of all Alpine ibex individuals included in the study. C) Nucleotide diversity per population. D) Proportion of the genome within runs of homozygosity (ROH) longer than 2.5 Mb. E) *Rxy* analysis contrasting the strongly bottlenecked Alpi Marittime population with all other Alpine ibex populations across the spectrum of impact categories. *Rxy* < 1 indicates a relative frequency deficit of the corresponding category in the Alpi Marittime population. *Rxy* distributions are based on jack-knifing across chromosomes. Circles with a black outline indicate the first three reintroduced populations in Switzerland that were used for all subsequent population reintroductions of Alpine ibex. Colors associate founder and descendant populations.

Bottlenecks affect the landscape of deleterious mutations by randomly increasing or decreasing allele frequencies at individual loci. We find that individuals from populations that underwent stronger bottlenecks carry significantly more homozygotes for nearly neutral and mildly deleterious mutations (*i.e.* modifier, low and moderate impact mutations; Figure 4A). In contrast, individuals showed no meaningful difference in the number of homozygotes for highly deleterious (*i.e.* high impact) mutations across populations. The stability in the number of homozygotes for highly deleterious mutations through successive bottlenecks despite a step-wise increase in the number of homozygotes for weaker impact mutations, supports that purging occurred over the course of the Alpine ibex reintroductions. We repeated the analyses using an alternative scoring of mutations based on the phylogenetic conservation of the region in which the mutations were found (*i.e.* Genomic Evolutionary Rate Profiling; GERP). The number of homozygotes for mutations in highly conserved regions showed a slight upwards trend still indicating purifying but not necessarily purging for this category of mutations (Figure S15). This suggests that the mutational impact (*e.g.* premature stop codons) rather than degree of conservation predicts whether purging is likely to occur. This suggests also that mean fitness should be more directly assessed using *e.g.* simulation datasets (see below). Because the above findings are contingent on a model where deleterious mutations are recessive, we also analyzed the total number of derived alleles per individual. We find a consistent but less pronounced increase in total number of derived alleles per individual for nearly neutral and mildly deleterious mutations (Figure 4B). In contrast, the total number of derived alleles for highly deleterious mutations did not correlate with the strength of bottleneck and was lowest in the most severely bottlenecked Alpi Marittime population (Figure 4B), suggesting that the most deleterious mutations were purged in this population. The *Rxy* statistics showed a corresponding strong deficit in the Alpi Marittime population (Figure 3E). We repeated the *Rxy* analyses using four additional mutation scoring methods (*i.e.* SIFT, REVEL, CADD and VEST3) and found a consistent deficit of the highest impact category in the Alpi Marittime (Figure S12). Overall, we find evidence for more purging in the most bottlenecked Alpine ibex population.

**Figure 4:**
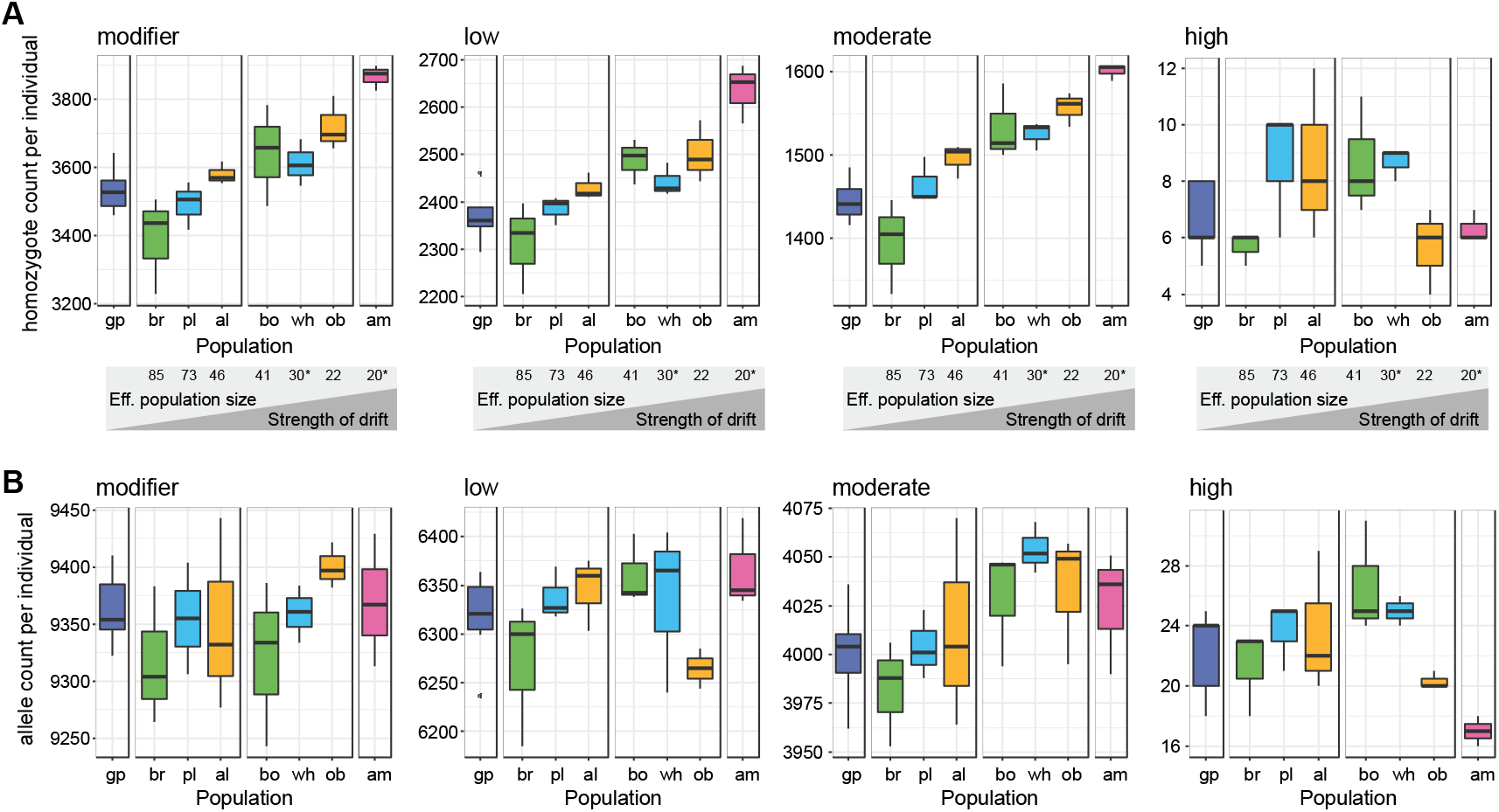
Impact of the recolonization on the individual mutation load. (A) Homozygote counts and (B) allele counts per individual for each Alpine ibex population. The schematic between A and B indicates the harmonic mean of the census size of each population, which is inversely correlated with the strength of drift. ^*)^ Estimated numbers. Colors associate founder and descendant populations (see also Figure 3A).

We analyzed the impact of highly deleterious mutations by predicting the protein truncation using homology-based inferences. Focusing on mutations segregating in Alpine ibex, we found that nearly all mutations disrupted conserved protein family (PFAM) domains encoded by the affected genes (Figure 5). This shows that highly deleterious mutations not only are altering the length of open reading frames but that evolutionarily conserved protein domains are affected by the mutations.

**Figure 5:**
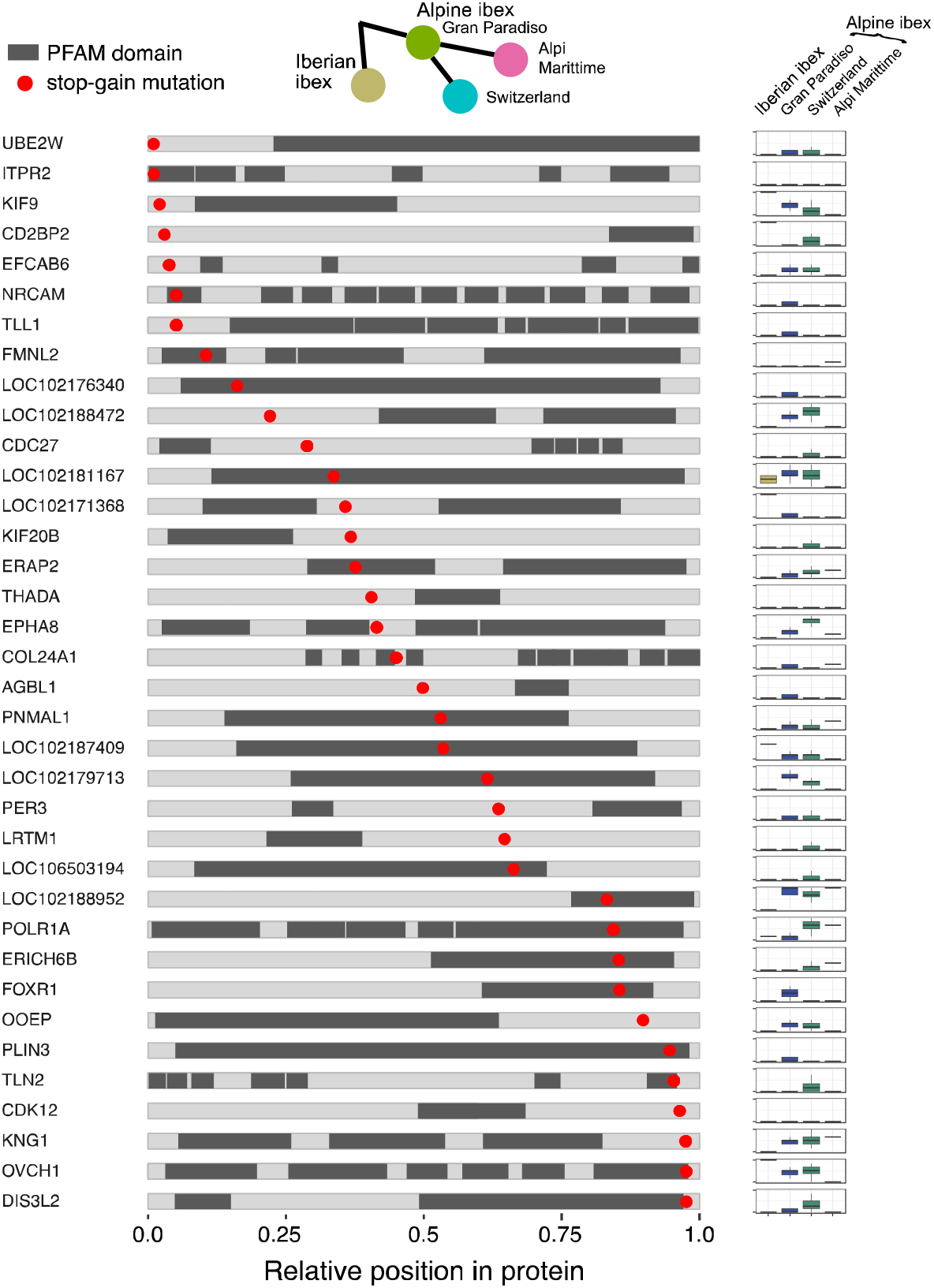
Homology-based inference of the impact of highly deleterious mutations. The localization of protein family (PFAM) domains are highlighted in dark. Red dots indicate the relative position of a highly deleterious mutation segregating in Alpine ibex. The frequencies of highly deleterious mutations are summarized for Iberian ibex and three groups of Alpine ibex. Allele frequencies are shown as boxplots per group representing the outcome of downsampling to the smallest group (Alpi Marittime, *n* = 3) based on 100 replicates.

We further analyzed the impact of bottlenecks on different mutation classes using an individual-based forward simulation model parametrized with the demographic record ^49^ (Figure 6A). The model realistically reproduced the reintroduction history and populations were parametrized with the actual founder size (Figure 6A, S16, Table S1). We used *Rxy* to analyze the evolution of deleterious mutation frequencies through the reintroduction bottlenecks. The simulations showed a deficit of highly deleterious mutations, but an increase of mildly deleterious mutations after the reintroduction bottlenecks (Figure 6B). This is consistent with our empirical evidence for purging during the species bottleneck (Figure 2E). We computed genetic load defined as the mean individual fitness in females and found an increase in Alpine ibex following the species bottleneck (Figure S17). Hence, the accumulation of mildly deleterious mutations was reducing overall fitness despite purging. The simulations also supported purging at the level of individual populations as found in the extremely bottlenecked Alpi Marittime population (Figure 3E). The number of derived mildly deleterious homozygotes increased with the strength of drift experienced by individual populations (Figure 6C, Figures S18-S21). In contrast, the median number of homozygote counts for high impact mutations were lower for Alpi Marittime than Gran Paradiso but not statistically significant (Figure 6C, Mann-Whitney U, *p* = 0.39). The highly deleterious allele counts were significantly lower for Alpi Marittime compared to Gran Paradiso (Figure S19, *p* = 0.001). We also analyzed *Rxy* for the simulation data and found that high impact mutations were indeed relatively less frequent in Alpi Marittime compared to all other Alpine ibex populations (Figure 6D) and compared to Gran Paradiso (Figure S22). Overall, the realistically parametrized model recapitulated all major empirical findings of mutation accumulation and purging across bottlenecks.

**Figure 6:**
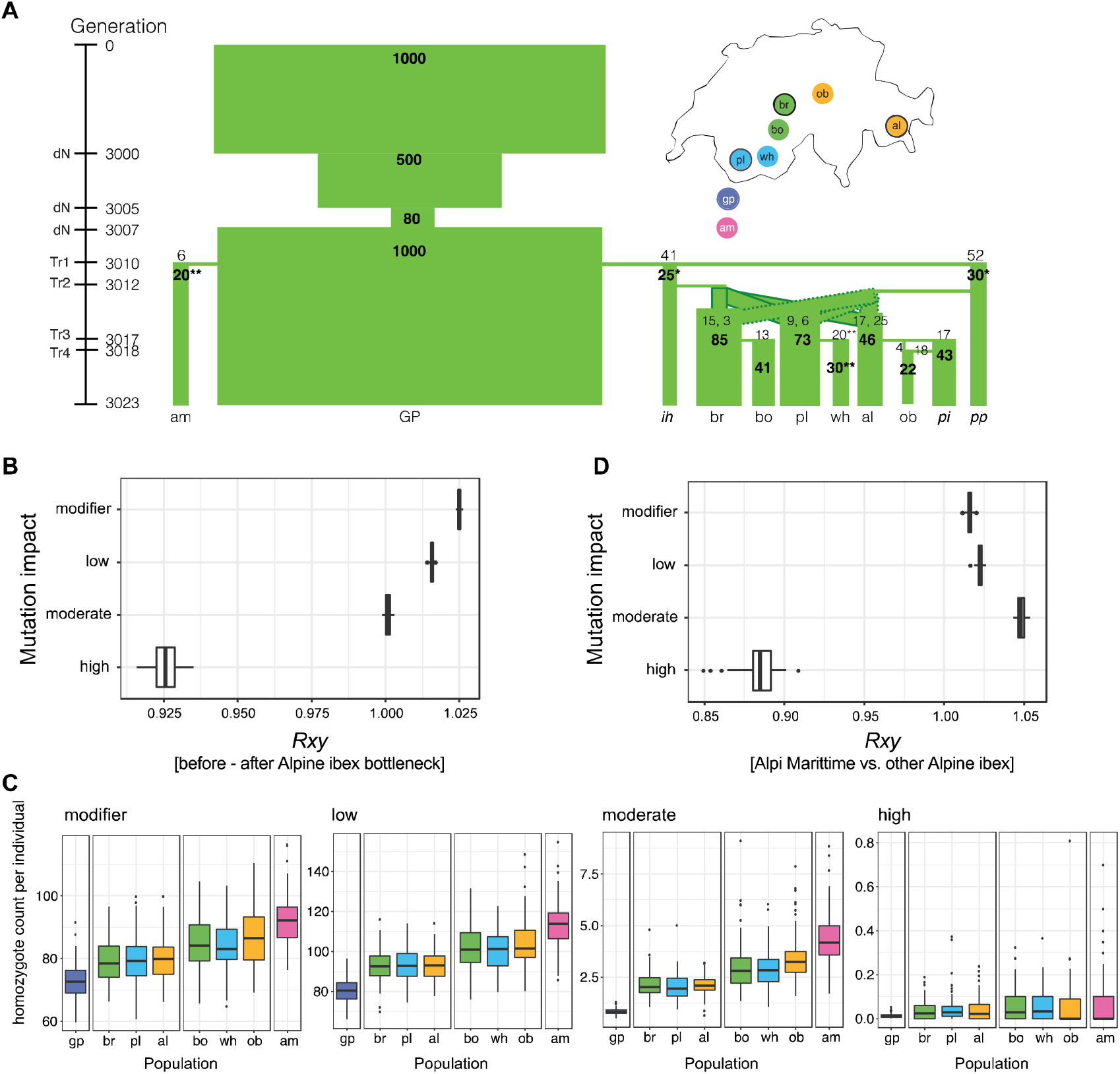
Individual-based simulations of the reintroduction history of Alpine ibex. A) The demographic model was parametrized using census data and historical records. Bold numbers show effective population sizes. Numbers not in bold indicate the number of individuals released to found each population. If a population was established from two source populations, the individual numbers are separated by commas. *) Upwards adjusted harmonic means of the census size (historical records were ih=16, Zoo Interlaken Harder, and pp = 20, Wildpark Peter and Paul). The adjustment was necessary to prevent extinction of zoo populations. **) Census numbers were estimated based on historical records of the population but no long-term data census data was available. B) Relative frequency comparison (*Rxy*) of Alpine ibex just before and after the species bottleneck and recolonization. C) Individual homozygote counts per impact category. Boxplots summarize 100 population means across simulation replicates. Colors associate founder and descendant populations (see also Figure 3A). D) *Rxy* analysis contrasting the strongly bottlenecked Alpi Marittime population with all other Alpine ibex populations across the spectrum of impact categories.

Ibex species with recently reduced population sizes accumulated deleterious mutations compared to closely related species. This accumulation was particularly pronounced in the Iberian ibex that experienced a severe bottleneck and Alpine ibex that went nearly extinct. We show that even though Alpine ibex carry an overall higher mutation burden than related species, the strong bottlenecks imposed by the reintroduction events purged highly deleterious mutations. Importantly, purging was only effective against highly deleterious mutations. Mildly deleterious mutations actually accumulated through the reintroductions. Hence, the overall number of deleterious mutations increased with bottleneck strength. This is consistent with the finding that population-level inbreeding, which is a strong indicator of past bottlenecks, is correlated with lower population growth rates in Alpine ibex ^50^.

Empirical evidence for purging in the wild is scarce ^7,20,36^. Here, we show that a few dozen generations were sufficient to reduce the burden of highly deleterious mutations. Purging may occur widely in populations undergoing severe bottlenecks contingent on populations surviving the consequences of inbreeding depression. Failure to purge under extreme bottlenecks can have severely deleterious consequences for wild populations such as shown for the Isle Royal wolves (Robinson et al. 2019). Our empirical results from the Alpine ibex reintroduction are in line with theoretical predictions that populations with an effective size below 100 individuals can accumulate a substantial burden of mildly deleterious mutations within a relatively short time resulting in increased extinction risks ^5^. The burden of deleterious mutations evident in Iberian ibex supports the notion that even population sizes of ~1000 still accumulate mildly deleterious mutations. High loads of deleterious mutations have been shown to increase the extinction risk of a species ^5^. Thus, conservation efforts aimed at keeping effective population sizes above a minimum of 1000 individuals ^2^ are critical for the long-term survival of managed species.

## Methods

### Genomic data acquisition

DNA samples from 29 Alpine ibex, 4 Iberian ibex, 2 Nubian ibex, 2 Siberian ibex and 1 Markhor individuals were sequenced on an Illumina Hiseq2500 or Hiseq4000 to a depth of 15-38 (median of 17). Table S2 specifies individual sampling locations. Libraries were produced using the TruSeq DNA Nano kit. Illumina sequencing data of 6 Bezoar and 16 domestic goat (coverage 6x – 14x, median 12x) were generated by the NextGen Consortium (https://nextgen.epfl.ch). The corresponding raw data was downloaded from the EBI Short Read Archive: ftp://ftp.sra.ebi.ac.uk/vol1/fastq/.

### Read alignment and variant calling

Trimmomatic v.0.36 ^51^ was used for quality and adapter trimming before reads were mapped to the domestic goat reference genome (version CHIR1, ^52^) using Bowtie2 v.2.2.5 ^53^. MarkDuplicates from Picard (http://broadinstitute.github.io/picard, v.1.130) was used to mark duplicates. Genotype calling was performed using HaplotypeCaller and GenotypeGVCF (GATK, v.3.6 ^54,55^). VariantFiltration of GATK was used to remove single nucleotide polymorphisms (SNP) if: QD <2.0, FS > 40.0, SOR > 5.0, MQ < 20.0, −3.0 > MQRandkSum > 3.0, −3.0 > ReadPosRankSum > 3.0 and AN < 62 (80% of all Alpine ibex individuals). Indels up to 10 bp were also retained and filtered using the same filters and filter parameters, except for not including the filter MQRankSum, because this measure is more likely to be biased for indels of several base pairs. Filtering parameters were chosen based on genome-wide quality statistics distributions (see Figures S23 – S40). Variant positions were independently validated by using the SNP caller Freebayes (v1.0.2-33-gdbb6160 ^56^) with the following settings: --no-complex --use-best-n-alleles 6 --min-base-quality 3 --min-mapping-quality 20 --no-population-priors --hwe-priors-off.

To ensure high-quality SNPs, we only retained SNPs that were called and passed filtering using GATK, and that were confirmed by Freebayes. Overall, 97.5 % of all high-quality GATK SNP calls were confirmed by Freebayes. This percentage was slightly lower for chromosome X (96.7%) and unplaced scaffolds (95.2%). We tested whether the independent SNP calls of GATK and Freebayes were concordant and we could validate 99.6% of the biallelic SNPs. We retained genotypes called by GATK and kept SNPs with a minimum genotyping rate of 90% for all further analysis. The total number of SNPs detected was 59.5 million among all species. Per species, the number of SNPs ranged from 21.9 million in the domestic goat (N=16) to 2.0 million in Markhor (N=1, Table S2).

### RNA-seq data generation

Tissue samples of a freshly harvested Alpine ibex female were immediately conserved in RNA*later* (QIAGEN) in the field and stored at −80°C until extraction. The following ten organs were sampled: retina/uvea, skin, heart, lung, lymph, bladder, ovary, kidney, liver and spleen. RNA was extracted using the AllPrep DNA/RNA Mini Kit from Qiagen following the manufacturer’s protocol. Homogenization of the samples was performed using a Retsch bead beater (Retsch GmbH) in RLT plus buffer (Qiagen). RNA was enriched using a PolyA enrichment protocol implemented in the TruSeq RNA library preparation kit. Illumina sequencing libraries were produced using the Truseq RNA stranded kit. Sequencing was performed on two lanes of an Illumina Hiseq4000.

### Genetic diversity and runs of homozygosity

Genetic diversity measured as individual number of heterozygous sites and nucleotide diversity were computed using vcftools ^57^. Runs of homozygosity were called using BCFtools/RoH ^58^, an extension of the software package BCFtools, v.1.3.1. BCFtools/RoH uses a hidden Markov model to detect segments of autozygosity from next generation sequencing data. Due to the lack of a detailed linkage map, we used physical distance as a proxy for recombination rates with the option -M and assuming 1.2cM/Mb following sheep recombination rates ^59^. Smaller values for -M led to slightly longer ROH (Figures S3 –S5). Because of small per population sample size, we decided to fix the alternative allele frequency (option --AF-dflt) to 0.4. Estimates for the population with the largest sample size (Gran Paradiso, N=7) were very similar if actual population frequencies (option --AF-estimate sp) were used (Figures S4 and S5). Option --viterbi-training was used to estimate transition probabilities before running the HMM. Running the analysis without the option --viterbi-training led to less but longer ROH (Figures S3-S5). ROH were also estimated using PLINK (v1.90b5, https://www.cog-genomics.org/plink/) with the following settings: --homozyg-window-het 2, --homozyg-window-missing 5, --homozyg-snp 100, --homozyg-kb 500, --homozyg-density 10, --homozyg-gap 100, --homozyg-window-threshold.0. ROH estimates based on PLINK were overall slightly lower but the qualitative trends hold among species and population (Figures S41 and S42).

### Identification of high-confidence deleterious mutations

Three lines of evidence were used to identify high-confidence deleterious mutations. First, variants leading to a functional change are candidates for deleterious mutations. We used snpEff ^60^ v.4.3 for the functional annotation of each variant. The annotation file ref_CHIR_1.0_top_level.gff3 was downloaded from: ftp://ftp.ncbi.nlm.nih.gov/genomes/Capra_hircus/GFF and then converted to gtf using gffread. Option -V was used to discard any mRNAs with CDS having in-frame stop codons. SnpEff predicts the effects of genetic variants (e.g. stop-gain variants) and assesses the expected impact. The following categories were retrieved: high (e.g. stop-gain or frameshift variant), moderate (e.g missense variant, in-frame deletion), low (e.g. synonymous variant) and modifier (e.g. exon variant, downstream gene variant). In the case of overlapping transcripts for the same variant, we used the primary transcript for further analysis. A total of 49.0 % of all detected SNPs were located in intergenic regions, 43.2 % in introns, 6.5 % down- and upstream of genes. A total of 0.7% of variants were within CDS, of which ~60% were synonymous and ~40% were missense variants. Overall, 0.002 % were stop-gain mutations.

Protein sequences were annotated using InterProScan v.5.33 by identifying conserved protein family (PFAM) domains ^61^.

Second, we assessed the severity of a variant by its phylogenetic conservation score. A non-synonymous variant is more likely to be deleterious if it occurs in a conserved region of the genome. We used GERP conservation scores, which are calculated as the number of substitutions observed minus the number of substitutions expected from the species tree under a neutral model. We downloaded GERP scores (accessed from http://mendel.stanford.edu/SidowLab/downloads/gerp), which have been computed for the human reference genome version hg19. The alignment was based on 35 mammal species but did not include the domestic goat (see https://genome.ucsc.edu/cgi-bin/hgTrackUi?db=hg19&g=allHg19RS_BW for more information). Exclusion of the focal species domestic goat is recommended for the computation of conservation scores, as the inclusion of the reference genome may lead to biases ^62^. In order to remap the GERP scores associated to hg19 positions to the domestic goat reference genome positions, we used liftOver (hgdownload.cse.ucsc.edu, v.287) and the chain file downloaded from hgdownload-test.cse.ucsc.edu/goldenPath/capHir1.

Third, we ascertained support for gene models annotated in the domestic goat genome with expression analyses of Alpine ibex tissue samples. We included expression data from 10 organs of an Alpine ibex female (see RNA-seq data section above) to assess expression levels of each gene model. Quality filtering of the raw data was performed using Trimmomatic ^51^ v.0.36. Hisat2 ^63^ v.2.0.5 was used to map the reads of each organ to the domestic goat reference genome. The mapping was run with option --rna-strandness RF (stranded library) and supported by including a file with known splice sites (option --known-splicesite-infile). The input file was produced using the script hisat2_extract_splice_sites.py (part of hisat2 package) from the same gtf file as the one used for the snpEff analyis (see above). For each organ, featureCounts ^64^ (subread-1.5.1) was used to count reads per each exon using the following options: -s 2 (reverse stranded) –f (count reads at the exon level), –O (assign reads to all their overlapping features), –C (excluding read pairs mapping to different chromosomes or the same chromosome but on a different strand). The R package edgeR ^65^ was used to calculate FPKM (Fragments Per Kilobase Of Exon Per Million Fragments Mapped) per each gene and organ. For variant sites that were included in more than one exon, the highest FPKM value was used. We found that 16’013 out of 17’998 genes showed transcriptional activity of at least one exon (FPKM > 0.3). Overall 166’973 out of 178’504 exons showed evidence for transcription. In a total of 1928 genes, one or more exons showed no evidence for transcription. Retained SNPs were found among 118’756 exons and 17’685 genes. Overall 611’711 out of 677’578 SNPs were located in genes with evidence for transcription.

Deleterious mutations are assumed to be overwhelmingly derived mutations. We used all ibex species except Alpine and Iberian ibex as an outgroup to define the derived state. For each biallelic site, which was observed in alternative state in Alpine ibex or Iberian ibex, the alternative state was defined as derived if its frequency was zero in all other species (a total of 44’730 autosomal SNPs). For loci with more than two alleles, the derived state was defined as unknown. For comparisons among all species, we only used the following criteria to select SNPs (370’853 biallelic SNPs retained): transcriptional activity (FPKM > 0.3 in at least one organ) and GERP > −2. The minimum GERP score cut-off was set to retain only high-quality chromosomal regions following previously established practice e.g. ^24^. We further followed the following categorizations adopted for human populations to identify moderate to highly deleterious mutations e.g. ^24^: −2 to +2 is considered neutral or near neutral, 2 to 4 considered as moderate, 4 to 6 as large and >6 as extreme effects. We also required a minimal distance to the next SNP of 3bp to avoid confounding effects of potential multi-nucleotide polymorphisms (MNPs).

### Population genetic analyses

Site frequency spectra (SFS) were calculated using the R packages *plyr* and *dplyr*. SFS analyses were performed for SnpEff and GERP categories and two additional conservation scores: phyloP ^66^ and phastCons ^67^. We chose a cutoff of 1 to distinguish conserved from less conserved sites. In the case of phyloP, sites with a score above 1 were defined as conserved. For phastCons, sites with a score equal to 1 (the maximum observed value) were considered as conserved.

For individual counts of derived alleles or homozygotes, we used all biallelic sites polymorphic either in Alpine or Iberian ibex (or both) for which the derived state was known with a maximal missing rate per locus of 10%. We retained all sites matching these criteria for any downstream analyses even if a particular site was not polymorphic in any given population. The effective rate of missing data per locus was between 0.03-0.07% (Figure S43) and no correlation was found between missing rate per population and counts. We found no qualitative differences if we included only loci with a 100% genotyping rate (Figure S44).

We calculated the relative number of derived alleles *Rxy* ^29^ for the different categories of mutations. *Rxy* compares the number of derived alleles found at sites within a specific category. Following^9,29^, we used a random set of 65592 intergenic SNPs for standardization, which makes *Rxy* robust against sampling effects and population substructure. We performed the *Rxy* analysis for the four SnpEff categories as well as four additional mutation scoring methods: SIFT ^68^, REVEL ^69^, CADD ^70^ and VEST3 ^71^. Human scores mapped to hg38 chromosomal positions were retrieved using the web interface of the Variant Effect Predictor (VEP) by ensemble^72^. The scores were mapped to chromosomal positions in the domestic goat reference genome using liftOver (http://hgdownload.cse.ucsc.edu, v.287) with the chain file accessed from http://hgdownload-test.cse.ucsc.edu/goldenPath/capHir1. As we applied these scores outside of humans, we did not have pathological evidence underpinning score cut-offs for deleteriousness. We conservatively used score cut-offs proposed as best-practices by the tool developers (Revel: 0.75, CADD: 20, SIFT: 0.05) or used a very conservative percentile (99%, 0.91 VEST3).

### Individual-based simulations with Nemo

Individual-based forward simulations were run using the software Nemo ^49^ v.2.3.51. A customized version of aNEMOne ^73^ was used to prepare input files for parameter exploration. The sim.ini file for the final set of parameters run in 100 replicates is available as Supplementary File 1. All populations relevant for the founding of the populations under study were included in the model. See Figure 6A for the simulated demography, which was modeled with the actual founder numbers (assuming a sex-ratio of 1:1), while the translocations were simplified into four phases (data from ^48^, DRYAD entry doi:10.5061/dryad.274b1 and ^47^). The harmonic mean of the population census from the founding up to the final sampling year (2007) was used to define the population carrying capacity. Mating was assumed to be random and fecundity (mean number of offspring per female) set to five. The selection coefficients of 5000 biallelic loci subject to selection were drawn from a gamma distribution with a mean of 0.01 and a shape parameter of 0.3 resulting in *s* < 1% for 99.2% of all loci ^74^ (Figure S45). Based on empirical evidence, we assumed a negative relationship between *h* and *s* ^75^. We used the exponential equation *h* = exp(−51\**s*)/2 with a mean *h* set to 0.37 following ^76^. We assumed hard selection acting at the offspring level. In addition to the 5000 loci under selection, we simulated 500 neutral loci. Recombination rates among each neutral or deleterious locus was set to 0.5. This corresponds to an unlinked state. Initial allele frequencies for the burn-in were set to *μ* / *h* * *s* = 0.0014 (corresponding to the expected mean frequency at mutation-selection balance ^77^). Mutation rate *μ* was set to 5e-05 and deleterious mutations were allowed to back-mutate at a rate of 5e-07.

A burn-in of 3000 generations was run with one population (*N* = 1000) representing the entire species allowing to reach a quasi-equilibrium. *N* was reduced to *N* = 500 for five generations before a brief, two generation bottleneck of *N* = 80. At generation 3007, the population recovered to *N* = 1000 and three generations later the reintroduction was started with the founding of the two zoos Interlaken Harder (ih) and Peter and Paul (pp). The founding of new populations was modeled by migration of offspring into an empty patch.

The zoo ih (Interlaken Harder) and several populations did not survive all replicates of the simulations. Extinction rates were as follows: ih (Zoo Interlaken Harder) 84%, bo (Bire Öschinen) 3%, wh (Weisshorn) 3%, ob (Oberbauenstock) 9%, am (Alpi Marittime) 14% and pil (Pilatus) 2%. The high extinction rate of the zoo Interlaken Harder did not affect the outcome of the simulations. The extinctions were a result of the strong reduction in population size during the founding and occurred always after the founding (see also Figure S16). The extinctions of the reintroduced populations did not affect the estimates of derived allele counts but reduced sample sizes and, hence, affected the variance of estimators. Genetic load information was retrieved using the *delet* statistics option and is defined in Nemo as the mean realized fitness (L = 1 − W_mean_/W_max_, where W_max_ is the maximum number of surviving offspring per female computed from her deleterious mutations).

### Data availability

Raw whole-genome sequencing data produced for this project was deposited at the NCBI Short Read Archive under the Accession nos. SAMN10736122–SAMN10736160 (BioProject PRJNA514886). Raw RNA sequencing data produced for this project was deposited at the NCBI Short Read Archive under the Accession nos. SAMN10839218-SAMN10839227 (BioProject PRJNA517635).

## Acknowledgments

We thank the following organizations and colleagues who contributed samples to this project. Iris Biebach, the Swiss hunting authorities of the cantons of Bern, Nidwalden, Obwalden, Uri, Graubünden and Wallis; the Gran Paradiso National Park (Alice Brambilla) and the Alpi Marittime National Park (Laura Martinelli), Sebastien Regnaut and Richard Kock, Zoological Society of London, Christian Siegenthaler, Ruedi Kunz and Samer Angelone-Alasaad. We are thankful to Glauco Camenisch and Kasia Sluzek, who provided access to Alpine ibex RNAseq datasets. We thank Laurent Excoffier, Stephan Peischl, Kimberly Gilbert, Heidi Lischer, Stefan Wyder, Thomas Wicker, Alan Brelsford, Jessica Purcell, Sarah P. Otto, Andreas Wagner, Sam Yeaman and Nicolas Perrin for helpful advice and comments on previous versions of the manuscript. We are grateful for drawings by Nadine Coline of the Zoological Museum of Zürich. This work was supported by the University of Zurich through a University Research Priority Program “Evolution in Action” pilot project grant and the Swiss Federal Office for the Environment. DC and CG were supported by the Swiss National Science Foundation (grant 31003A_173265 and 31003A_182343, respectively). This study makes use of data generated by the NextGen Consortium, which was supported by grant agreement number 244356 of the European Union’s Seventh Framework Programme (FP7/2010-2014).

## References

1. Pimm, S. L. et al. The biodiversity of species and their rates of extinction, distribution, and protection. Science 344, 1246752–1246752 (2014).

2. Frankham, R., Bradshaw, C. J. A. & Brook, B. W. Genetics in conservation management: Revised recommendations for the 50/500 rules, Red List criteria and population viability analyses. Biol Conserv 170, 56–63 (2014).

3. Cruz, F., Vilà, C. & Webster, M. T. The legacy of domestication: accumulation of deleterious mutations in the dog genome. Mol Biol Evol 25, 2331–2336 (2008).

4. Renaut, S. & Rieseberg, L. H. The accumulation of deleterious mutations as a consequence of domestication and improvement in sunflowers and other *compositae* crops. Mol Biol Evol 32, 2273–2283 (2015).

5. Lynch, M., Conery, J. & Burger, R. Mutation accumulation and the extinction of small populations. Am Nat 146, 489–518 (1995).

6. Marsden, C. D. et al. Bottlenecks and selective sweeps during domestication have increased deleterious genetic variation in dogs. PNAS 113, 152–157 (2016).

7. Robinson, J. A., Brown, C., Kim, B. Y., Lohmueller, K. E. & Wayne, R. K. Purging of strongly deleterious mutations explains long-term persistence and absence of inbreeding depression in Island Foxes. Curr Biol 1–13 (2018). doi:10.1016/j.cub.2018.08.066

8. Laenen, B. et al. Demography and mating system shape the genome-wide impact of purifying selection in Arabis alpina. PNAS 115, 816–821 (2018).

9. Xue, Y. Mountain gorilla genomes reveal the impact of long-term population decline and inbreeding. Science 348, 239–242 (2015).

10. Charlesworth, B. Effective population size and patterns of molecular evolution and variation. Nat Rev Genet 10, 195–205 (2009).

11. Kimura, M. Stochastic processes and distribution of gene frequencies under natural selection. Cold Spring Harb. Symp. Quant. Biol. 20, 33–53 (1955).

12. Keller, L. F. & Waller, D. Inbreeding effects in wild populations. Trends Ecol Evol 17, 230–241 (2002).

13. Glémin, S. How are deleterious mutations purged? Drift versus nonrandom mating. Evolution 57, 2678–2687 (2003).

14. Garcia-Dorado, A. Understanding and predicting the fitness decline of shrunk populations: inbreeding, purging, mutation, and standard selection. Genetics 190, 1461–1476 (2012).

15. Kirkpatrick, M. & Jarne, P. The effects of a bottleneck on inbreeding depression and the genetic load. Am Nat 155, 154–167 (2000).

16. Crow, J. F. in Mathematical Topics in Population Genetics (ed. Kojima, K. I.) 128–177 (Springer-Verlag, 1970).

17. Robinson, J. A. et al. Genomic flatlining in the endangered island fox. Curr Biol 26, 1183–1189 (2016).

18. Bataillon, T. & Kirkpatrick, M. Inbreeding depression due to mildly deleterious mutations in finite populations: size does matter. Genet Res 75, 75–81 (2000).

19. Willi, Y., Fracassetti, M., Zoller, S. & Van Buskirk, J. Accumulation of mutational load at the edges of a species range. Mol Biol Evol 22, 140 (2018).

20. Hedrick, P. W. & Garcia-Dorado, A. Understanding inbreeding depression, purging, and genetic rescue. Trends Ecol Evol 31, 940–952 (2016).

21. Rogers, R. L. & Slatkin, M. Excess of genomic defects in a woolly mammoth on Wrangel island. PLoS Genet 13, e1006601 (2017).

22. Robinson, J. A., Brown, C., Kim, B. Y., Lohmueller, K. E. & Wayne, R. K. Purging of strongly deleterious mutations explains long-term persistence and absence of inbreeding depression in island foxes. Curr Biol 1–13 (2018). doi:10.1016/j.cub.2018.08.066

23. Simons, Y. B., Turchin, M. C., Pritchard, J. K. & Sella, G. The deleterious mutation load is insensitive to recent population history. Nat Genet 46, 220–224 (2014).

24. Henn, B. M. et al. Distance from sub-Saharan Africa predicts mutational load in diverse human genomes. PNAS 113, E440–E449 (2015).

25. Henn, B. M., Botigué, L. R., Bustamante, C. D., Clark, A. G. & Gravel, S. Estimating the mutation load in human genomes. Nature 16, 333–343 (2015).

26. Lopez, M. et al. The demographic history and mutational load of African hunter-gatherers and farmers. Nature Ecology & Evolution 2, 1–13 (2018).

27. Scott, E. M. et al. Characterization of Greater Middle Eastern genetic variation for enhanced disease gene discovery. Nat Genet 48, 1071–1076 (2016).

28. Peischl, S. et al. Relaxed selection during a recent human expansion. Genetics 208, 763–777 (2018).

29. Do, R. et al. No evidence that selection has been less effective at removing deleterious mutations in Europeans than in Africans. Nature 47, 126–131 (2015).

30. Lohmueller, K. E. The distribution of deleterious genetic variation in human populations. Curr Opin Genet Dev 29, 139–146 (2014).

31. Fu, W., Gittelman, R. M., Bamshad, M. J. & Akey, J. M. Characteristics of Neutral and Deleterious Protein-Coding Variation among Individuals and Populations. Am J Hum Genet 95, 421–436 (2014).

32. Laws, R. J. & Jamieson, I. G. Is lack of evidence of inbreeding depression in a threatened New Zealand robin indicative of reduced genetic load? Animal Conservation 14, 47–55 (2011).

33. Kennedy, E. S., Grueber, C. E., Duncan, R. P. & Jamieson, I. G. Severe inbreeding depression and no evidence of purging in an extremely inbred wild species--the Chatham Island black robin. Evolution 68, 987–995 (2014).

34. Crnokrak, P. & Barrett, S. Perspective: Purging the genetic load: A review of the experimental evidence. Evolution 56, 2347–2358 (2002).

35. Kalinowski, S. T., Hedrick, P. W. & Miller, P. S. Inbreeding depression in the Speke’s gazelle captive breeding program. Conserv Biol 14, 1375–1384 (2000).

36. Xue, Y. et al. Mountain gorilla genomes reveal the impact of long-term population decline and inbreeding. Science 348, 242–245 (2015).

37. Grodinsky, C. & Stuwe, M. The reintroduction of the Alpine ibex to the Swiss Alps. Smithsonian 18, 68–77 (1987).

38. Biebach, I. & Keller, L. F. A strong genetic footprint of the re-introduction history of Alpine ibex (*Capra ibex ibex*). Mol Ecol 18, 5046–5058 (2009).

39. Grossen, C., Biebach, I., Angelone-Alasaad, S., Keller, L. F. & Croll, D. Population genomics analyses of European ibex species show lower diversity and higher inbreeding in reintroduced populations. Evol Appl 11, 123–139 (2018).

40. IUCN 2018. httpwww.iucnredlist.org Available at: (Accessed: 10 November 2018)

41. Couturier, M. Le Bouquetin des Alpes. (1962).

42. Reading, R. & Shank, C. Capra sibirica: The IUCN Red List of Threatened Species 2008. doi:10.2305/IUCN.UK.2008.RLTS.T42398A10695735.en

43. McQuillan, R. et al. Runs of homozygosity in European populations. The American Journal of Human Genetics 83, 359–372 (2008).

44. Cooper, G. M. et al. Distribution and intensity of constraint in mammalian genomic sequence. Genome Res 15, 901–913 (2005).

45. Terrier, G. & Rossi, P. Le bouquetin (*Capra ibex ibex*) dans les alpes maritimes franco-italiennes: occupation de l’éspace, clonisation et régulation naturelles. Traveaux Scientifiques du Parc National de la Vanoise XVIII, 271–288 (1994).

46. Maudet, C. et al. Microsatellite DNA and recent statistical methods in wildlife conservation management: applications in Alpine ibex *Capra ibex (ibex)*. Mol Ecol 11, 421–436 (2002).

47. Biebach, I. & Keller, L. F. Inbreeding in reintroduced populations: the effects of early reintroduction history and contemporary processes. Conserv Genet 11, 527–538 (2010).

48. Aeschbacher, S., Futschik, A. & Beaumont, M. A. Approximate Bayesian computation for modular inference problems with many parameters: the example of migration rates. Mol Ecol 22, 987–1002 (2013).

49. Guillaume, F. & Rougemont, J. Nemo: an evolutionary and population genetics programming framework. Bioinformatics 22, 2556–2557 (2006).

50. Bozzuto, C., Biebach, I., Muff, S., Ives, A. R. & Keller, L. F. Inbreeding reduces long-term growth of Alpine ibex populations. Nature Ecology & Evolution in press

51. Bolger, A. M., Lohse, M. & Usadel, B. Trimmomatic: a flexible trimmer for Illumina sequence data. Bioinformatics 30, 2114–2120 (2014).

52. Dong, Y. et al. Sequencing and automated whole-genome optical mapping of the genome of a domestic goat (*Capra hircus*). Nat Biotechnol 31, 135–141 (2012).

53. Langmead, B. & Salzberg, S. L. Fast gapped-read alignment with Bowtie 2. Nature Methods 9, 357–359 (2012).

54. McKenna, A. et al. The Genome Analysis Toolkit: a MapReduce framework for analyzing next-generation DNA sequencing data. Genome Res 20, 1297–1303 (2010).

55. DePristo, M. A. et al. A framework for variation discovery and genotyping using next-generation DNA sequencing data. Nat Genet 43, 491–498 (2011).

56. Garrison, E. & Marth, G. Garrison: Haplotype-based variant detection from short-read sequencing. arXiv 1207.3907 (2012)

57. Danecek, P. et al. The variant call format and VCFtools. Bioinformatics 27, 2156–2158 (2011).

58. Narasimhan, V. et al. BCFtools/RoH: a hidden Markov model approach for detecting autozygosity from next-generation sequencing data. Bioinformatics 32, 1749–1751 (2016).

59. Dumont, B. L. Payseur, Bret A. Evolution of the genomic rate of recombination in mammals. Evolution 62, 276–294 (2008).

60. Cingolani, P. et al. A program for annotating and predicting the effects of single nucleotide polymorphisms, SnpEff: SNPs in the genome of Drosophila melanogaster strain w(1118); iso-2; iso-3. fly 6, 80–92 (2012).

61. Jones, P. et al. InterProScan 5: genome-scale protein function classification. Bioinformatics 30, 1236–1240 (2014).

62. Fu, W., Gittelman, R. M., Bamshad, M. J. & Akey, J. M. Characteristics of neutral and deleterious protein-coding variationamong individuals and populations. Am J Hum Genet 95, 421–436 (2014).

63. Kim, D., Langmead, B. & Salzberg, S. L. HISAT: a fast spliced aligner with low memory requirements. Nature Methods 12, 357–360 (2015).

64. Liao, Y., Smyth, G. K. & Shi, W. featureCounts: an efficient general purpose program for assigning sequence reads to genomic features. Bioinformatics 30, 923–930 (2014).

65. Robinson, M. D., McCarthy, D. J., Bioinformatics, G. S. 2010. edgeR: a Bioconductor package for differential expression analysis of digital gene expression data.

66. Pollard, K. S., Hubisz, M. J., Rosenbloom, K. R. & Siepel, A. Detection of nonneutral substitution rates on mammalian phylogenies. Genome Res 20, 110–121 (2010).

67. Siepel, A. et al. Evolutionarily conserved elements in vertebrate, insect, worm, and yeast genomes. Genome Res 15, 1034–1050 (2005).

68. Ng, P. C. SIFT: predicting amino acid changes that affect protein function. Nucleic Acids Research 31, 3812–3814 (2003).

69. Ioannidis, N. M. et al. REVEL: An Ensemble Method for Predicting the Pathogenicity of Rare Missense Variants. Am J Hum Genet 99, 877–885 (2016).

70. Kircher, M. et al. A general framework for estimating the relative pathogenicity of human genetic variants. Nat Genet 46, 310–315 (2014).

71. Carter, H., Douville, C., Stenson, P. D., Cooper, D. N. & Karchin, R. Identifying Mendelian disease genes with the Variant Effect Scoring Tool. Bmc Genomics 14, S3 (2013).

72. McLaren, W. et al. Deriving the consequences of genomic variants with the Ensembl API and SNP Effect Predictor. Bioinformatics 26, 2069–2070 (2010).

73. kjgilbert/aNEMOne. (2017).

74. Keightley, P. D. The distribution of mutation effects on viability in Drosophila melanogaster. Genetics 138, 1315–1322 (1994).

75. Agrawal, A. F. & Whitlock, M. C. Inferences about the distribution of dominance drawn from yeast gene knockout data. Genetics 187, 553–566 (2011).

76. Gilbert, K. J. et al. Local Adaptation Interacts with Expansion Load during Range Expansion: Maladaptation Reduces Expansion Load. Am Nat 189, 368–380 (2017).

77. Crow, J. F. & Kimura, M. An Introduction to Population Genetics Theory. (New Jersey: Blackburn Press, 1970).

