## Supplementary Information for "Purging of highly deleterious mutations through severe bottlenecks in Alpine ibex"

### SUPPLEMENTARY FIGURES

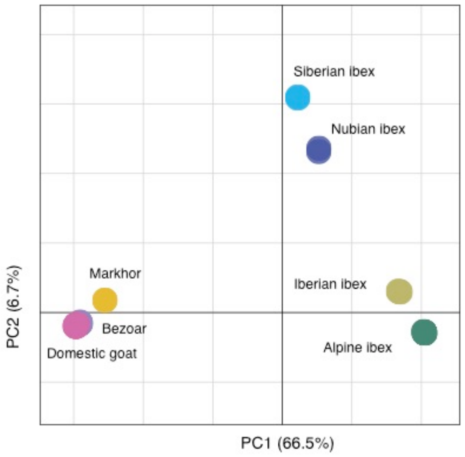

**Figure S1:** Principal component analysis including all ibex species under study. Colors indicate different species.

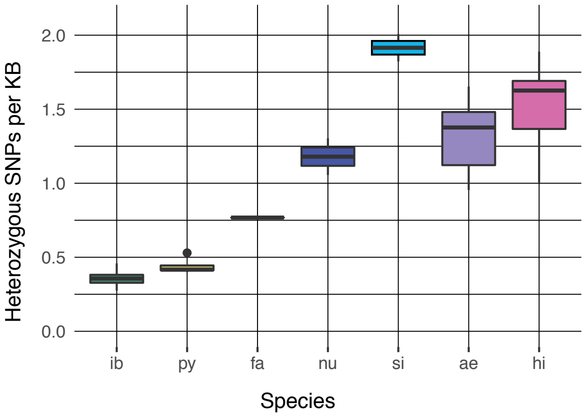

**Figure S2A:** Genome-wide heterozygosity per species represented as number of heterozygous autosomal SNPs per kb. ib: *Capra ibex*, py: *C. pyrenaica*, fa: *C. falconeri*, nu: *C. nubiana*, si: *C. sibirica*, ae: *C. aegagrus*, hi: *C. hircus*.

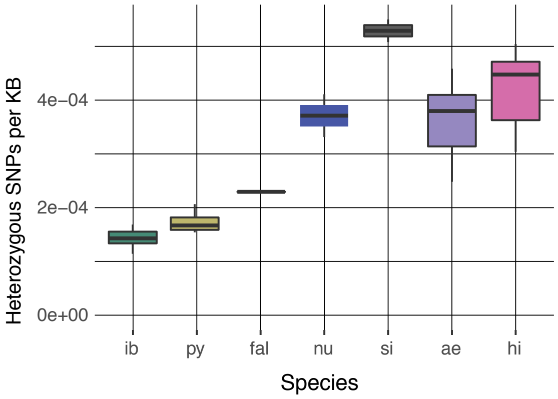

**Figure S2B:** Genome-wide heterozygosity per species in coding sequences (CDS) represented as number of heterozygous SNPs per kb of CDS. ib: *Capra ibex*, py: *C. pyrenaica*, fa: *C. falconeri*, nu: *C. nubiana*, si: *C. sibirica*, ae: *C. aegagrus*, hi: *C. hircus*.

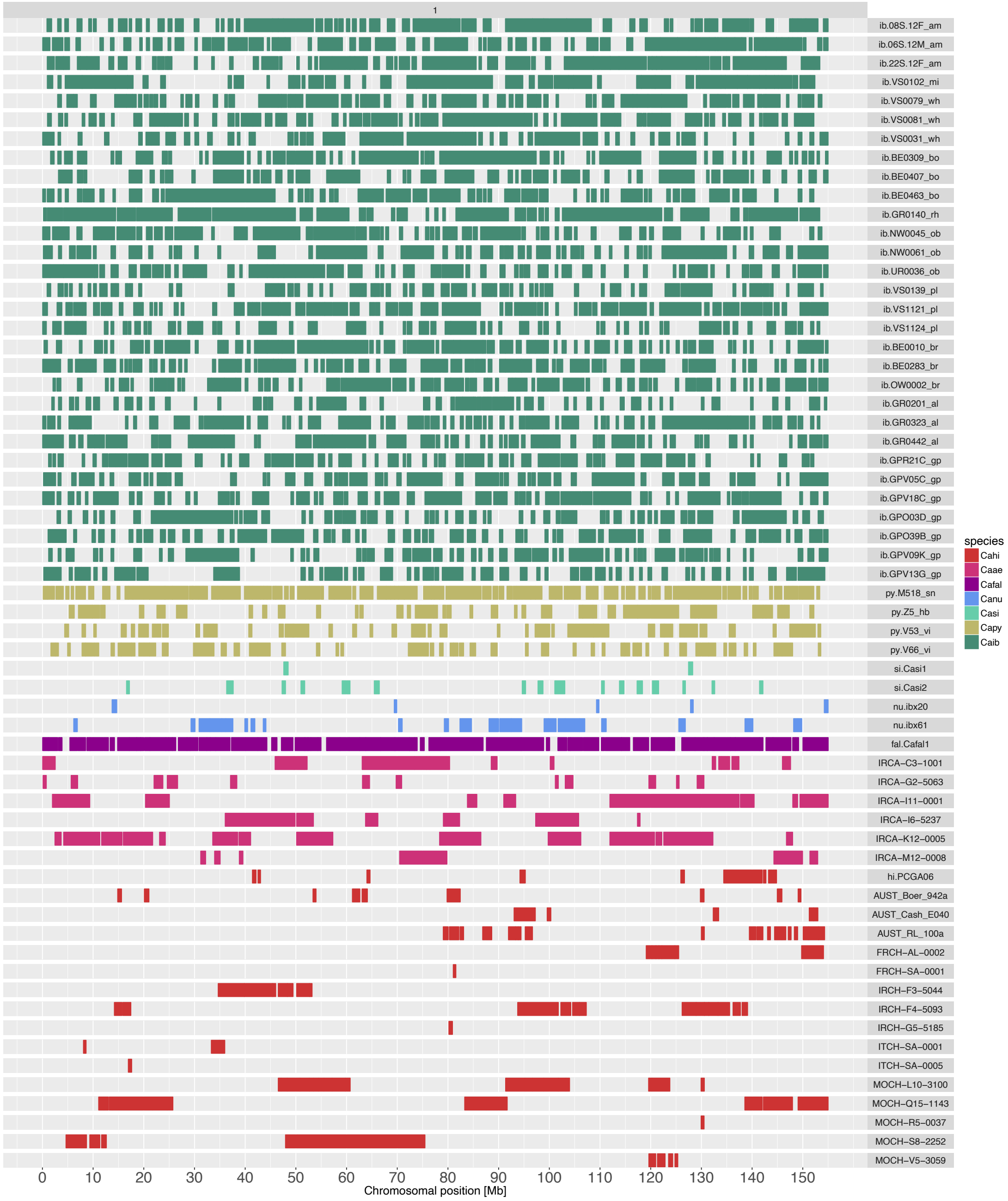

**Figure S3A:** Runs of homozygosity (ROH) along chromosome 1. Bcftools analysis with viterbi-training option turned off. The alternative allele frequency prior was set to 0.4. Colors represent species as in Figures S1 and S2. Caib: *Capra ibex*, Capy: *C. pyrenaica*, Cafal: *C. falconeri*, Canu: *C. nubiana*, Casi: *C. sibirica*, Caae: *C. aegagrus*, Cahi: *C. hircus*.

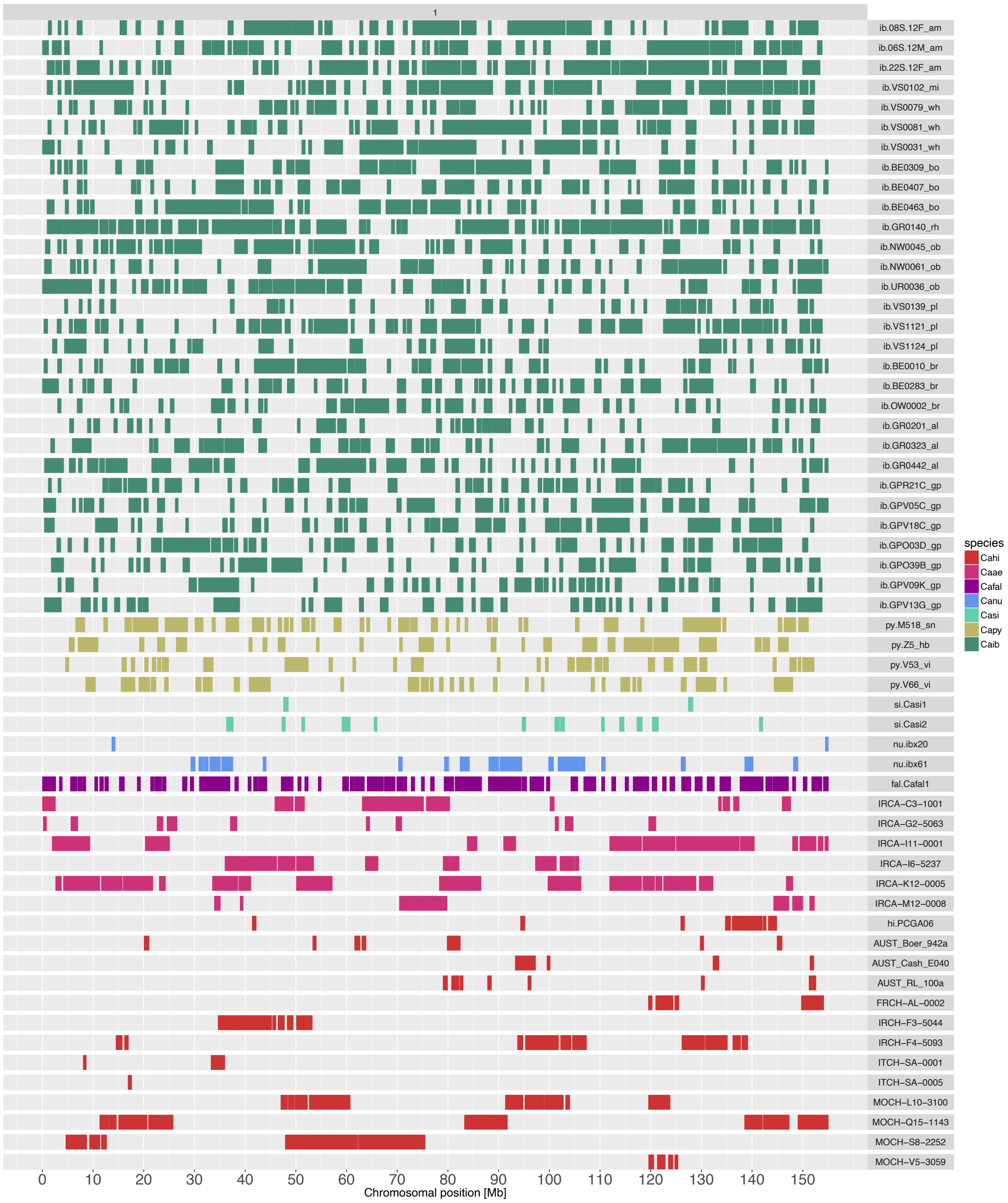

**Figure S3B:** Runs of homozygosity (ROH) along chromosome 1. Bcftools was run with the viterbi-training option turned on. The alternative allele frequency prior was set to 0.4. Colors represent species as in Figures S1 and S2. Caib: *Capra ibex*, Capy: *C. pyrenaica*, Cafal: *C. falconeri*, Canu: *C. nubiana*, Casi: *C. sibirica*, Caae: *C. aegagrus*, Cahi: *C. hircus*.

**Figures S4A - S4D:** Runs of homozygosity (ROH) analysis with different bcftools settings. Red lines show estimated state (0 corresponds to ROH region, 1 to non-ROH region). The blue line shows heterozygosity in a sliding window (window size=100 bp, step size=1 bp) along chromosomal positions. Data is shown for the *C. ibex* individual GPR21C. The window represents chromosome 23 and positions 15-30 Mb.

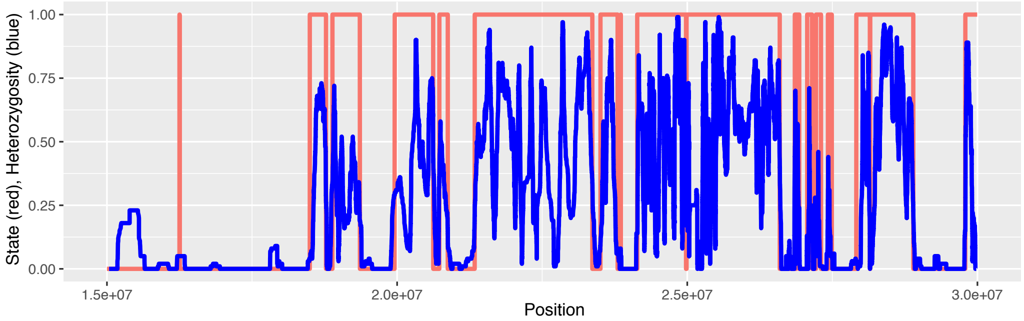

**Figure S4A:** Allele frequency prior (af.dtlt) set to 0.4.

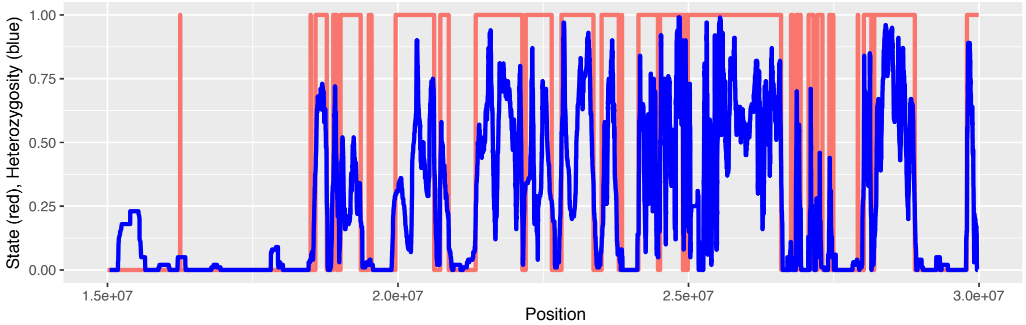

**Figure S4B:** Allele frequency prior (af.dtlt) set to 0.4 and viterbi-training option turned on.

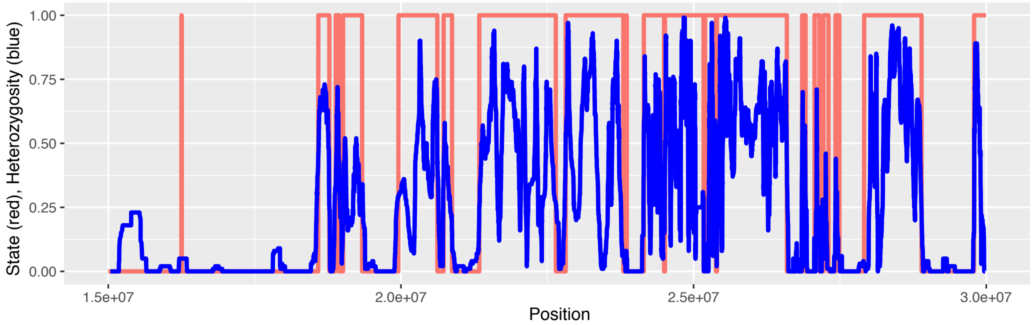

**Figure S4C:** Allele frequency priors were set to allele frequencies observed in the Gran Paradiso population.

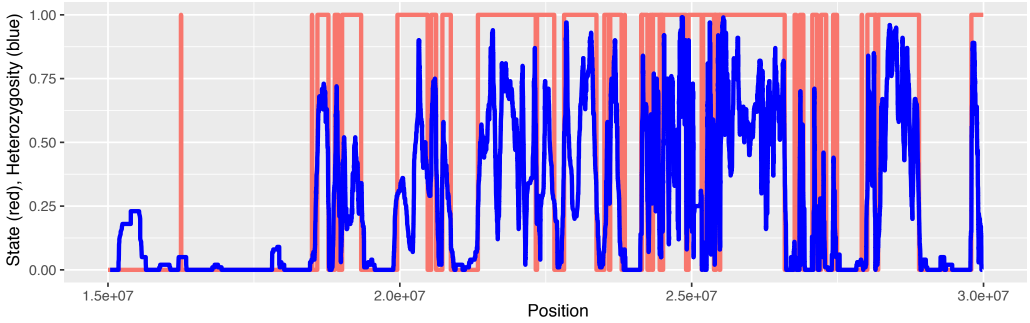

**Figure S4D:** Allele frequency priors were set to allele frequencies observed in the Gran Paradiso population and viterbi-training option turned on.

**Figures S5A - S5D:** Runs of homozygosity (ROH) analysis with different bcftools settings. Red lines show estimated state (0 corresponds to ROH region, 1 to non-ROH region). The blue line shows heterozygosity in a sliding window (window size=100 bp, step size=1 bp) along chromosomal positions. Data is shown for the *C. ibex* individual GPV05C. The window represents chromosome 23 and positions 15-30 Mb.

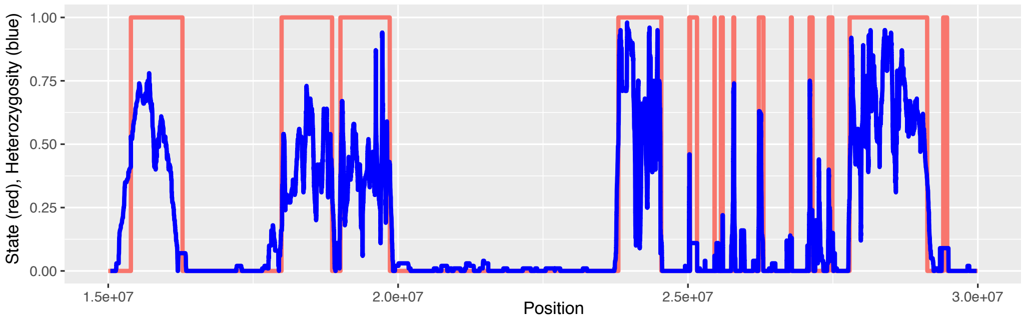

**Figure S5A:** Allele frequency prior (af.dtlt) set to 0.4.

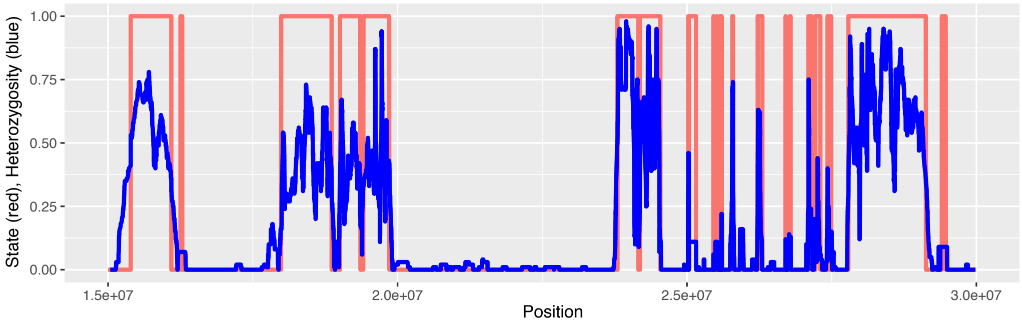

**Figure S5B:** Allele frequency prior (af.dtlt) set to 0.4 and viterbi-training option turned on.

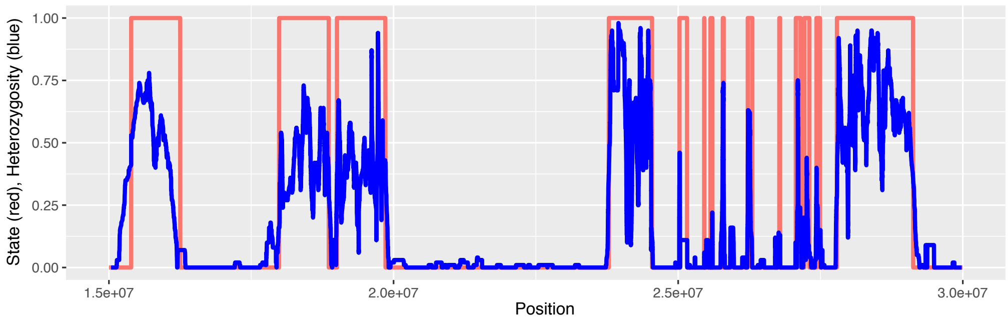

**Figure S5C:** Allele frequency priors were set to allele frequencies observed in the Gran Paradiso population.

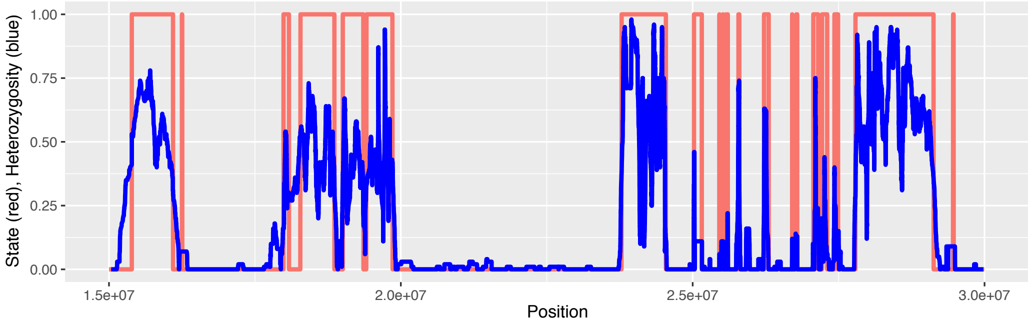

**Figure S5D:** Allele frequency priors were set to allele frequencies observed in the Gran Paradiso population and viterbi-training option turned on.

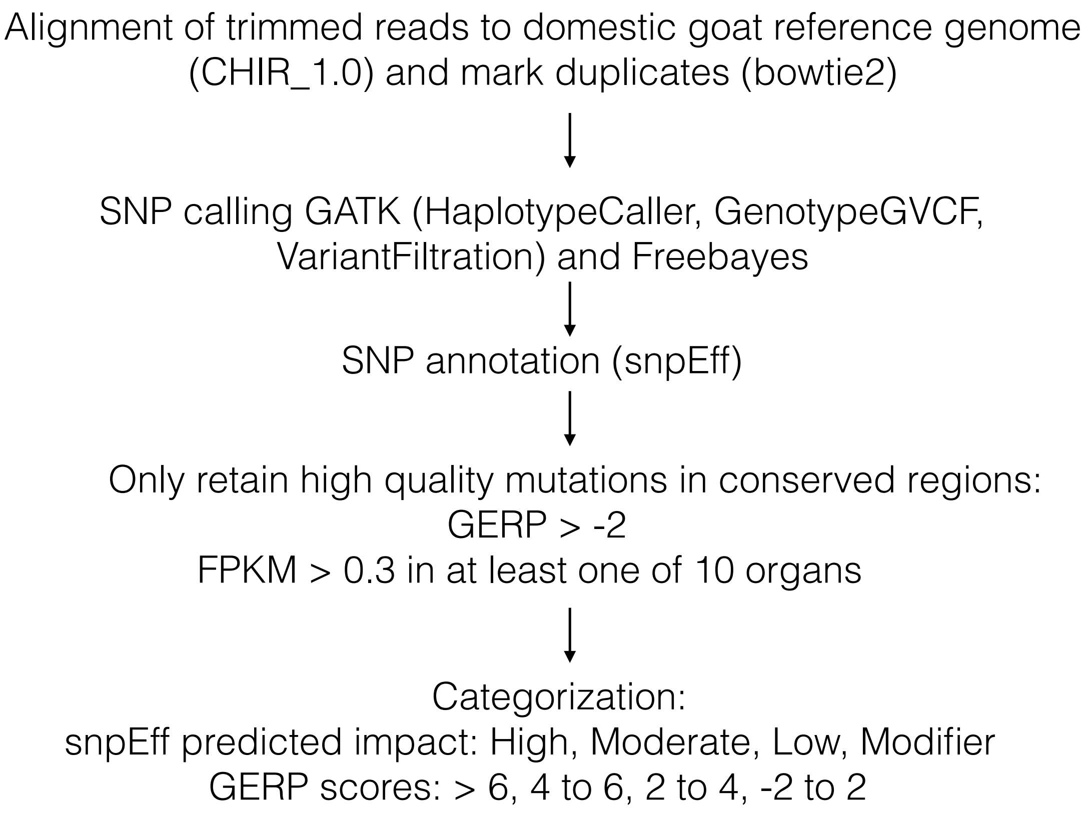

**Figure S6:** Pipeline for SNP calling, validation and categorization of mutational impact. See Methods for details on each step.

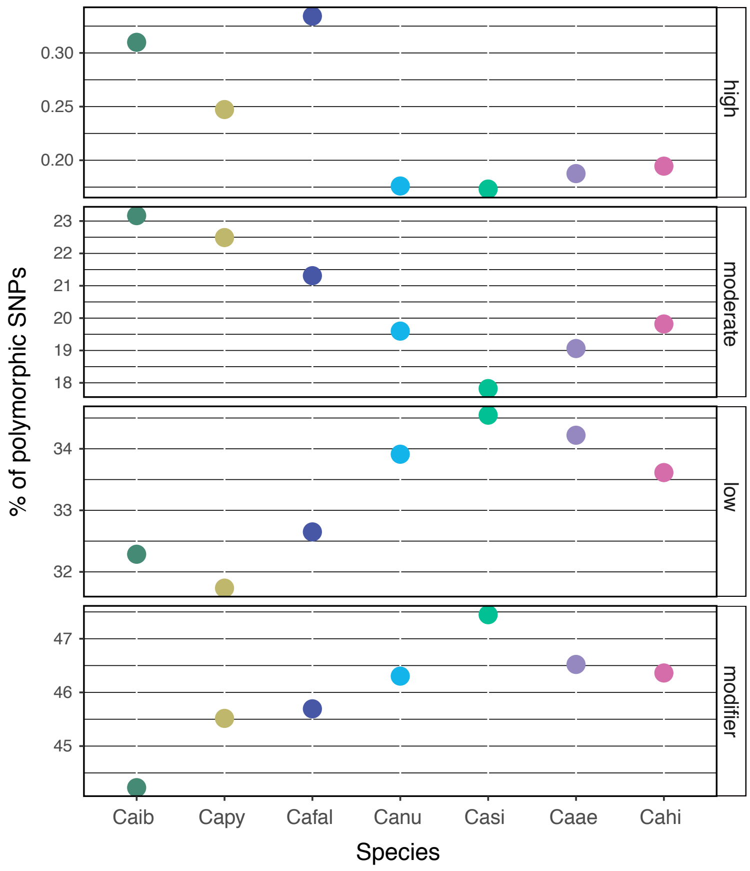

**Figure S7A:** Proportion of single nucleotide polymorphisms (SNPs) polymorphic within species categorized into four impact categories according to SnpEff (high, moderate, low, modifier). Caib: *Capra ibex*, Capy: *C. pyrenaica*, Cafal: *C. falconeri*, Canu: *C. nubiana*, Casi: *C. sibirica*, Caae: *C. aegagrus*, Cahi: *C. hircus*.

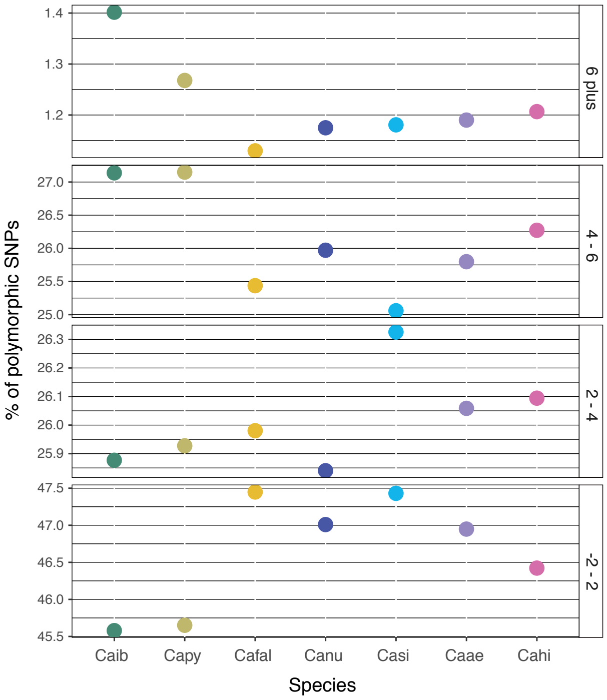

**Figure S7B:** Proportion of single nucleotide polymorphisms (SNPs) polymorphic within species categorized into GERP score ranges (>6, 4 - 6, 2 - 4, -2 - 2). Caib: *Capra ibex*, Capy: *C. pyrenaica*, Cafal: *C. falconeri*, Canu: *C. nubiana*, Casi: *C. sibirica*, Caae: *C. aegagrus*, Cahi: *C. hircus*.

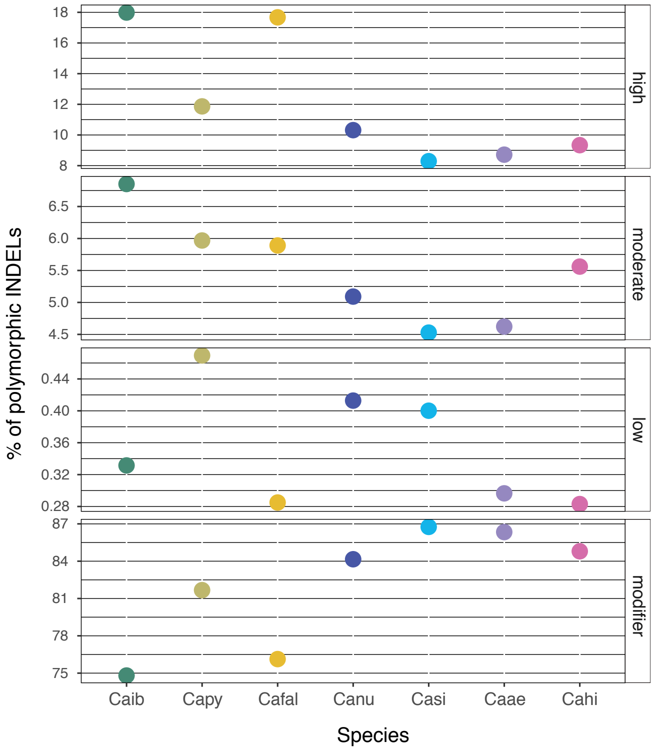

**Figure S7C:** Proportion of indels (≤10 bp) polymorphic within species categorized into four impact categories according to SnpEff (high, moderate, low, modifier). Caib: *Capra ibex*, Capy: *C. pyrenaica*, Cafal: *C. falconeri*, Canu: *C. nubiana*, Casi: *C. sibirica*, Caae: *C. aegagrus*, Cahi: *C. hircus*.

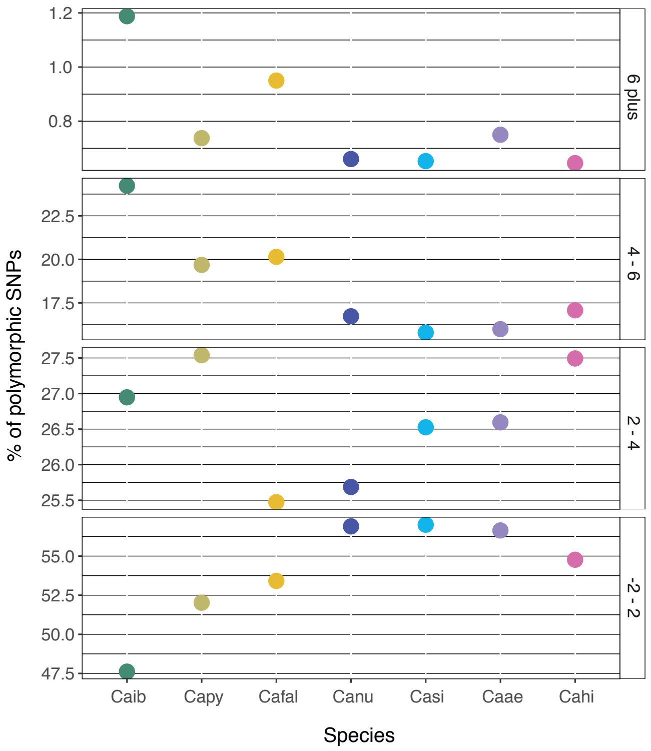

**Figure S7D:** Proportion of variant indels (up to 10 bp long) polymorphic within species categorized into GERP score ranges (>6, 4 - 6, 2 - 4, -2 - 2). Caib: *Capra ibex*, Capy: *C. pyrenaica*, Cafal: *C. falconeri*, Canu: *C. nubiana*, Casi: *C. sibirica*, Caae: *C. aegagrus*, Cahi: *C. hircus*.

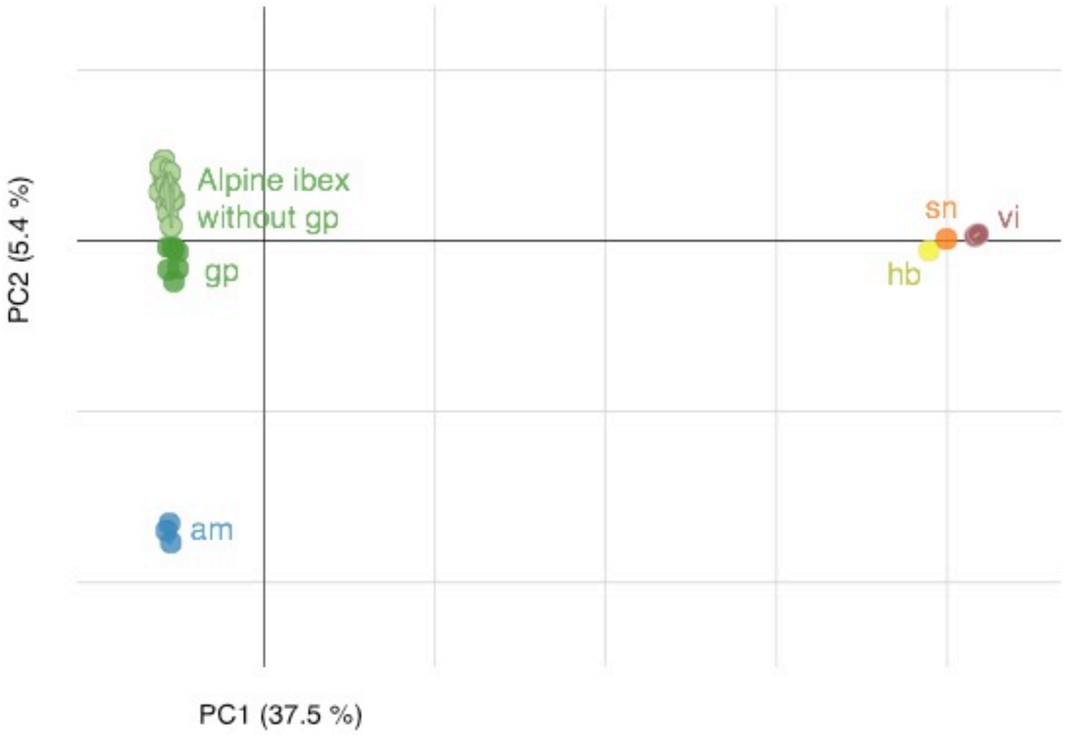

**Figure S8:** Principal component analyses of Alpine ibex (dark green: Gran Paradiso {gp}, light green: Swiss individuals, blue: Alpi Marittime {am}) and Iberian ibex (yellow: Maestrazgo {hb}, orange: Sierra Nevada {sn}, brown: Victoriae {vi}).

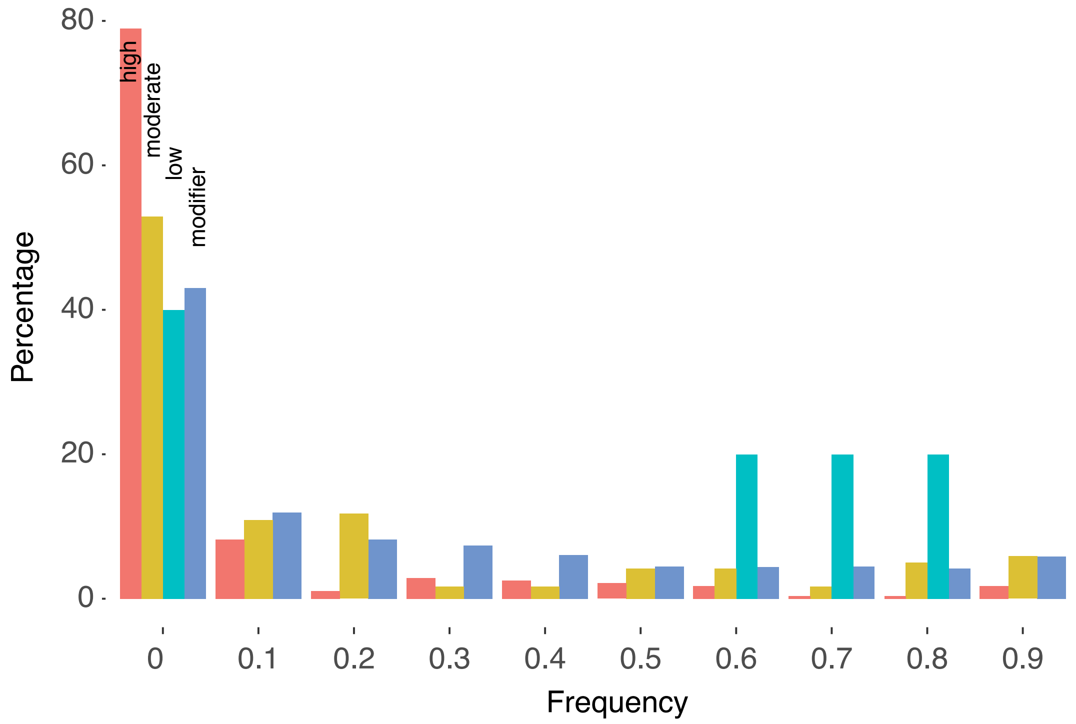

**Figure S9:** Site-frequency spectrum (SFS) of different SnpEff categories of indel mutations in Alpine ibex.

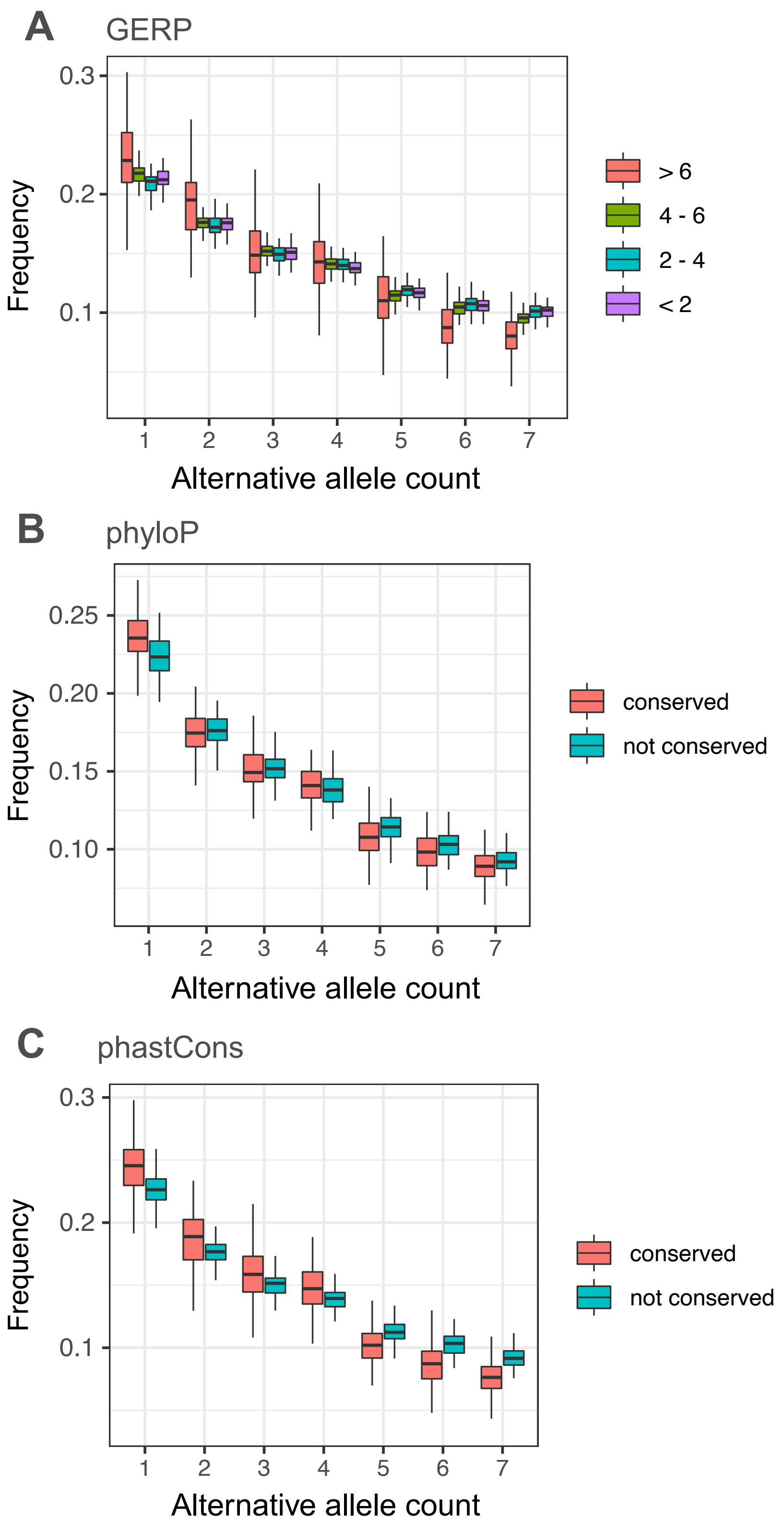

**Figure S10:** Site-frequency spectra (SFS) in Alpine ibex of different categories of SNP mutations based on A) GERP scores divided into four ranges, B) phyloP scores (conserved: > 1, not conserved: ≤ 1) and C) phastCons scores (conserved: =1, not conserved: < 1). See Figure 2 for the SFS of SNPs categorized into SnpEff categories high, moderate, low and modifier.

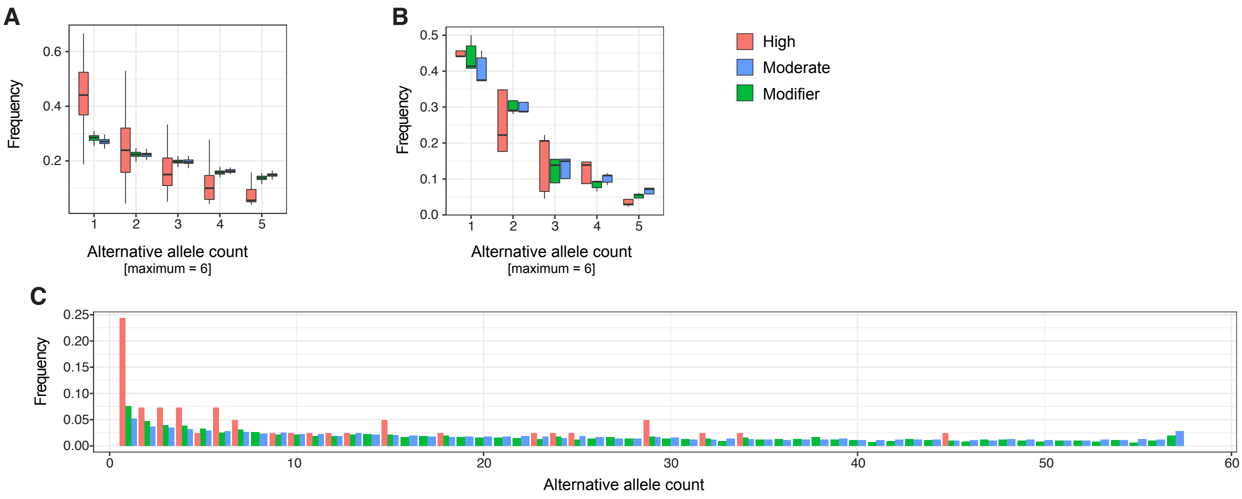

**Figure S11:** Site frequency spectra (SFS) for different categories of SNP mutations based on SnpEff (high, moderate and modifier categories). A) SFS downsampled to three Alpine ibex individuals based on jackknifing (100 replicates). B) SFS downsampled to three Iberian ibex individuals based on jackknifing (100 replicates). C) SFS for all Alpine ibex without grouping (see also Figure 2B).

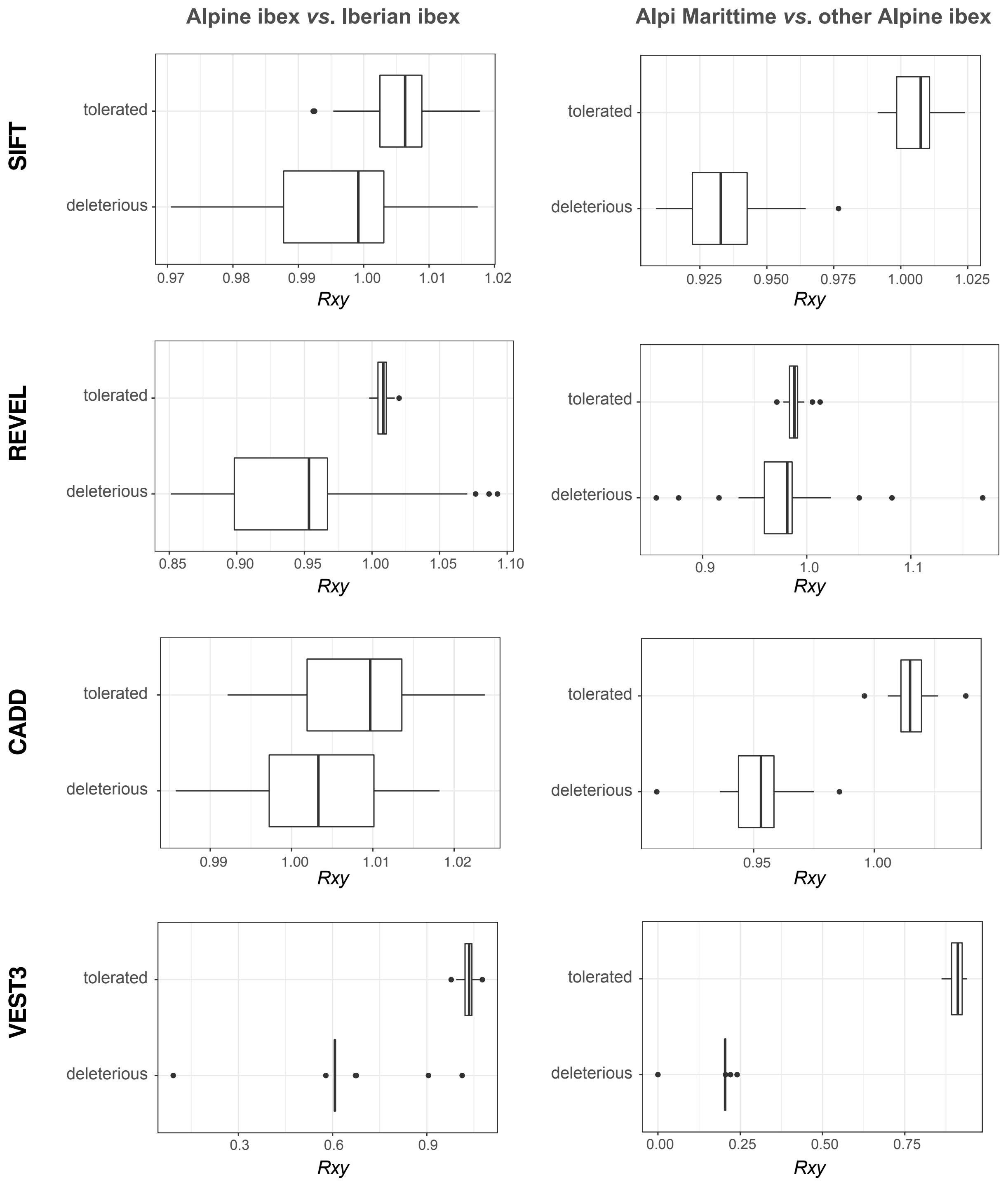

**Figure S12:** *Rxy* analysis performed based on four additional mutation scoring methods (SIFT, Revel, CADD, VEST3). *Rxy* < 1 indicates a relative frequency deficit of the corresponding category in Alpine ibex compared to Iberian ibex (left column) or Alpi Marittime compared to other Alpine ibex (right column). *Rxy* distributions are based on jack-knifing across chromosomes. Score cut-offs used to define deleteriousness were as follows: Revel: 0.75, CADD: 20, SIFT: 0.05, VEST3: 0.91 (99%-percentile). See methods for details.

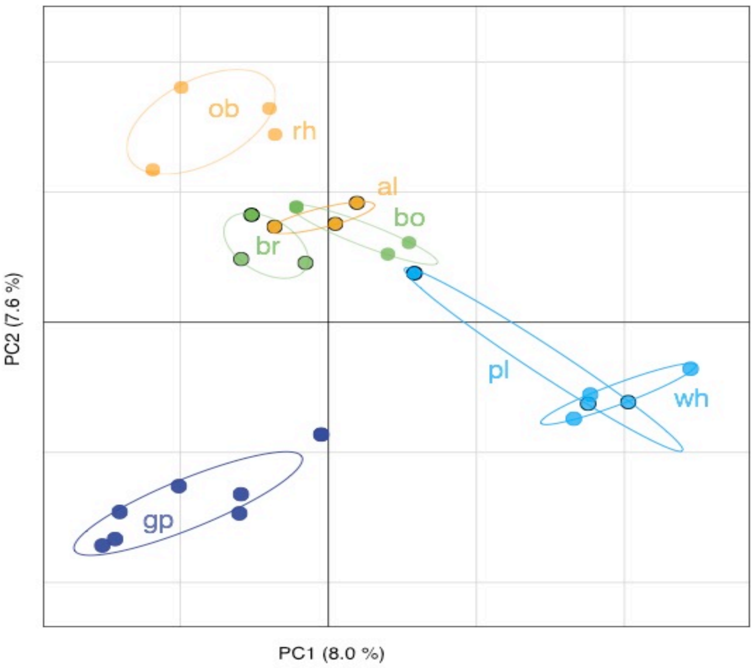

**Figure S13A:** Principal component analysis plot of all Alpine ibex individuals except the individuals from the highly divergent population Alpi Marittime. Circles with black rim indicate the three populations used for all further population reintroductions of Alpine ibex. Same colors of Alpine ibex populations join founder and descendent population. bo, Bire Öschinen; gp, Gran Paradiso; al, Albris; br, Brienzer Rothorn; ob, Oberbauenstock; pl, Pleureur; rh, Rheinwald; wh, Weisshorn.

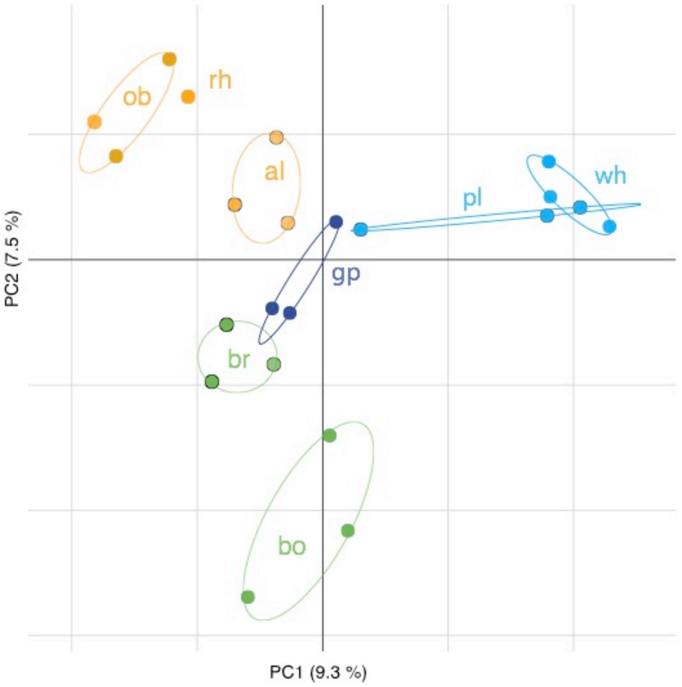

**Figure S13B:** Principal component analysis plot of all Alpine ibex individuals except the individuals from the highly divergent population Alpi Marittime. The population from Gran Paradiso was reduced to three randomly chosen individuals. bo, Bire Öschinen; gp, Gran Paradiso; al, Albris; br, Brienzer Rothorn; ob, Oberbauenstock; pl, Pleureur; rh, Rheinwald; wh, Weisshorn.

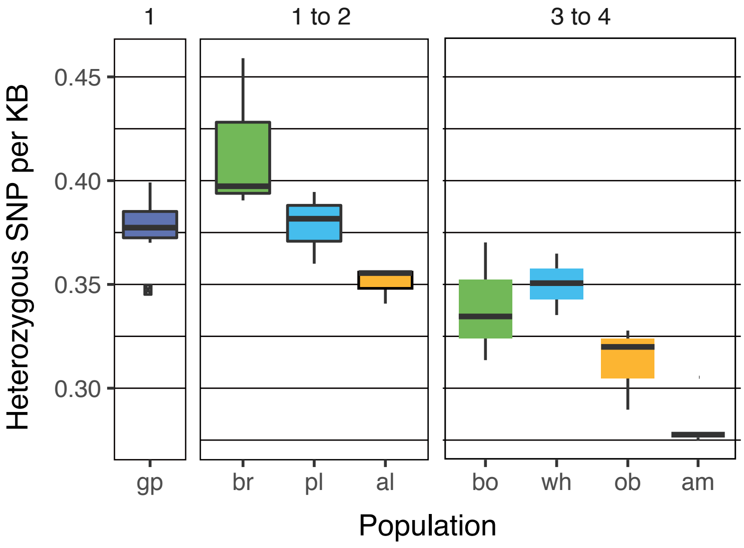

**Figure S14:** Genome-wide heterozygosity per Alpine ibex population represented as the number of heterozygous, autosomal SNPs per kb. Colors as in Figures 3 and S11. bo, Bire Öschinen; gp, Gran Paradiso; al, Albris; br, Brienzer Rothorn; ob, Oberbauenstock; pl, Pleureur; rh, Rheinwald; wh, Weisshorn.

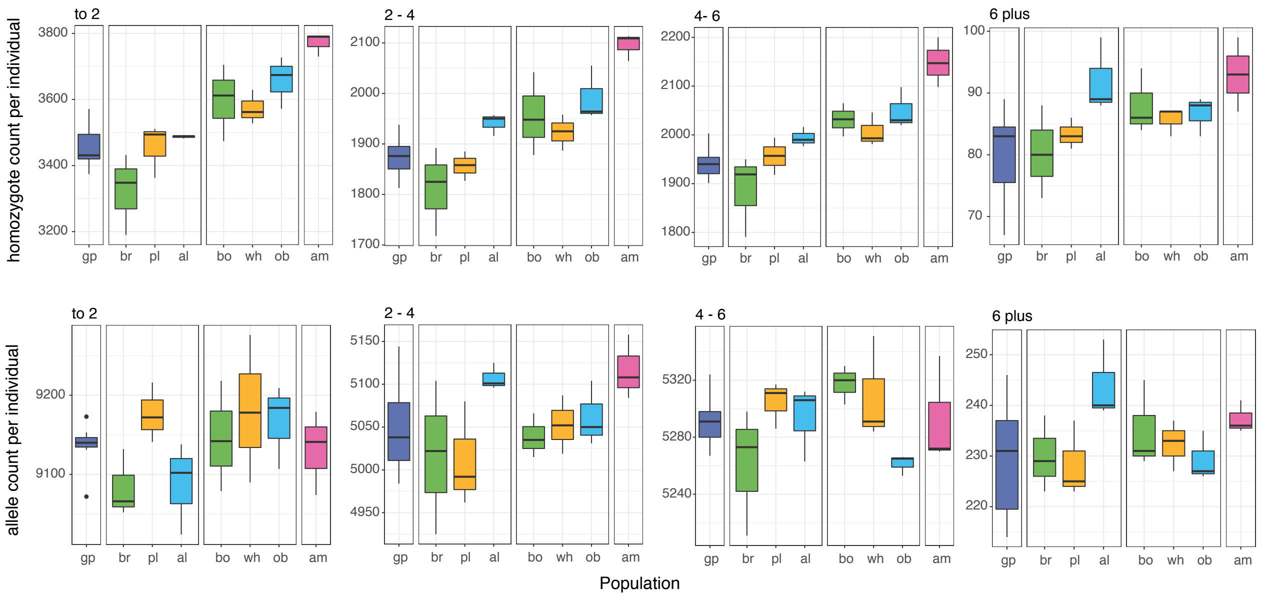

**Figure S15:** Individual homozygote counts (upper panel) and allele counts (lower panel) for each Alpine ibex population. Left to right shows GERP score ranges of -2 - 2, 2 - 4, 4 - 6 and >6 of SNP mutations. Colors group Alpine ibex populations into founder and descendent population pairs (see Figure 3).

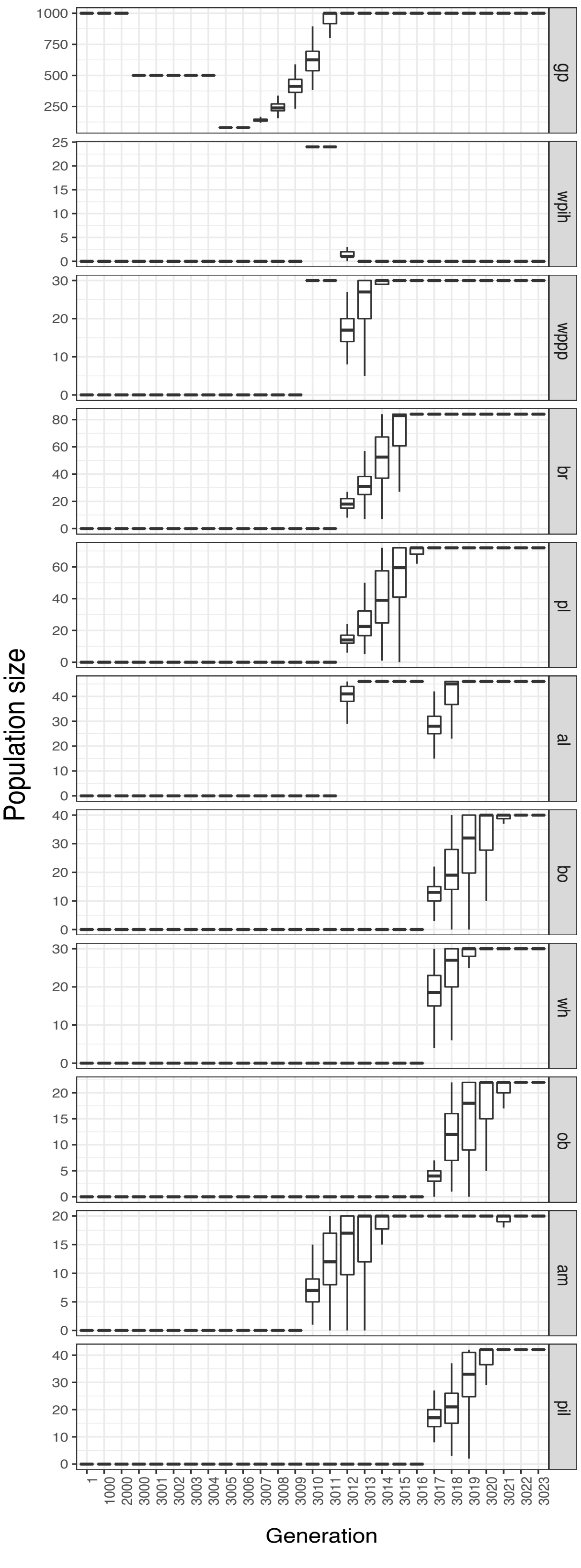

**Figure S16:** Population size changes over the course of the demographic simulations. Population size did not change between generation 1 and 2999. am: Alpi Marittime, gp: Gran Paradiso; ih: Zoo Interlaken Harder; al: Albris; bo: Bire Öschinen; br: Brienzer Rothorn; ob: Oberbauenstock; pl: Pleureur; rh: Rheinwald; wh: Weisshorn; pil: Pilatus; wpih: Wildpark Interlaken Harder; wppp: Wildpark Peter and Paul.

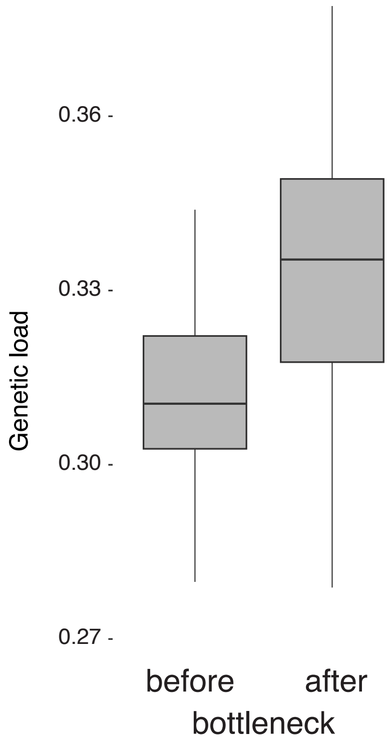

**Figure S17:** Boxplots showing genetic load before and after the simulated species bottleneck in Alpine ibex. Genetic load was defined as mean individual fitness of females.

Before species bottleneck

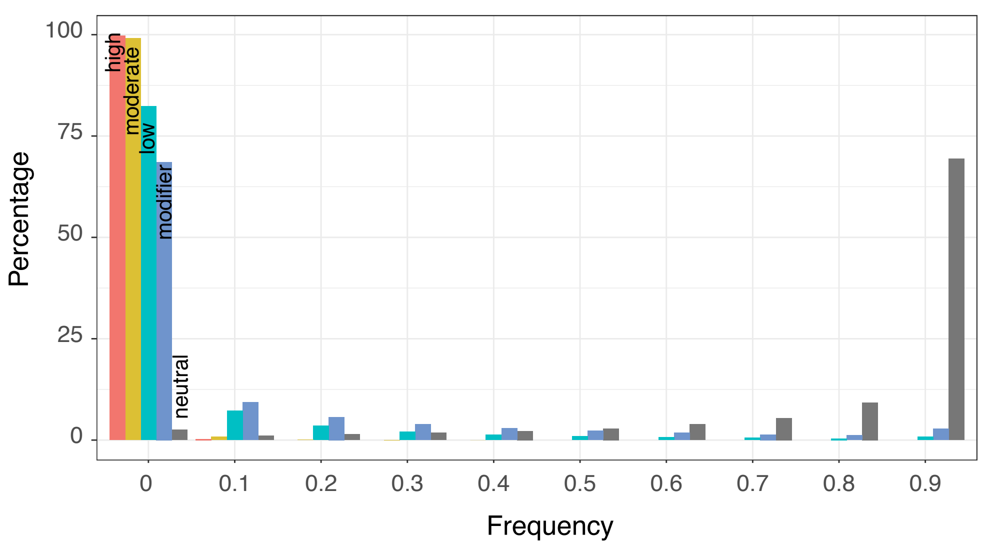

After species bottleneck

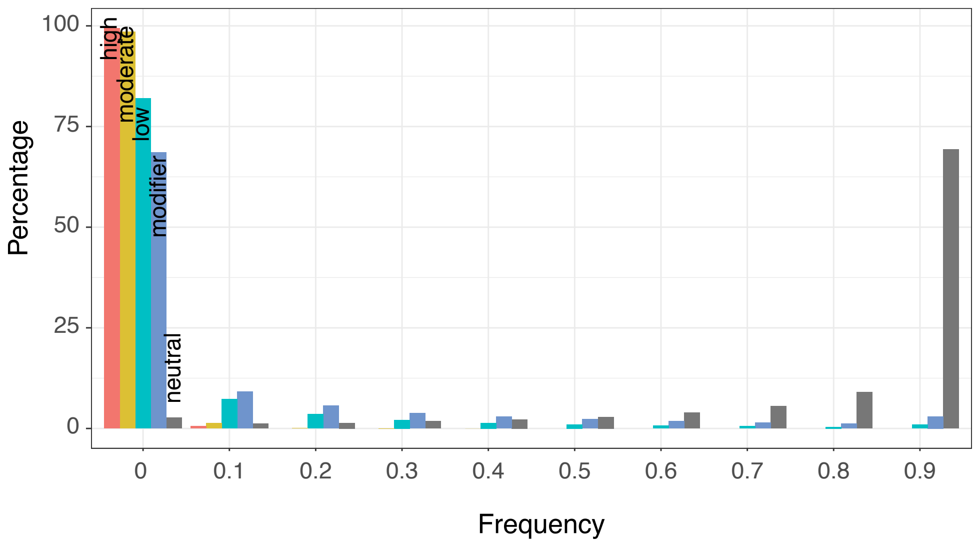

**Figure S18:** Outcome of the demographic simulations. Site-frequency spectra before and after the species bottleneck for different mutation categories (neutral, modifier, low, moderate, high). Deleterious mutation categories were defined based on the selection coefficient $s$ as follows: Modifier: $s < 0.0001$, Low: $0.0001 \leq s < 0.01$, Moderate: $0.01 \leq s < 0.1$, High: $0.1 \leq s$. The bottleneck occurred between generations 2999 and 3023.

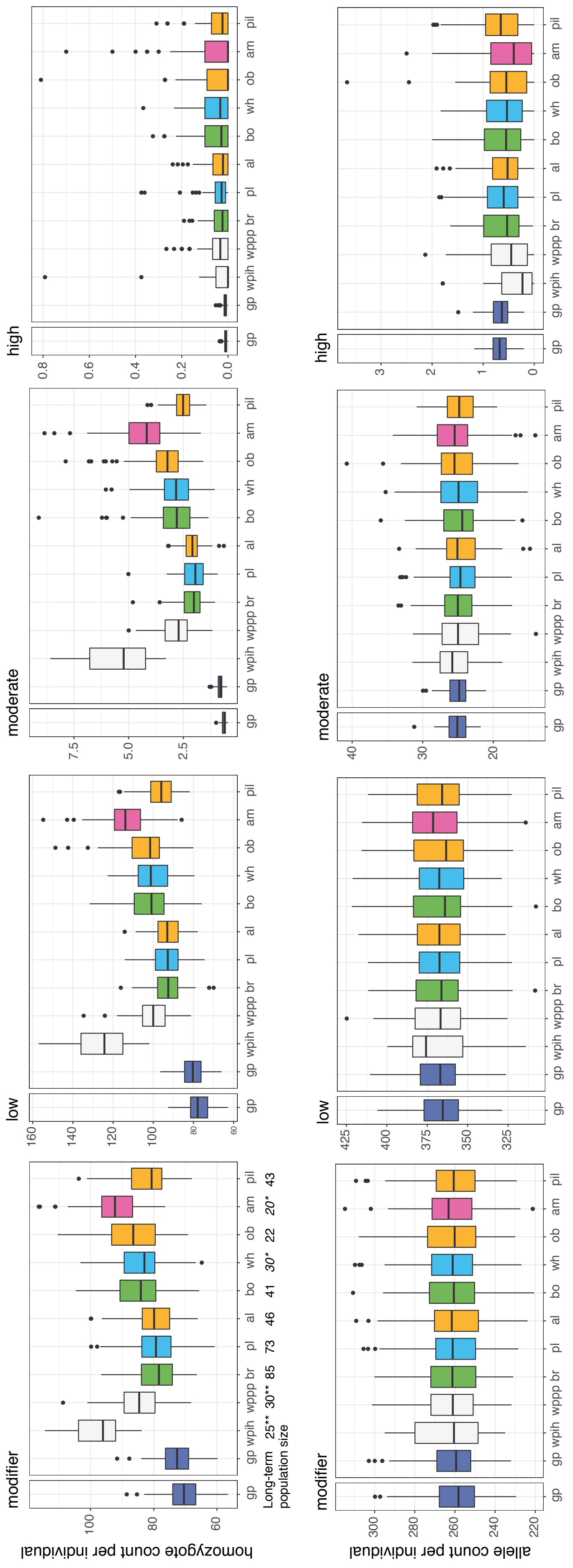

**Figure S19:** Outcome of the demographic simulations showing individual homozygote and allele counts grouped by population. Simulated populations included all major wild parks and populations that contributed to the founding of the Alpine ibex populations under study (see also Figure 3A). Matching colours group source populations with populations that were founded from them (except for wpih, wppp, gp and am). Deleterious mutation categories were defined based on the selection coefficient *s* as follows: Modifier: $s < 0.0001$, Low: $0.0001 \leq s < 0.01$, Moderate: $0.01 \leq s < 0.1$, High: $0.1 \leq s$. Abbreviations: gp: Gran Paradiso, wpih: Wildpark Interlaken Harder, wppp: Wildpark Peter and Paul, br: Brienzer Rothorn, pl: Pleureur, al: Albris, bo: Bire Oeschinen, wh: Weisshorn, ob: Oberbauenstock, am: Alpi Marittime, pil: Pilatus.

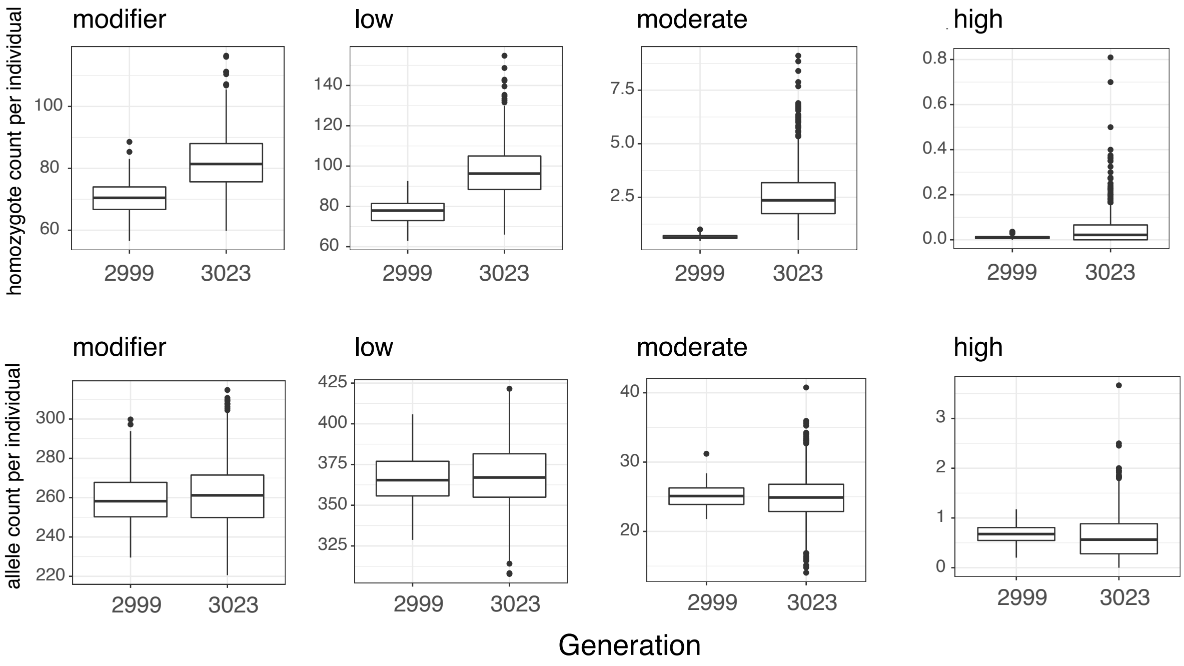

**Figure S20:** Outcome of the demographic simulations showing individual homozygote and allele counts before and after the species bottleneck (generations 2999 and 3023, respectively). Deleterious mutation categories were defined based on the selection coefficient *s* as follows: Modifier: $s < 0.0001$, Low: $0.0001 \leq s < 0.01$, Moderate: $0.01 \leq s < 0.1$, High: $0.1 \leq s$.

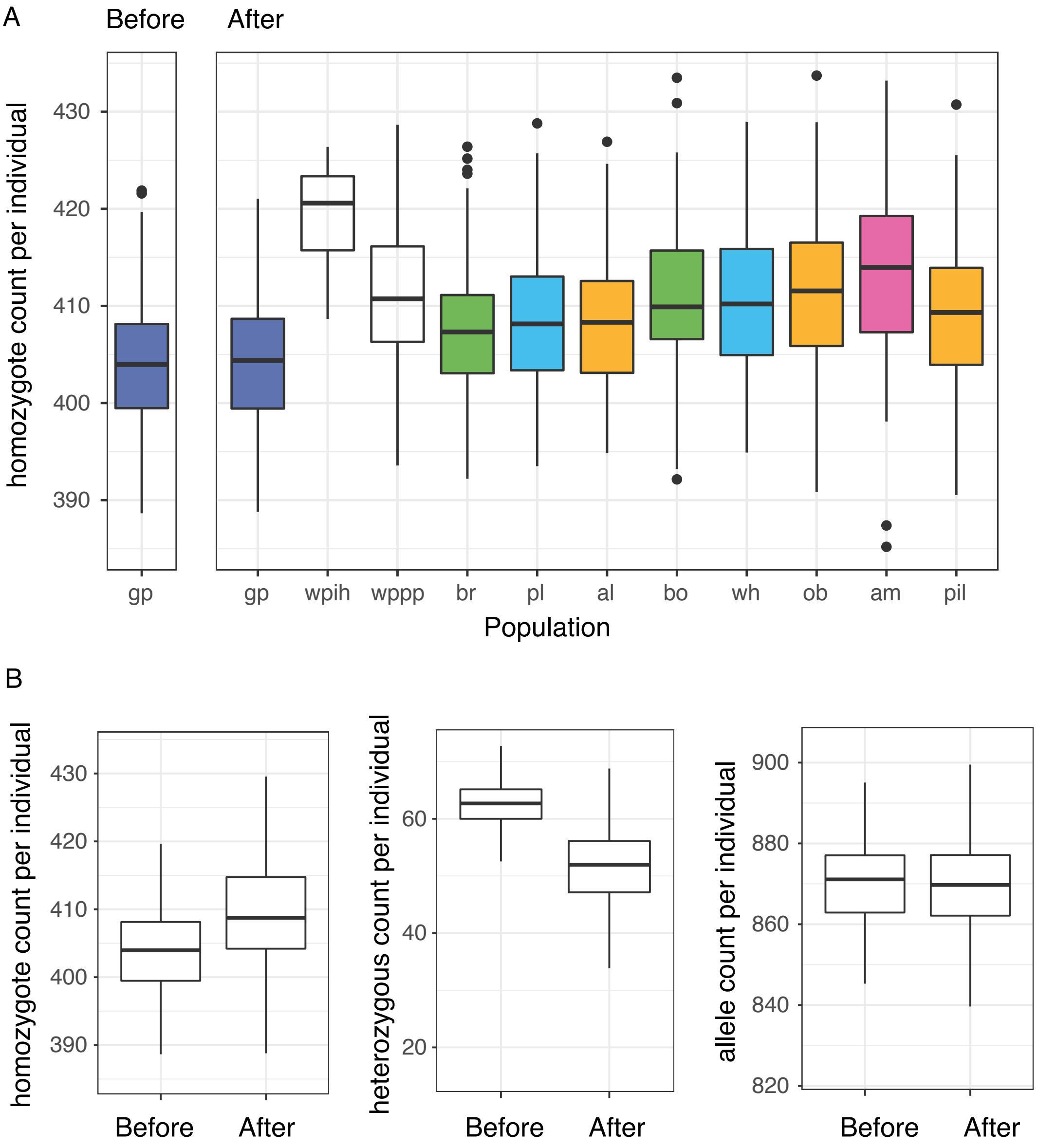

**Figure S21:** Outcome of the demographic simulations. A) Homozygote count for derived neutral alleles before and after the species bottleneck (generations 2999 and 3023, respectively). Simulated populations included all major wild parks and populations that contributed to the founding of the Alpine ibex populations under study (see also Figure 3A). Matching colours group source populations with populations that were founded from them (except for wpih, wppp, gp and am). Abbreviations: gp: Gran Paradiso, wpih: Wildpark Interlaken Harder, wppp: Wildpark Peter and Paul, br: Brienzer Rothorn, pl: Pleureur, al: Albris, bo: Bire Oeschinen, wh: Weisshorn, ob: Oberbauenstock, am: Alpi Marittime, pil: Pilatus. B) Homozygote and heterozygote counts, and allele count per individual of derived neutral alleles before and after the species bottleneck. The category "after" includes all populations for which genomic data was generated with the exception of the wild parks and the Pilatus (pil) population.

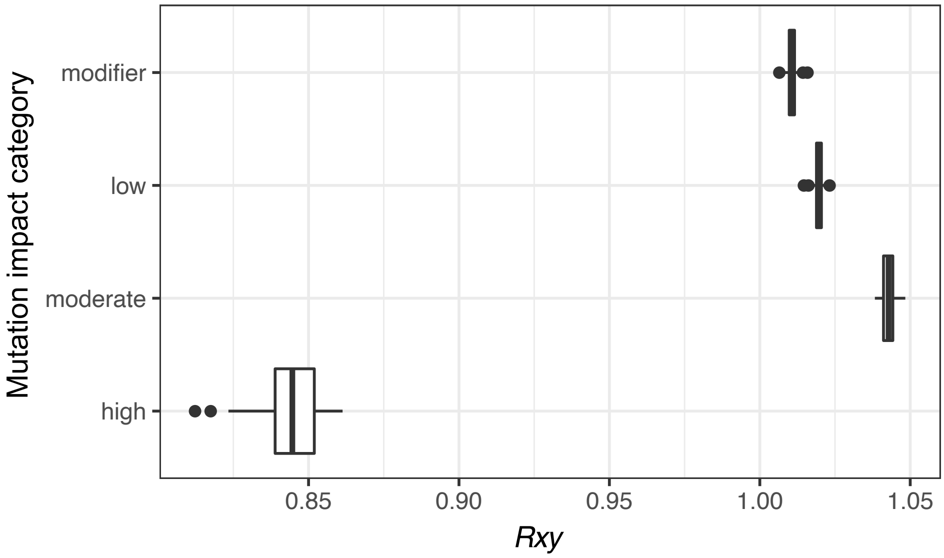

**Figure S22**: Rxy analysis contrasting the strongly bottlenecked Alpi Marittime population with the Gran Paradiso population across the spectrum of SnpEff impact categories.

**Figures S23 – S31:** Visualization of SNP variant quality parameters used for filtering. The applied cut-offs are shown with a vertical line.

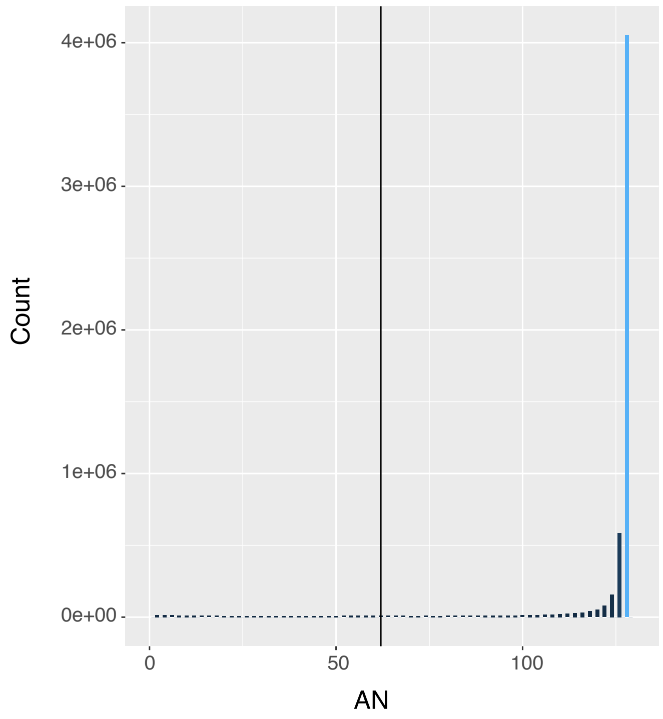

**Figure S23:** Distribution of the AN statistic. Number of called alleles (corresponds to twice the number of samples genotyped at the locus).

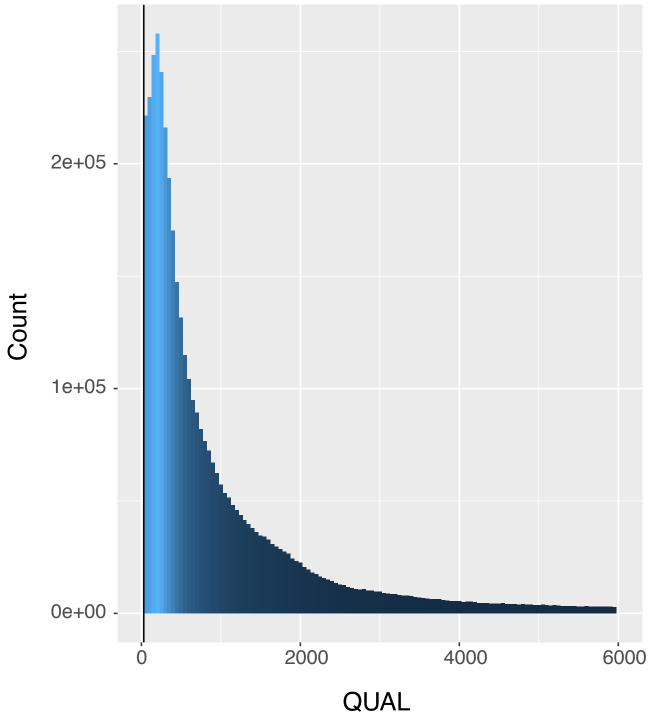

**Figure S24:** Distribution of the overall SNP quality (QUAL) statistic.

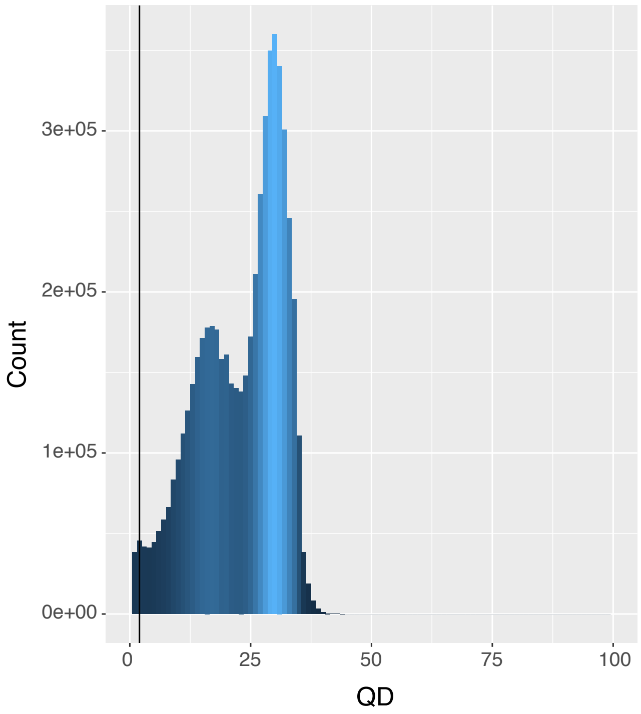

**Figure S25:** Distribution of the QualByDepth (QD) statistic. Confidence of the variant quality (QUAL) divided by the unfiltered depth of the samples called as non-reference.

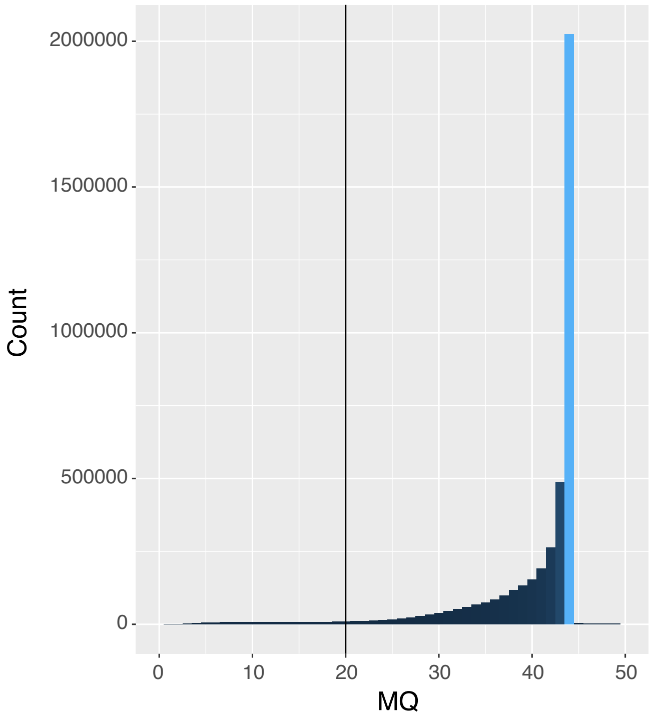

**Figure S26:** Distribution of the RMSMappingQuality (MQ) statistic. Root mean square of the mapping quality calculated across all reads from all samples.

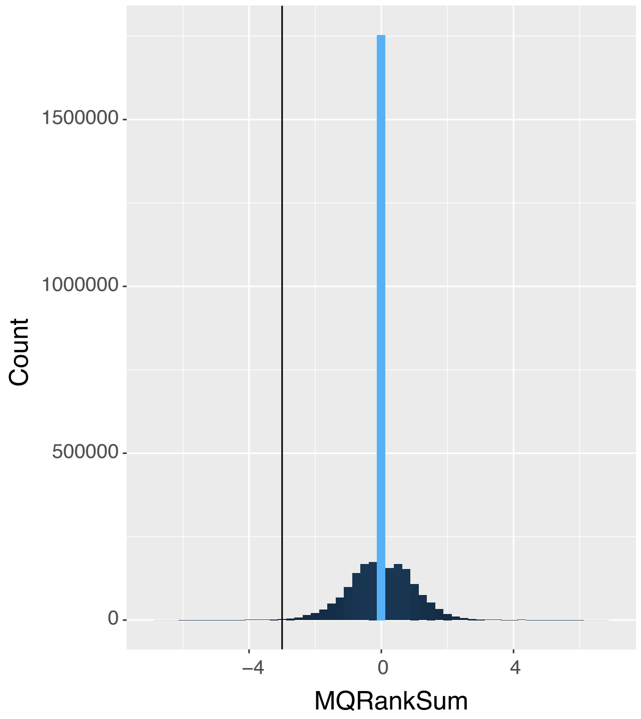

**Figure S27:** Distribution of the MappingQualityRankSumTest (MQRankSum) statistic. A test for differences in mapping quality between reference versus alternate allele.

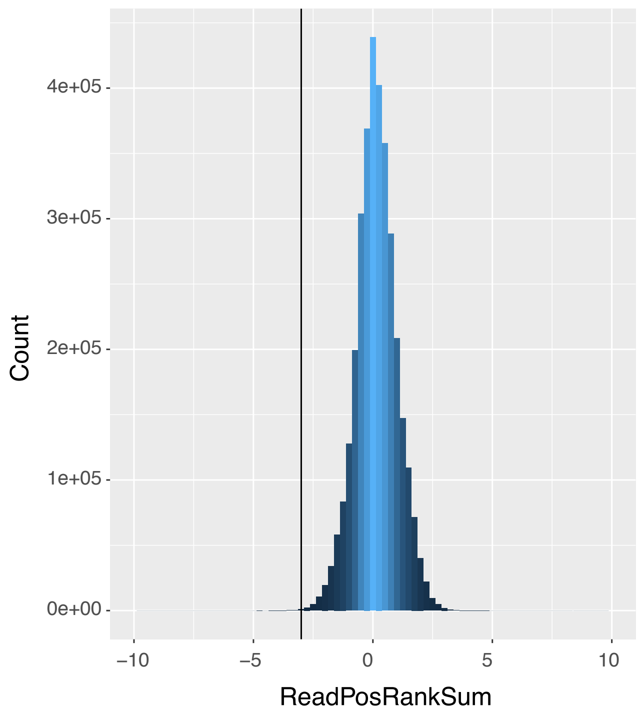

**Figure S28:** Distribution of the ReadPosRankSumTest statistic. A test for the even distribution of variants along reads.

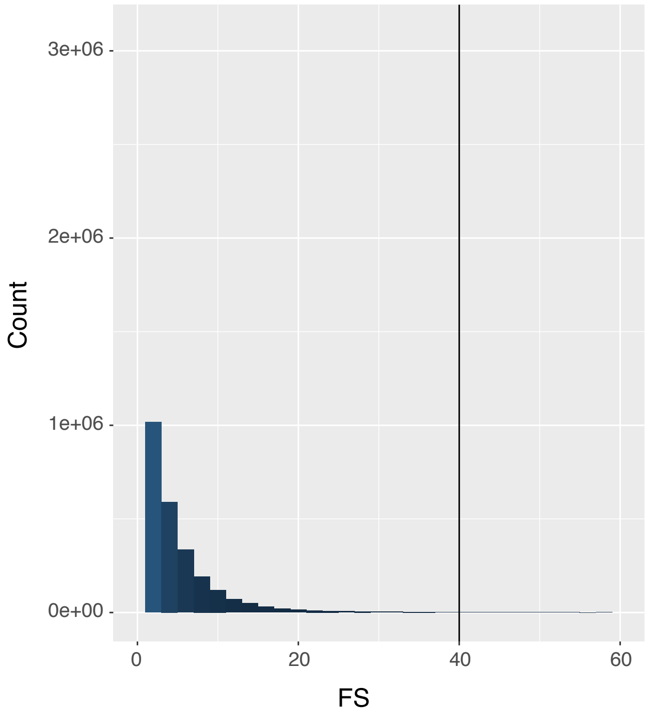

**Figure S29:** Distribution of the FisherStrand (FS) statistic. A test for strand bias, i.e. if a variant is predominantly associated with one of the two read strands (forward or reverse).

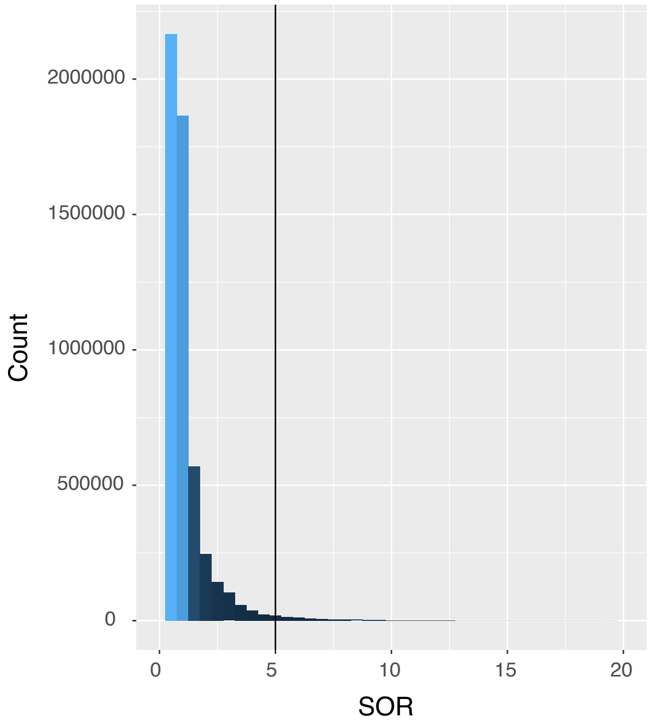

**Figure S30:** Distribution of the StrandOddsRatio (SOR) statistic.

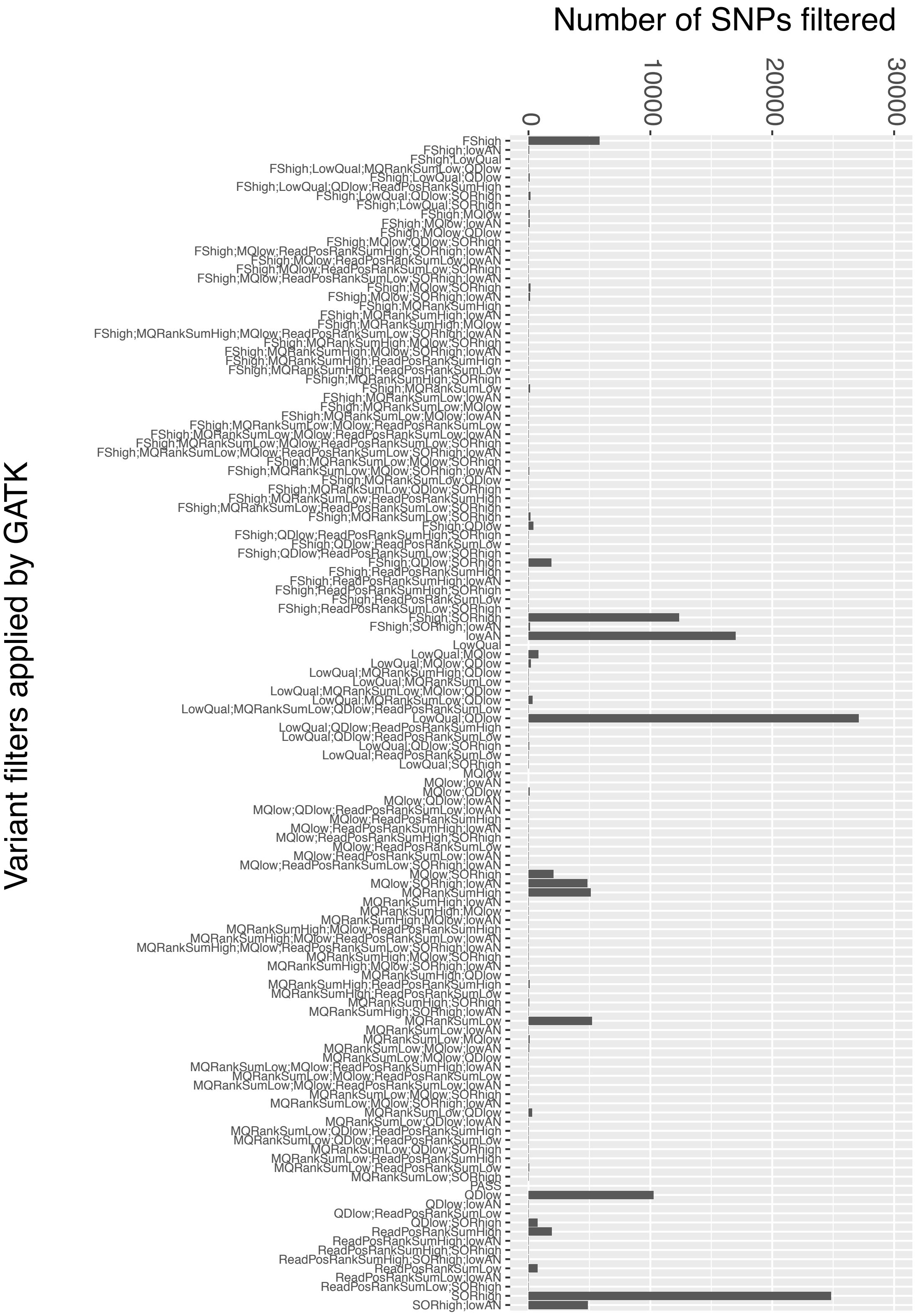

**Figure S31:** Candidate SNP variants filtered out by different filter combinations. See methods for individual filter settings.

**Figures S32 - S39:** Visualization of indel variant quality parameters used for filtering. The applied cut-offs are shown with a vertical line**.**

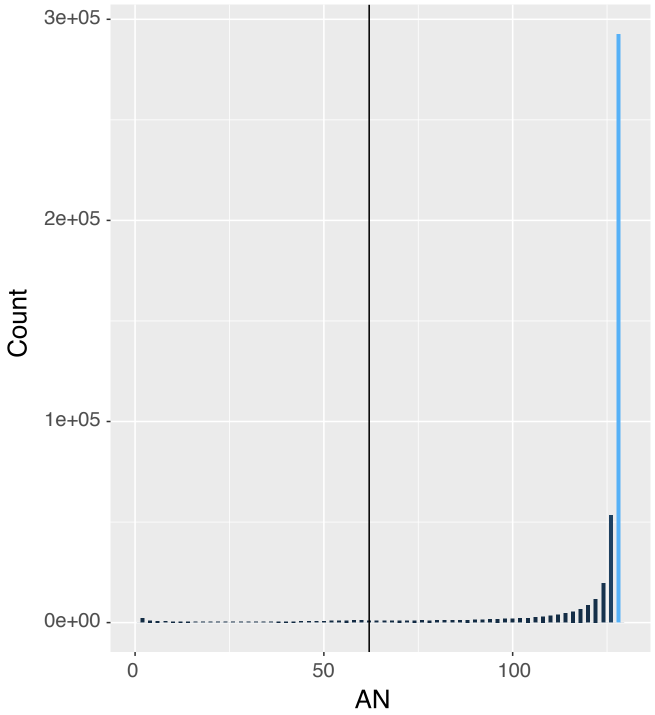

**Figure S32:** Distribution of the AN statistic. Number of called alleles (corresponds to twice the number of samples genotyped at the locus).

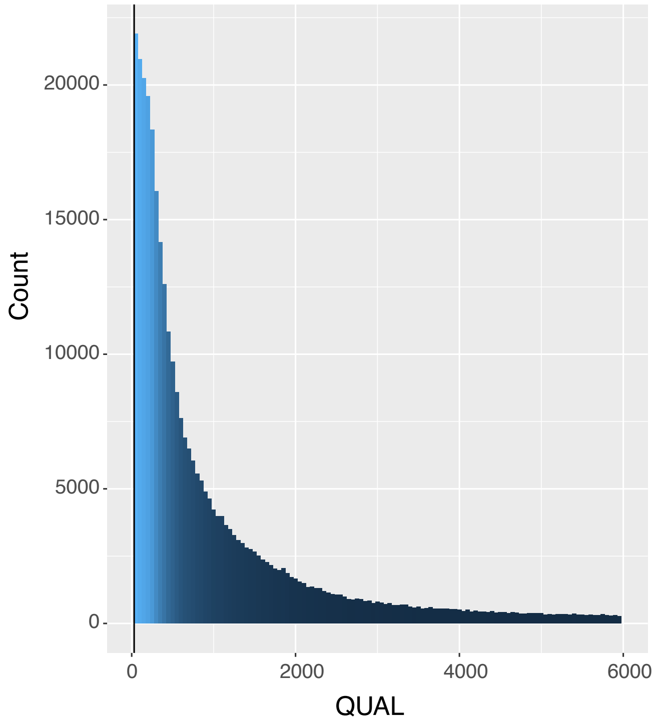

**Figure S33:** Distribution of the overall indel quality (QUAL) statistic.

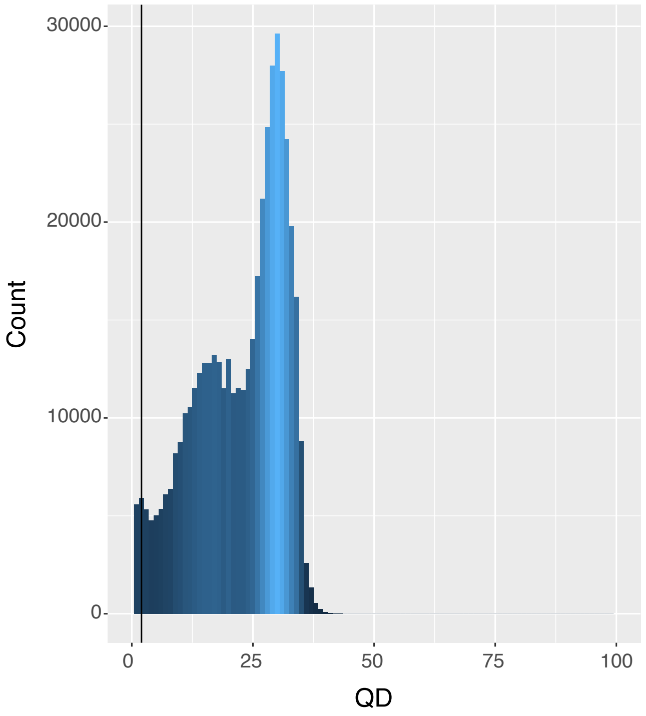

**Figure S34:** Distribution of the QualByDepth (QD) statistic. Confidence of the variant quality (QUAL) divided by the unfiltered depth of the samples called as non-reference.

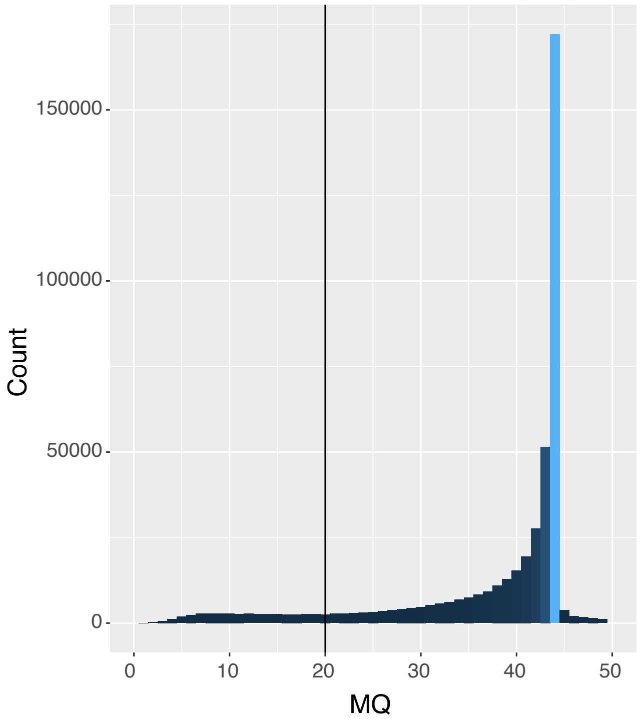

**Figure S35:** Distribution of the RMSMappingQuality (MQ) statistic. Root mean square of the mapping quality calculated across all reads from all samples.

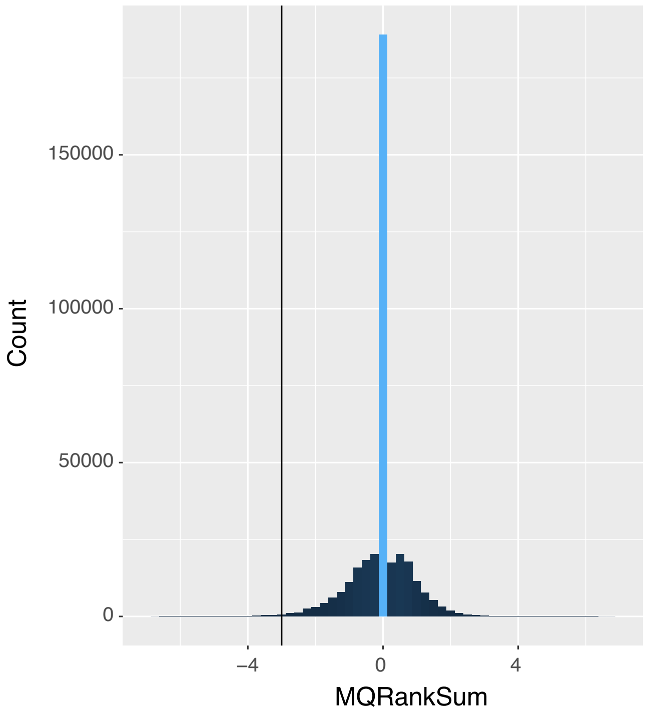

**Figure S36:** Distribution of the MappingQualityRankSumTest (MQRankSum) statistic. A test for differences in mapping quality between reference versus alternate allele.

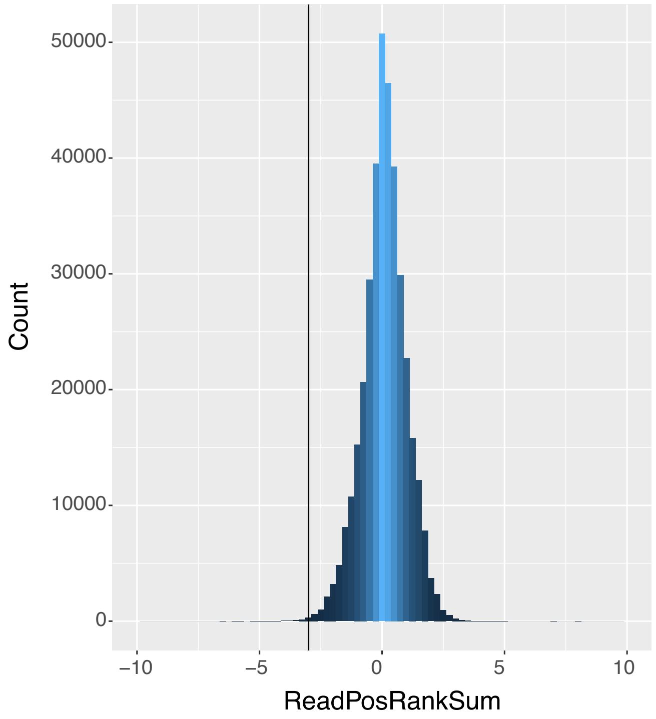

**Figure S37:** Distribution of the ReadPosRankSumTest statistic. A test for the even distribution of variants along reads.

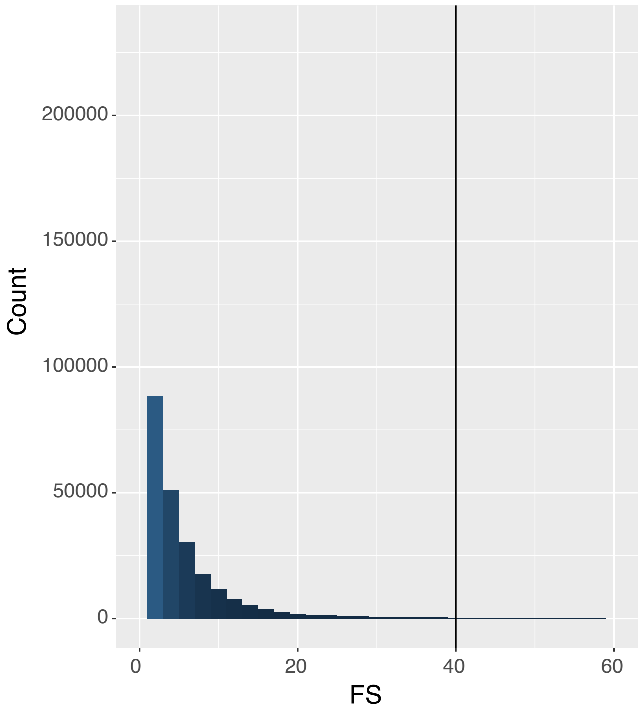

**Figure S38:** Distribution of the FisherStrand (FS) statistic. A test for strand bias, i.e. if a variant is predominantly associated with one of the two read strands (forward or reverse).

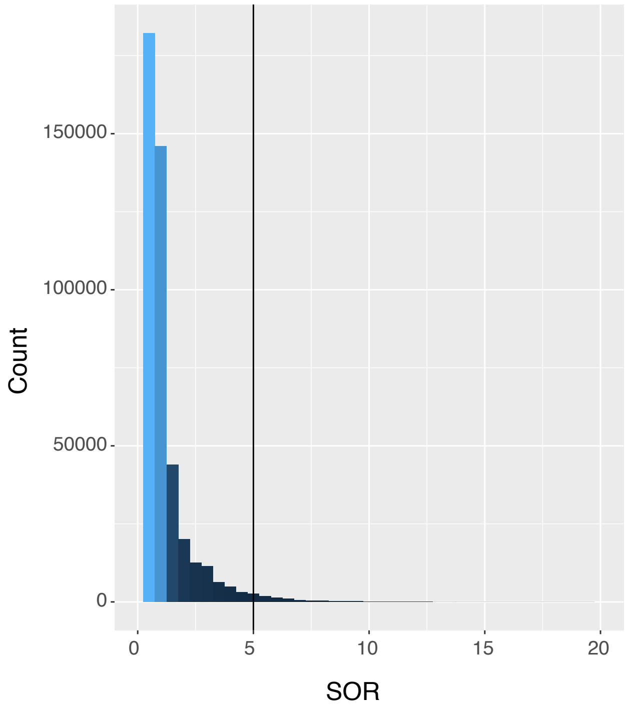

**Figure S39:** Distribution of the StrandOddsRatio (SOR) statistic.

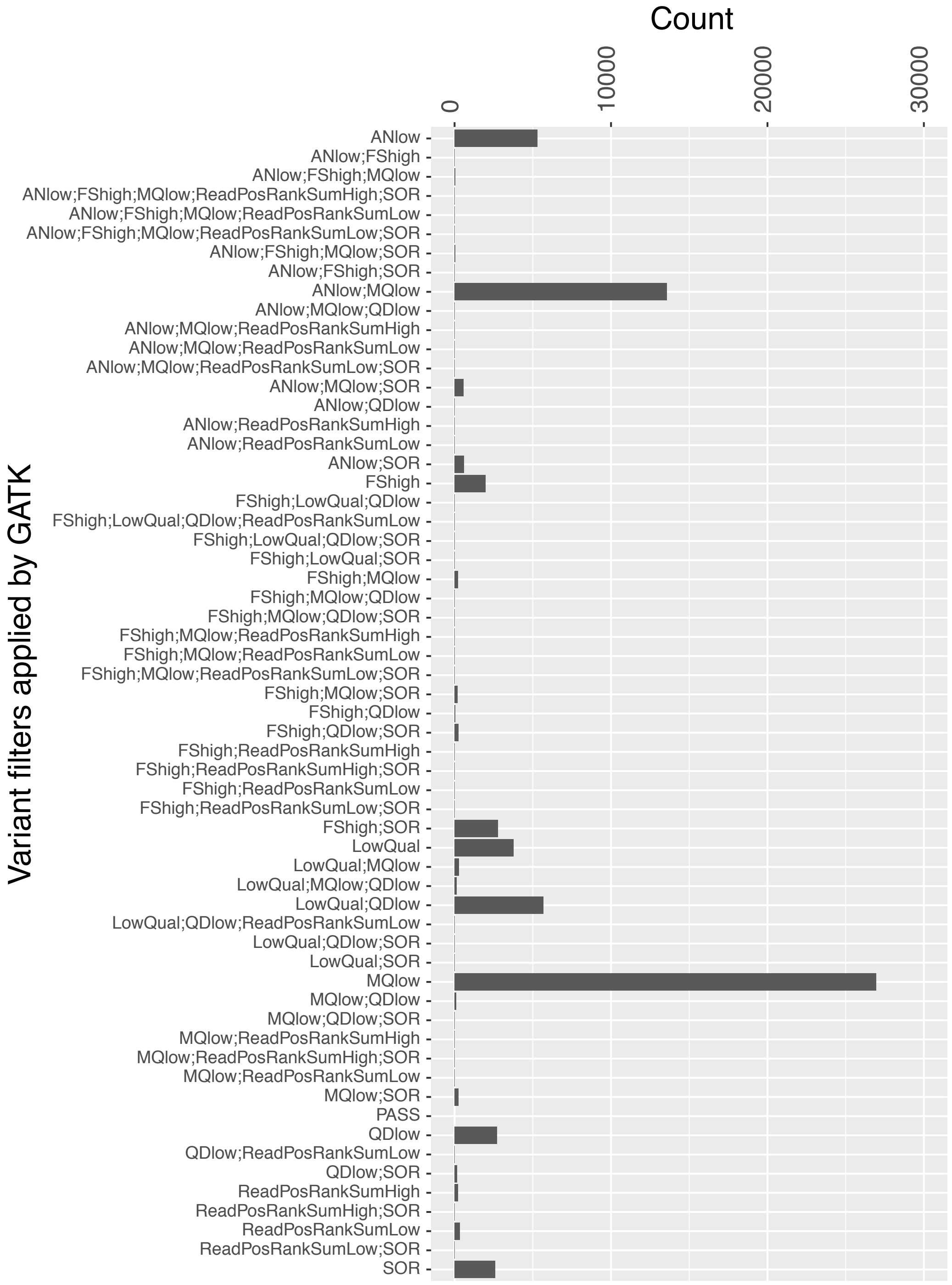

**Figure S40:** Candidate indel variants filtered out by different filter combinations. See methods for individual filter settings.

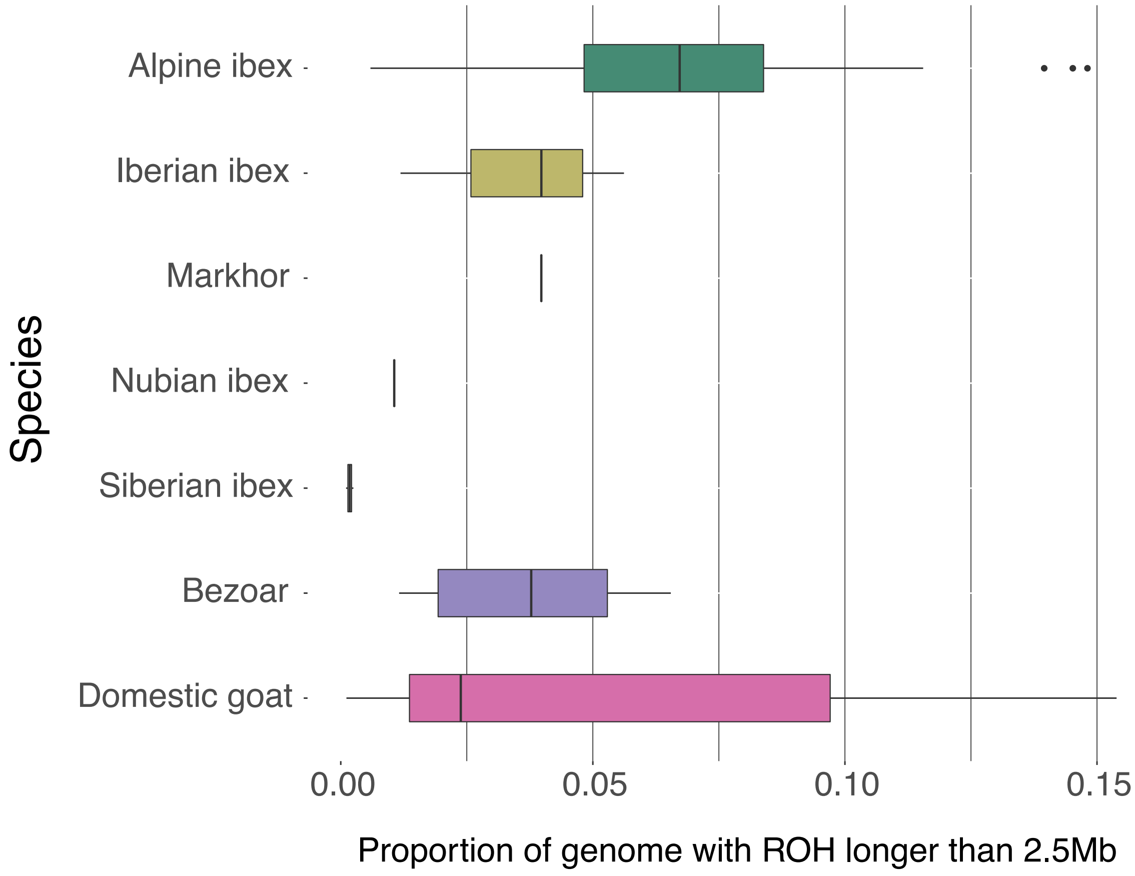

**Figure S41:** Proportion of the genome with runs of homozygosity (ROH) longer than 2.5 Mb estimated using the software PLINK.

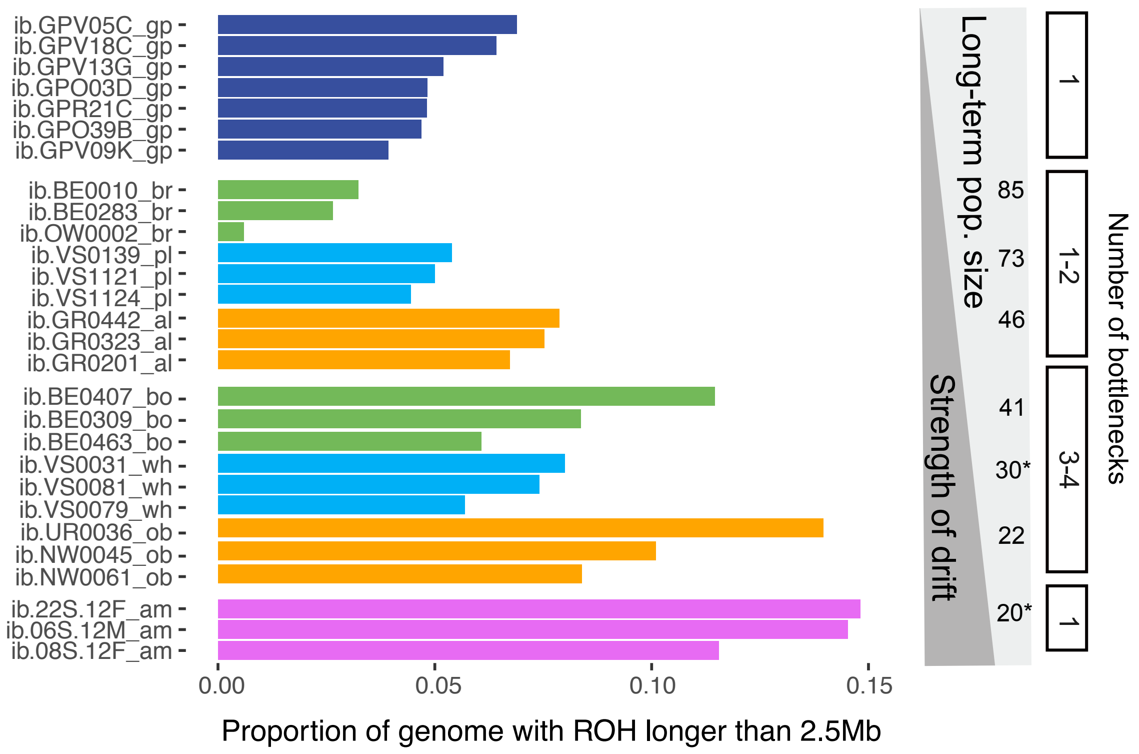

**Figure S42:** Proportion of the genome with runs of homozygosity (ROH) longer than 2.5 Mb estimated using the software PLINK.

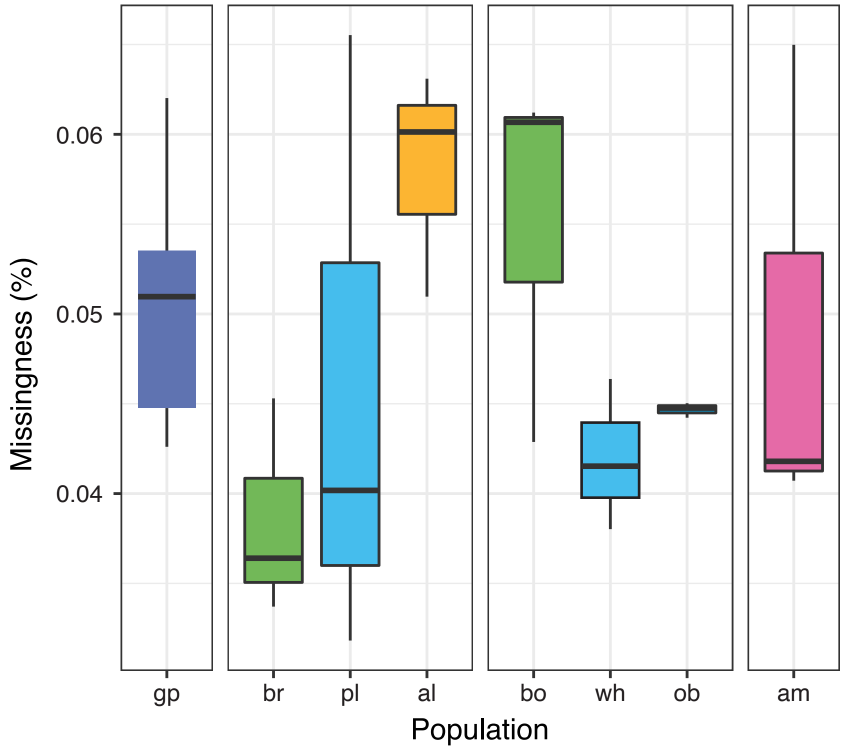

**Figure S43:** Boxplot showing the percentage of loci with missing data (missingness) reported by individual and grouped by population.

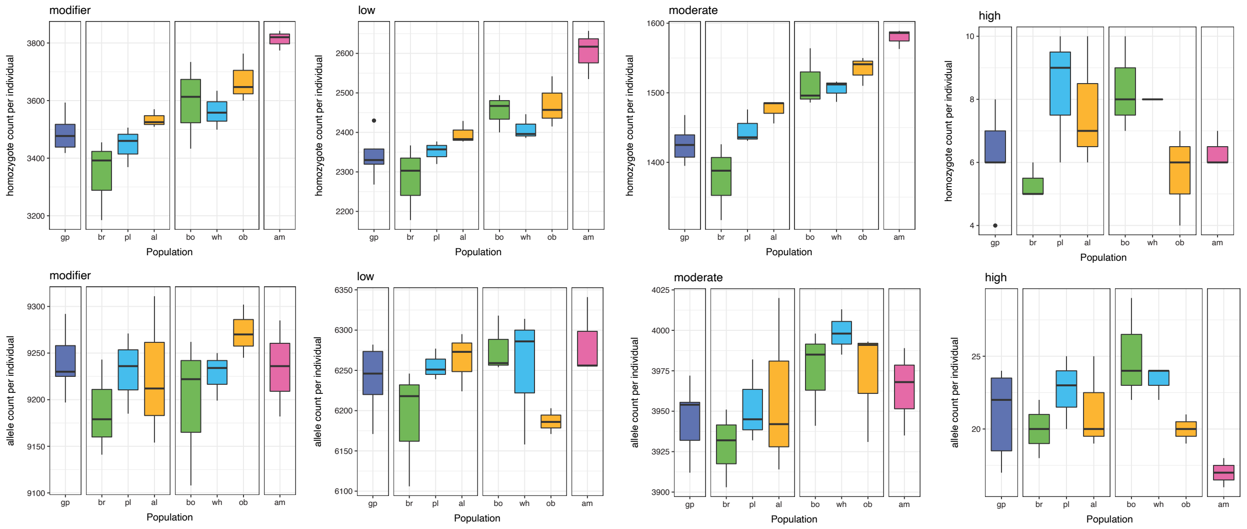

**Figure S44:** Individual homozygote counts (upper panel) and allele counts (lower panel) for each Alpine ibex population with no missingness allowed. Left to right shows SNP mutation category as predicted by SnpEff (modifier, low, moderate and high). Colors group Alpine ibex populations into founder and descendent population pairs (see also Figures 3 and S13A).

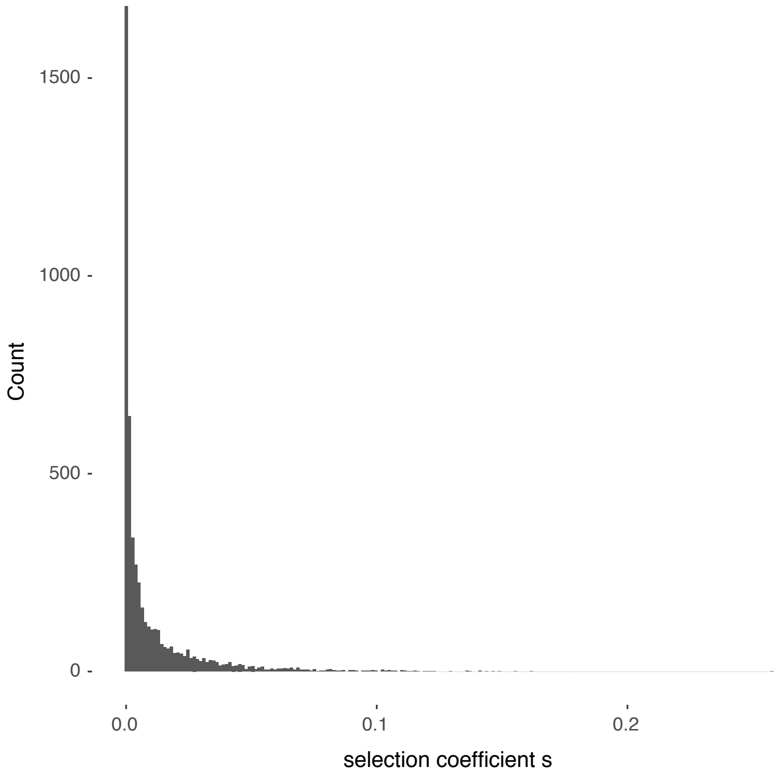

**Figure S45:** Distribution of the selection coefficients *s* used in the simulations.

### SUPPLEMENTARY TABLES

**Table S1A**: Demographic information on Alpine ibex populations. Gran Paradiso was the only surviving population and at the source of all introduced populations. Effective founder group size differs from total released numbers if knowledge was available about mortality after population establishment. Long-term population size was calculated as the harmonic mean of census size since population founding. *) Estimated numbers based on historic record.

**References**

1. Piodi, M., 1984. Le Bouquetin des Alpes - *Capra ibex*. In: C.I.C. Kommission Grosswild (Editor), Der Steinbock in Eurasien. C.I.C. Symposium, Pontresina, Switzerland, 24-25 February 1984
2. Stuwe, M. & Nievergelt, B. Recovery of alpine ibex from near extinction - the result of effective protection, captive breeding, and reintroductions. Applied Animal Behaviour Science **29,** 379–387 (1991).
3. Aeschbacher, S., Futschik, A. & Beaumont, M. A. Approximate Bayesian computation for modular inference problems with many parameters: the example of migration rates. Mol Ecol **22,** 987–1002 (2013). DRYAD entry doi:10.5061/dryad.274b1
4. Terrier, G. & Rossi, P. Le bouquetin (*Capra ibex ibex*) dans les alpes maritimes franco-italiennes: occupation de l'éspace, clonisation et régulation naturelles. Traveaux Scientifiques du Parc National de la Vanoise **XVIII,** 271–288 (1994).
5. Alpi Marittime Natural Park
6. Biebach, I. & Keller, L. F. Inbreeding in reintroduced populations: the effects of early reintroduction history and contemporary processes. Conserv Genet **11,** 527–538 (2010).

| **Alpine ibex** [extant census 50'000 individuals] | **Date of foundation** | **Males released** | **Females released** | **Total released** | **Effective founder group size** | **Long-term population size** (from foundation up to 2007) | **Census 2007** |
|---|---|---|---|---|---|---|---|
| **Population** | | | | | | | |
| **Gran Paradiso** [1,2] | | - | - | - | 100* | n/a | n/a |
| **Albris (al)** [3] | 1920-34 | 16 | 26 | 42 | 42 | 46 | 1033 |
| **Alpi Marittime (am)** [4,5] | 1920-30 | | | 25 | 6 | 20* | 750 |
| **Bire Oeschinen (bo)** [3] | 1961-62 | 4 | 9 | 13 | 13 | 41 | 68 |
| **Brienzer Rothorn (br)** [3] | 1921-25 | 8 | 10 | 18 | 18 | 85 | 288 |
| **Oberbauenstock (ob)** [3] | 1968-86 | 10 | 12 | 22 | 22 | 22 | 164 |
| **Pilatus (pi)** [3] | 1961-65 | 8 | 7 | (unknown sex: 2) 17 | 17 | 43 | 104 |
| **Pleureur (pl)** [3] | 1928-35 | 6 | 9 | 15 | 15 | 73 | (year 2003) 806 |
| **Weisshorn (wh)** [6] | 1962-69 | 18 | 6 | 24 | 24 | 30* | 354 |

**Table S1B**: Demographic information on three major Iberian ibex populations including both extant subspecies of *Capra pyrenaica.* Bottleneck periods represent the most dramatic, known reductions in population size. Sierra Nevada and Maestrazgo populations experienced only recent bottlenecks (1960s).

**References**

1. Couturier M. Le bouquetin des Alpes. Edited by author, Grenoble, France, (1962).
2. Pérez JM, Granados JE, Soriguer RC, Fandos P, Márquez FJ, Crampe JP, et al. Distribution, status and conservation problems of the Spanish Ibex, *Capra pyrenaica* (Mammalia: Artiodactyla). Mammal Review **32,** 26–39 (2002);.
3. Peña, J. *La cabra montés en España.* International Game Congress, Switzerland (1978).
4. Ortuño, F. & Peña, J. Reservas y cotos nacionales de caza. INCAFO, Madrid, UK. (1979)
5. Alados CL. Distribution and status of the Spanish ibex (*Capra pyrenaica*, Schinz). In: The Biology and Management of Mountain Ungulates (Ed. by Lovari S.), pp. 204–211. Croom-Helm, Beckenham London, UK, (1985).

| **Iberian ibex** | **Subspecies** | **Strengths of known bottlenecks periods** [estimated individuals] | **Population size changes over time** [if available] | **Current population size** |
|---|---|---|---|---|
| **Sierra Nevada (sn)** | *Capra pyrenaica hispanica* | ~600 (1960s)[1] | n/a | 16’000[2] |
| **Maestrazgo (hb)** | *Capra pyrenaica hispanica* | ~30 (1960s)[1] | 450 (1966)[2]<br>700 (1970)[2-5]<br>>3000 (1977)[2-5]<br>3800 (1979)[2-5]<br>> 5000 (1981)[2-5] | 7000[1] |
| **Sierra de Gredos (vi)** | *Capra pyrenaica victoriae* | 50 (1895)[1]<br>12 (1905)[1]<br>350 (1912)[1]<br>500 (1914)[1]<br>>1000 (1917)[1] | 2150 (1961)[1]<br>4000 (1976)[2-4]<br>9000 (1978)[2-4]<br>8000 (1993)[2-4] | 8000[1] |

**Table S2**: Overview of all sequenced individuals grouped by species including the IUCN conservation status and recent demographic events. Whole-genome sequencing data, reference genome mapping and single nucleotide polymorphism (SNP) statistics are shown.

| Sample | Population | Sex | Location | Coordinates (Swiss grid or latitude/longitude) | | Sampling date | Raw sequencing data (Gbp) | Mapping rate (%) | Coverage (X) | Number of SNPs genotyped | Number of heterozygote SNPs |
|---|---|---|---|---|---|---|---|---|---|---|---|
| **Alpine ibex: Recent bottlenecks, IUCN: Least concern**<br>N=29; 3'539'135 SNPs; nucleotide diversity: 0.00039 | | | | | | | | | | | |
| **ib.06S.12M_am** | Alpi Maritimi | male | Rovina | | | 5.2012 | 60.0 | 99.0 | 17.5 | 56'126'898 | 667'046 |
| **ib.08S.12F_am** | Alpi Maritimi | female | Matto | | | 5.2012 | 65.2 | 98.9 | 18.8 | 56'126'955 | 669'730 |
| **ib.22S.12F_am** | Alpi Maritimi | female | Barra | | | 5.2012 | 60.2 | 99.1 | 17.6 | 56'126'911 | 660'179 |
| **ib.BE0010_br** | Brienzer Rothorn | female | Stelli, Hofstetten | 181400 | 649600 | 2.9.2005 | 64.3 | 99.2 | 19.5 | 56'127'089 | 938'208 |
| **ib.BE0283_br** | Brienzer Rothorn | female | Salenwang | 180600 | 650250 | 7.11.2006 | 69.5 | 99.1 | 21.1 | 56'127'180 | 954'605 |
| **ib.BE0309_bo** | Bire Oeschinen | female | Schwarzhorn Oeschinen | 151400 | 624850 | 25.10.2006 | 59.6 | 98.7 | 18.1 | 56'126'958 | 803'869 |
| **ib.BE0407_bo** | Bire Oeschinen | male | Oeschinensee, Heuberg | 150750 | 621800 | 13.6.2007 | 55.5 | 99.1 | 17.2 | 56'126'864 | 753'333 |
| **ib.BE0463_bo** | Bire Oeschinen | male | Oeschinensee, Underbärgli | 150750 | 622600 | 23.5.2008 | 59.7 | 99.1 | 18.7 | 56'126'974 | 889'571 |
| **ib.GPO03D_gp** | Gran Paradiso | male | Orco, Chiapili sup | 45.4684 | 7.1474 | 1.1.2004 | 58.8 | 98.9 | 17.3 | 56'126'869 | 906'666 |
| **ib.GPO39B_gp** | Gran Paradiso | male | Orco | 45.4644 | 7.1588 | 1.1.2002 | 58.3 | 99.0 | 17.2 | 56'126'868 | 958'991 |
| **ib.GPR21C_gp** | Gran Paradiso | male | Rhemes | 45.5637 | 7.1400 | 1.1.2003 | 57.2 | 99.2 | 17.7 | 56'126'849 | 925'983 |
| **ib.GPV05C_gp** | Gran Paradiso | female | Valsaveranche, Levionaz | 45.5821 | 7.2262 | 1.1.2003 | 60.9 | 99.2 | 18.4 | 56'126'978 | 834'684 |
| **ib.GPV09K_gp** | Gran Paradiso | male | Valsaveranche, Levionaz | 45.5746 | 7.2398 | 10.6.2014 | 56.4 | 99.0 | 17.0 | 56'126'827 | 924'811 |
| **ib.GPV13G_gp** | Gran Paradiso | male | Valsaveranche, Levionaz | 45.5779 | 7.2333 | 10.6.2014 | 56.8 | 99.0 | 16.6 | 56'126'804 | 900'629 |
| **ib.GPV18C_gp** | Gran Paradiso | male | Valsaveranche, Levionaz | 45.5779 | 7.2333 | 1.1.2003 | 63.5 | 99.1 | 20.0 | 56'127'046 | 889'186 |
| **ib.GR0140_rh** | Rheinwaldhorn | female | Scalutta Gem Safien | 166250 | 741820 | 6.10.2005 | 54.3 | 98.7 | 16.8 | 56'126'827 | 733'524 |
| **ib.GR0201_al** | Albris | female | Munt Blais, Gm S-chanf | 166065 | 799210 | 11.11.2004 | 56.1 | 99.2 | 17.4 | 56'126'937 | 854'091 |
| **ib.GR0323_al** | Albris | female | Paradies Suot, Pontresina | 150740 | 791325 | 10.10.2005 | 60.8 | 99.1 | 18.3 | 56'126'987 | 856'238 |
| **ib.GR0442_al** | Albris | male | Val da Fain Tschüffer, Pontresina | 150160 | 797230 | 18.10.2005 | 60.2 | 99.0 | 18.6 | 56'127'038 | 818'724 |
| **ib.NW0045_ob** | Oberbauerstock | male | Hoh Brisen | 194450 | 678200 | 1.9.2006 | 55.6 | 98.9 | 17.3 | 56'126'878 | 768'646 |
| **ib.NW0061_ob** | Oberbauerstock | female | Niederbauen | 200200 | 685050 | 29.9.2007 | 58.9 | 99.2 | 17.8 | 56'126'939 | 787'534 |
| **ib.OW0002_br** | Brienzer Rothorn | male | Eisee, Giswil | 182251 | 646972 | 8.8.2006 | 63.8 | 98.6 | 19.6 | 56'127'113 | 1'102'820 |
| **ib.UR0036_ob** | Oberbauerstock | female | IsenthalBärenstock | 193371 | 681175 | 25.10.2005 | 61.0 | 97.8 | 18.1 | 56'126'939 | 695'986 |
| **ib.VS0031_wh** | Weisshorn | female | Jungen; St. Nklaus | 115700 | 627300 | 23.10.2004 | 56.5 | 99.1 | 17.4 | 56'126'843 | 842'412 |
| **ib.VS0079_wh** | Weisshorn | female | Seetal, Gde. Grächen | 117000 | 627750 | 18.7.2005 | 62.3 | 99.1 | 20.2 | 56'127'095 | 876'610 |
| **ib.VS0081_wh** | Weisshorn | female | Jungen, Gde. St. Niklaus | 117000 | 627750 | 9.8.2005 | 60.0 | 99.0 | 19.5 | 56'126'935 | 805'493 |
| **ib.VS0139_pl** | Pleureur | male | Dixènce-Evolène | 101000 | 595000 | 20.6.2005 | 62.0 | 99.0 | 19.4 | 56'127'046 | 916'987 |
| **ib.VS1121_pl** | Pleureur | female | Verbier | 105100 | 586700 | 19.10.2007 | 60.9 | 98.9 | 18.4 | 56'126'971 | 864'945 |
| **ib.VS1124_pl** | Pleureur | male | Boussine | 87250 | 592900 | 15.10.2007 | 54.4 | 98.8 | 17.0 | 56'126'830 | 947'985 |
| **Iberian ibex: Recent bottlenecks, IUCN: Least concern**<br>N=4; 4'353'580 SNPs; nucleotide diversity: 0.00066 | | | | | | | | | | | |
| **py.V53_vi** | | | Sierra de Gredos | | | 2004-2007 | 60.0 | 91.3 | 15.3 | 56'126'235 | 1'274'362 |
| **py.V66_vi** | | | Sierra de Gredos | | | 2004-2007 | 60.3 | 94.2 | 15.3 | 56'126'292 | 1'000'193 |
| **py.M518_sn** | Sierra Nevada | | Sierra Nevada | | | 2003-2008 | 88.0 | 99.1 | 26.9 | 56'127'272 | 1'000'228 |
| **py.Z5_hb** | | | Maestrazgo | | | 2003-2008 | 56.4 | 99.2 | 17.0 | 56'126'788 | 960'783 |
| **Nubian ibex: Fragmented populations, IUCN: Vulnerable**<br>N=2, 6'980'112 SNPs, nucleotide diversity: 0.00151 | | | | | | | | | | | |
| **nu.ibx20** | | | Sinai, Egypt | | | . | 102.5 | 98.6 | 17.1 | 56'126'494 | 3'131'909 |
| **nu.ibx61** | | | Howtat, Central Saudi Arabia | | | . | 46.9 | 98.7 | 18.4 | 56'126'885 | 2'536'433 |
| **Markhor: Fragmented populations, IUCN: Near Threatened**<br>N=1, 1'988'577 SNPs, nucleotide diversity: 0.00077 | | | | | | | | | | | |
| **fal.Cafal1** | | | Dasht-i Jum Reserve, Tajikistan | | | 24.11.2015 | 148.7 | 98.9 | 38.6 | 56'127'403 | 1'845'346 |
| **Siberian ibex: Large populations, IUCN: Least concern**<br>N=2, 10'433'689 SNPs, nucleotide diversity: 0.00224 | | | | | | | | | | | |
| **si.Casi1** | | male | South of Issyk lake | | | 2010 | 57.4 | 98.7 | 15.8 | 56'126'287 | 4'379'849 |
| **si.Casi2** | | male | Dasht-i Jum Reserve, Tajikistan | | | 22.11.2015 | 80.1 | 98.8 | 22.0 | 56'126'976 | 4'823'262 |
| **Bezoar: Large populations, IUCN: Vulnerable**<br>N=6, 15'182'016 SNPs, nucleotide diversity: 0.00191 | | | | | | | | | | | |
| **IRCA-C3-1001** | | male | Marakan | 45.16 | 38.63 | 30.08.10 | 37.6 | 99.1 | 11.5 | 56'127'088 | 3'212'565 |
| **IRCA-G2-5063** | | male | Arasbaran | 47.13 | 39.18 | 17.12.09 | 36.5 | 98.7 | 11.6 | 56'127'115 | 3'610'951 |
| **IRCA-I11-0001** | | male | Khan Gormaz | 48.20 | 34.60 | 29.01.11 | 19.1 | 99.0 | 5.9 | 56'126'263 | 2'293'061 |
| **IRCA-I6-5237** | | male | Karnagh | 48.39 | 37.16 | 05.10.11 | 34.6 | 99.1 | 11.3 | 56'127'111 | 3'402'754 |

**Table S3:** Total length of runs of homozygosity (ROH) per ROH category. Note that categories are overlapping. Populations abbreviated by am: Alpi Marittime, gp: Gran Paradiso; al: Albris; bo: Bire Öschinen; br: Brienzer Rothorn; ob: Oberbauenstock; pl: Pleureur; rh: Rheinwald; wh: Weisshorn. NA: no population/breed assigned. A) Analysis settings af.dflt 0.4, M=0.0000012, run with Viterbi. B) Settings af.dflt 0.4, M=0.0000012.

| Individual | 100-500 Kb | >100 kb | >500 kb | >2.5 Mb | 2.5-10 Mb | >10 Mb | Population/ Breed | Species |
|---|---|---|---|---|---|---|---|---|
| | Settings: af.dflt 0.4, M=0.0000012, with Viterbi | | | | | | | |
| **ib.GR0201_al** | 617'316'226 | 1'499'036'113 | 881'719'887 | 278'054'595 | 278'054'595 | 0 | al | *C. ibex* |
| **ib.GR0323_al** | 648'739'982 | 1'568'056'539 | 919'316'557 | 271'420'074 | 271'420'074 | 0 | al | *C. ibex* |
| **ib.GR0442_al** | 638'635'462 | 1'619'373'279 | 980'737'817 | 293'579'234 | 293'579'234 | 0 | al | *C. ibex* |
| **ib.08S.12F_am** | 550'622'660 | 1'883'994'916 | 1'333'372'256 | 580'980'545 | 569'620'719 | 11'359'826 | am | *C. ibex* |
| **ib.06S.12M_am** | 526'257'448 | 1'814'566'314 | 1'288'308'866 | 531'521'579 | 531'521'579 | 0 | am | *C. ibex* |
| **ib.22S.12F_am** | 485'235'630 | 1'826'304'781 | 1'341'069'151 | 604'013'148 | 604'013'148 | 0 | am | *C. ibex* |
| **ib.BE0309_bo** | 624'378'681 | 1'625'733'750 | 1'001'355'069 | 325'649'131 | 314'522'057 | 11'127'074 | bo | *C. ibex* |
| **ib.BE0407_bo** | 529'902'205 | 1'628'035'706 | 1'098'133'501 | 444'950'345 | 444'950'345 | 0 | bo | *C. ibex* |
| **ib.BE0463_bo** | 714'827'140 | 1'586'641'537 | 871'814'397 | 233'878'805 | 233'878'805 | 0 | bo | *C. ibex* |
| **ib.BE0010_br** | 743'914'314 | 1'653'878'572 | 909'964'258 | 238'950'631 | 238'950'631 | 0 | br | *C. ibex* |
| **ib.BE0283_br** | 808'881'190 | 1'775'794'639 | 966'913'449 | 264'644'590 | 254'642'196 | 10'002'394 | br | *C. ibex* |
| **ib.OW0002_br** | 761'260'031 | 1'667'152'587 | 905'892'556 | 199'762'116 | 199'762'116 | 0 | br | *C. ibex* |
| **ib.GPR21C_gp** | 678'473'743 | 1'434'407'783 | 755'934'040 | 175'346'650 | 175'346'650 | 0 | gp | *C. ibex* |
| **ib.GPV05C_gp** | 650'121'652 | 1'594'127'217 | 944'005'565 | 266'705'862 | 266'705'862 | 0 | gp | *C. ibex* |
| **ib.GPV18C_gp** | 777'624'987 | 1'682'216'743 | 904'591'756 | 236'158'730 | 236'158'730 | 0 | gp | *C. ibex* |
| **ib.GPO03D_gp** | 664'865'678 | 1'520'317'013 | 855'451'335 | 227'583'908 | 227'583'908 | 0 | gp | *C. ibex* |
| **ib.GPO39B_gp** | 709'970'926 | 1'471'650'951 | 761'680'025 | 151'298'231 | 151'298'231 | 0 | gp | *C. ibex* |
| **ib.GPV09K_gp** | 662'015'971 | 1'492'528'666 | 830'512'695 | 167'263'444 | 167'263'444 | 0 | gp | *C. ibex* |
| **ib.GPV13G_gp** | 647'435'577 | 1'493'675'120 | 846'239'543 | 194'612'790 | 194'612'790 | 0 | gp | *C. ibex* |
| **ib.NW0045_ob** | 573'664'491 | 1'615'991'379 | 1'042'326'888 | 370'839'466 | 360'195'918 | 10'643'548 | ob | *C. ibex* |
| **ib.NW0061_ob** | 595'069'740 | 1'616'525'100 | 1'021'455'360 | 336'713'979 | 325'950'950 | 10'763'029 | ob | *C. ibex* |
| **ib.UR0036_ob** | 561'427'932 | 1'762'789'124 | 1'201'361'192 | 514'766'948 | 504'014'248 | 10'752'700 | ob | *C. ibex* |
| **ib.VS0139_pl** | 746'261'010 | 1'630'720'974 | 884'459'964 | 225'660'318 | 225'660'318 | 0 | pl | *C. ibex* |
| **ib.VS1121_pl** | 634'868'136 | 1'645'766'132 | 1'010'897'996 | 326'843'120 | 326'843'120 | 0 | pl | *C. ibex* |
| **ib.VS1124_pl** | 636'991'188 | 1'368'015'802 | 731'024'614 | 139'861'308 | 139'861'308 | 0 | pl | *C. ibex* |
| **ib.GR0140_rh** | 543'598'033 | 1'597'032'136 | 1'053'434'103 | 324'940'186 | 324'940'186 | 0 | rh | *C. ibex* |
| **ib.VS0079_wh** | 738'754'492 | 1'697'873'813 | 959'119'321 | 273'245'245 | 249'102'537 | 24'142'708 | wh | *C. ibex* |
| **ib.VS0081_wh** | 656'387'409 | 1'725'855'351 | 1'069'467'942 | 333'444'887 | 333'444'887 | 0 | wh | *C. ibex* |
| **ib.VS0031_wh** | 635'312'736 | 1'548'163'493 | 912'850'757 | 301'607'030 | 291'518'068 | 10'088'962 | wh | *C. ibex* |
| **nu.ibx20** | 355'804'025 | 398'512'218 | 42'708'193 | 0 | 0 | 0 | NA | *C. nubiana* |
| **nu.ibx61** | 386'976'121 | 803'470'296 | 416'494'175 | 130'605'348 | 130'605'348 | 0 | NA | *C. nubiana* |
| **py.Z5_hb** | 391'166'242 | 1'262'581'533 | 871'415'291 | 281'367'004 | 256'892'451 | 24'474'553 | hb | *C. pyrenaica* |
| **py.M518_sn** | 950'904'052 | 2'037'382'478 | 1'086'478'426 | 303'317'928 | 303'317'928 | 0 | sn | *C. pyrenaica* |
| **py.V53_vi** | 579'842'010 | 1'102'731'313 | 522'889'303 | 47'817'848 | 47'817'848 | 0 | vi | *C. pyrenaica* |
| **py.V66_vi** | 508'832'531 | 1'164'493'864 | 655'661'333 | 117'445'961 | 117'445'961 | 0 | vi | *C. pyrenaica* |
| **si.Casi1** | 129'673'268 | 265'145'849 | 135'472'581 | 42'751'180 | 42'751'180 | 0 | NA | *C. sibirica* |
| **si.Casi2** | 387'043'290 | 544'294'224 | 157'250'934 | 15'395'187 | 15'395'187 | 0 | NA | *C. sibirica* |
| **IRCA-C3-1001** | 116'433'824 | 606'632'709 | 490'198'885 | 268'128'032 | 256'267'102 | 11'860'930 | NA | *C. aegagrus* |
| **IRCA-G2-5063** | 146'599'124 | 364'865'275 | 218'266'151 | 65'684'795 | 65'684'795 | 0 | NA | *C. aegagrus* |
| **IRCA-I11-0001** | 82'755'412 | 828'874'800 | 746'119'388 | 374'898'069 | 363'284'902 | 11'613'167 | NA | *C. aegagrus* |
| **IRCA-I6-5237** | 103'888'535 | 592'035'177 | 488'146'642 | 199'925'786 | 199'925'786 | 0 | NA | *C. aegagrus* |
| **IRCA-K12-0005** | 89'520'127 | 765'249'396 | 675'729'269 | 325'751'977 | 325'751'977 | 0 | NA | *C. aegagrus* |
| **IRCA-M12-0008** | 99'873'735 | 310'410'173 | 210'536'438 | 36'558'518 | 36'558'518 | 0 | NA | *C. aegagrus* |
| **fal.Cafal1** | 573'621'042 | 2'314'024'081 | 1'740'403'039 | 549'883'291 | 448'500'903 | 101'382'388 | NA | *C. falconeri* |
| **FRCH-AL-0002** | 111'890'971 | 322'924'190 | 211'033'219 | 92'718'720 | 92'718'720 | 0 | Alpine | *C. hircus* |
| **IRCH-B3-5031** | 53'738'058 | 60'736'586 | 6'998'528 | 0 | 0 | 0 | NA | *C. hircus* |
| **IRCH-C5-5206** | 71'091'645 | 786'587'520 | 715'495'875 | 450'008'904 | 427'535'728 | 22'473'176 | NA | *C. hircus* |
| **IRCH-F3-5044** | 59'526'367 | 299'946'902 | 240'420'535 | 120'980'706 | 120'980'706 | 0 | NA | *C. hircus* |
| **IRCH-F4-5093** | 57'578'725 | 696'091'487 | 638'512'762 | 371'049'415 | 358'859'987 | 12'189'428 | NA | *C. hircus* |
| **IRCH-G5-5185** | 72'654'707 | 87'703'744 | 15'049'037 | 0 | 0 | 0 | NA | *C. hircus* |
| **MOCH-AA10-2195** | 90'182'118 | 271'781'073 | 181'598'955 | 107'678'451 | 92'861'325 | 14'817'126 | landrace | *C. hircus* |
| **MOCH-L10-3100** | 106'103'980 | 981'754'026 | 875'650'046 | 447'547'432 | 426'912'917 | 20'634'515 | landrace | *C. hircus* |
| **MOCH-Q15-1143** | 94'753'992 | 1'103'076'736 | 1'008'322'744 | 639'531'078 | 587'996'693 | 51'534'385 | landrace | *C. hircus* |
| **MOCH-R5-0037** | 76'703'976 | 484'745'657 | 408'041'681 | 245'431'572 | 233'806'996 | 11'624'576 | landrace | *C. hircus* |
| **MOCH-S8-2252** | 96'022'567 | 1'028'257'715 | 932'235'148 | 552'459'778 | 510'277'731 | 42'182'047 | landrace | *C. hircus* |
| **MOCH-V5-3059** | 104'574'015 | 171'184'148 | 66'610'133 | 13'521'240 | 13'521'240 | 0 | landrace | *C. hircus* |

| Individual | 100-500 Kb | >100 kb | >500 kb | >2.5 Mb | 2.5-10 Mb | >10 Mb | Population/ Breed | Species |
|---|---|---|---|---|---|---|---|---|
| | Settings: af.dflt 0.4, M=0.0000012, without Viterbi | | | | | | | |
| **ib.GR0201_al** | 783'649'374 | 1'995'664'565 | 1'212'015'191 | 450'312'926 | 429'182'212 | 21'130'714 | al | *C. ibex* |
| **ib.GR0323_al** | 771'984'838 | 2'074'769'627 | 1'302'784'789 | 493'636'590 | 482'683'987 | 10'952'603 | al | *C. ibex* |
| **ib.GR0442_al** | 744'636'342 | 2'110'321'145 | 1'365'684'803 | 531'212'709 | 507'408'809 | 23'803'900 | al | *C. ibex* |
| **ib.08S.12F_am** | 506'440'668 | 2'215'138'383 | 1'708'697'715 | 886'398'299 | 757'676'854 | 128'721'445 | am | *C. ibex* |
| **ib.06S.12M_am** | 535'367'583 | 2'182'318'972 | 1'646'951'389 | 843'786'402 | 769'462'039 | 74'324'363 | am | *C. ibex* |
| **ib.22S.12F_am** | 504'894'041 | 2'177'064'831 | 1'672'170'790 | 924'503'363 | 788'231'278 | 136'272'085 | am | *C. ibex* |
| **ib.BE0309_bo** | 731'956'217 | 2'104'565'569 | 1'372'609'352 | 567'146'789 | 519'862'556 | 47'284'233 | bo | *C. ibex* |
| **ib.BE0407_bo** | 682'427'109 | 2'062'002'272 | 1'379'575'163 | 673'696'514 | 593'848'718 | 79'847'796 | bo | *C. ibex* |
| **ib.BE0463_bo** | 766'167'748 | 2'112'275'126 | 1'346'107'378 | 443'543'188 | 421'161'453 | 22'381'735 | bo | *C. ibex* |
| **ib.BE0010_br** | 746'218'372 | 2'155'341'482 | 1'409'123'110 | 431'098'700 | 420'768'536 | 10'330'164 | br | *C. ibex* |
| **ib.BE0283_br** | 648'107'833 | 2'231'657'101 | 1'583'549'268 | 543'880'783 | 505'040'297 | 38'840'486 | br | *C. ibex* |
| **ib.OW0002_br** | 725'021'312 | 2'177'987'516 | 1'452'966'204 | 477'831'355 | 454'859'576 | 22'971'779 | br | *C. ibex* |
| **ib.GPR21C_gp** | 876'341'653 | 2'008'306'378 | 1'131'964'725 | 363'481'599 | 352'877'217 | 10'604'382 | gp | *C. ibex* |
| **ib.GPV05C_gp** | 772'343'629 | 2'102'615'900 | 1'330'272'271 | 453'794'181 | 441'906'799 | 11'887'382 | gp | *C. ibex* |
| **ib.GPV18C_gp** | 729'634'379 | 2'175'335'815 | 1'445'701'436 | 434'715'929 | 424'683'247 | 10'032'682 | gp | *C. ibex* |
| **ib.GPO03D_gp** | 784'511'145 | 2'060'500'449 | 1'275'989'304 | 379'316'440 | 340'167'171 | 39'149'269 | gp | *C. ibex* |
| **ib.GPO39B_gp** | 828'638'496 | 2'037'365'688 | 1'208'727'192 | 316'277'004 | 316'277'004 | 0 | gp | *C. ibex* |
| **ib.GPV09K_gp** | 814'803'919 | 2'028'835'721 | 1'214'031'802 | 354'293'490 | 354'293'490 | 0 | gp | *C. ibex* |
| **ib.GPV13G_gp** | 791'183'333 | 2'006'827'487 | 1'215'644'154 | 342'628'909 | 342'628'909 | 0 | gp | *C. ibex* |
| **ib.NW0045_ob** | 683'383'101 | 2'055'828'846 | 1'372'445'745 | 595'495'883 | 564'218'995 | 31'276'888 | ob | *C. ibex* |
| **ib.NW0061_ob** | 722'075'711 | 2'094'887'534 | 1'372'811'823 | 622'284'739 | 573'736'535 | 48'548'204 | ob | *C. ibex* |
| **ib.UR0036_ob** | 617'068'750 | 2'158'114'200 | 1'541'045'450 | 767'497'546 | 697'638'277 | 69'859'269 | ob | *C. ibex* |
| **ib.VS0139_pl** | 758'740'285 | 2'145'084'721 | 1'386'344'436 | 449'421'486 | 415'295'277 | 34'126'209 | pl | *C. ibex* |
| **ib.VS1121_pl** | 731'045'337 | 2'127'252'036 | 1'396'206'699 | 534'826'955 | 469'751'317 | 65'075'638 | pl | *C. ibex* |
| **ib.VS1124_pl** | 873'505'705 | 1'924'745'798 | 1'051'240'093 | 275'850'892 | 275'850'892 | 0 | pl | *C. ibex* |
| **ib.GR0140_rh** | 669'983'274 | 2'001'957'025 | 1'331'973'751 | 575'091'026 | 550'389'677 | 24'701'349 | rh | *C. ibex* |
| **ib.VS0079_wh** | 734'275'706 | 2'152'755'412 | 1'418'479'706 | 502'302'612 | 454'745'847 | 47'556'765 | wh | *C. ibex* |
| **ib.VS0081_wh** | 685'673'558 | 2'142'824'732 | 1'457'151'174 | 579'985'076 | 541'409'137 | 38'575'939 | wh | *C. ibex* |
| **ib.VS0031_wh** | 759'969'197 | 2'029'774'334 | 1'269'805'137 | 476'045'828 | 428'999'287 | 47'046'541 | wh | *C. ibex* |
| **nu.ibx20** | 645'141'286 | 764'369'563 | 119'228'277 | 4'946'168 | 4'946'168 | 0 | NA | *C. nubiana* |
| **nu.ibx61** | 812'058'278 | 1'328'614'042 | 516'555'764 | 260'303'299 | 260'303'299 | 0 | NA | *C. nubiana* |
| **py.Z5_hb** | 815'478'576 | 1'896'275'992 | 1'080'797'416 | 493'413'944 | 408'045'097 | 85'368'847 | hb | *C. pyrenaica* |
| **py.M518_sn** | 332'469'317 | 2'347'523'839 | 2'015'054'522 | 845'012'933 | 831'829'914 | 13'183'019 | sn | *C. pyrenaica* |
| **py.V53_vi** | 819'670'708 | 1'640'855'629 | 821'184'921 | 186'497'356 | 186'497'356 | 0 | vi | *C. pyrenaica* |
| **py.V66_vi** | 736'060'943 | 1'618'831'409 | 882'770'466 | 256'608'832 | 227'041'412 | 29'567'420 | vi | *C. pyrenaica* |
| **si.Casi1** | 255'207'615 | 408'162'919 | 152'955'304 | 75'253'542 | 51'229'618 | 24'023'924 | NA | *C. sibirica* |
| **si.Casi2** | 1'006'207'772 | 1'244'095'194 | 237'887'422 | 38'624'317 | 38'624'317 | 0 | NA | *C. sibirica* |
| **IRCA-C3-1001** | 149'214'281 | 673'260'511 | 524'046'230 | 367'252'324 | 276'738'287 | 90'514'037 | NA | *C. aegagrus* |
| **IRCA-G2-5063** | 195'537'312 | 445'690'958 | 250'153'646 | 95'027'576 | 84'593'640 | 10'433'936 | NA | *C. aegagrus* |
| **IRCA-I11-0001** | 68'939'368 | 846'990'938 | 778'051'570 | 541'984'632 | 454'595'429 | 87'389'203 | NA | *C. aegagrus* |
| **IRCA-I6-5237** | 120'388'494 | 641'820'877 | 521'432'383 | 336'343'434 | 324'887'694 | 11'455'740 | NA | *C. aegagrus* |
| **IRCA-K12-0005** | 74'421'878 | 779'866'962 | 705'445'084 | 465'174'475 | 384'455'993 | 80'718'482 | NA | *C. aegagrus* |
| **IRCA-M12-0008** | 93'266'760 | 338'597'836 | 245'331'076 | 97'126'960 | 97'126'960 | 0 | NA | *C. aegagrus* |
| **fal.Cafal1** | 86'633'715 | 2'391'817'852 | 2'305'184'137 | 1'702'176'670 | 1'340'824'547 | 361'352'123 | NA | *C. falconeri* |
| **FRCH-AL-0002** | 193'289'812 | 416'876'710 | 223'586'898 | 150'003'969 | 150'003'969 | 0 | Alpine | *C. aegagrus hircus* |
| **IRCH-B3-5031** | 102'635'552 | 113'614'146 | 10'978'594 | 0 | 0 | 0 | NA | *C. aegagrus hircus* |
| **IRCH-C5-5206** | 80'643'047 | 824'805'115 | 744'162'068 | 614'532'865 | 537'104'184 | 77'428'681 | NA | *C. aegagrus hircus* |
| **IRCH-F3-5044** | 89'815'929 | 342'912'438 | 253'096'509 | 181'835'799 | 165'180'180 | 16'655'619 | NA | *C. aegagrus hircus* |
| **IRCH-F4-5093** | 71'749'818 | 727'932'777 | 656'182'959 | 511'594'515 | 395'940'769 | 115'653'746 | NA | *C. aegagrus hircus* |
| **IRCH-G5-5185** | 124'475'913 | 147'157'579 | 22'681'666 | 2'708'642 | 2'708'642 | 0 | NA | *C. aegagrus hircus* |
| **MOCH-AA10-2195** | 146'220'987 | 340'288'629 | 194'067'642 | 151'554'435 | 114'959'593 | 36'594'842 | landrace | *C. aegagrus hircus* |
| **MOCH-L10-3100** | 120'386'963 | 1'032'323'265 | 911'936'302 | 708'108'515 | 574'495'587 | 133'612'928 | landrace | *C. aegagrus hircus* |
| **MOCH-Q15-1143** | 118'049'121 | 1'156'701'237 | 1'038'652'116 | 878'378'405 | 615'022'573 | 263'355'832 | landrace | *C. aegagrus hircus* |
| **MOCH-R5-0037** | 115'316'268 | 542'454'919 | 427'138'651 | 341'618'532 | 278'102'708 | 63'515'824 | landrace | *C. aegagrus hircus* |
| **MOCH-S8-2252** | 114'340'854 | 1'082'372'825 | 968'031'971 | 800'852'904 | 593'627'379 | 207'225'525 | landrace | *C. aegagrus hircus* |
| **MOCH-V5-3059** | 178'078'661 | 256'774'033 | 78'695'372 | 21'118'017 | 21'118'017 | 0 | landrace | *C. aegagrus hircus* |
| **hi.PCGA06** | 828'868'429 | 990'581'676 | 161'713'247 | 91'561'239 | 91'561'239 | 0 | Peacock | *C. aegagrus hircus* |
| **FRCH-SA-0001** | 172'403'707 | 326'762'731 | 154'359'024 | 90'564'251 | 57'648'233 | 32'916'018 | Saanen | *C. aegagrus hircus* |
| **ITCH-SA-0001** | 198'698'577 | 342'352'391 | 143'653'814 | 90'281'895 | 78'559'944 | 11'721'951 | Saanen | *C. aegagrus hircus* |
| **ITCH-SA-0005** | 186'683'295 | 307'887'462 | 121'204'167 | 79'505'337 | 79'505'337 | 0 | Saanen | *C. aegagrus hircus* |

**Table S4**: Deleterious mutations segregating among and within species. SNPs were filtered as detailed in the Methods. For Alpine and Iberian ibex, we jointly defined a subset of SNPs for which either the Alpine and/or the Iberian ibex carried the derived allele. The SNPs include only sites for which no other species than Alpine and Iberian ibex carried the derived allele. See Methods for details on ancestral/derived state identification procedures.

| **Variable sites within groups/species:** | **All species** | | | *C. ibex* | | *C. ibex* derived | | *C. pyrenaica* | | *C. pyrenaica* derived | | *C. hircus* | | *C. aegagrus* | | *C. sibirica* | | *C. falconeri* | | *C. nubiana* | |
|---|---|---|---|---|---|---|---|---|---|---|---|---|---|---|---|---|---|---|---|---|---|
| | number of SNPs | % | | number of SNPs | % | number of SNPs | % | number of SNPs | % | number of SNPs | % | number of SNPs | % | number of SNPs | % | number of SNPs | % | number of SNPs | % | number of SNPs | % |
| **Total** | **370'853** | | | **22'255** | | **16'545** | | **29'100** | | **23'656** | | **123'791** | | **83'088** | | **58'850** | | **11'062** | | **43'141** | |
| **SnpEff category** | | | | | | | | | | | | | | | | | | | | | |
| Stop gained | 444 | 0.12 | | 60 | 0.27 | 38 | 0.23 | 60 | 0.21 | 39 | 0.16 | 163 | 0.13 | 108 | 0.13 | 70 | 0.12 | 29 | 0.26 | 58 | 0.13 |
| High | 614 | 0.17 | | 69 | 0.31 | 41 | 0.25 | 72 | 0.25 | 47 | 0.2 | 241 | 0.19 | 156 | 0.19 | 102 | 0.17 | 37 | 0.33 | 76 | 0.18 |
| Moderate | 70'873 | 19.1 | | 5157 | 23.2 | 3754 | 22.7 | 6546 | 22.5 | 5254 | 22.2 | 24540 | 19.8 | 15841 | 19.1 | 10491 | 17.8 | 2358 | 21.3 | 8457 | 19.6 |
| Low | 122'677 | 33.1 | | 7186 | 32.3 | 5068 | 30.6 | 9236 | 31.7 | 7214 | 30.5 | 41614 | 33.6 | 28435 | 34.2 | 20332 | 34.5 | 3612 | 32.7 | 14631 | 33.9 |
| Modifier | 176'689 | 47.6 | | 9843 | 44.2 | 7682 | 46.4 | 13246 | 45.5 | 11141 | 47.1 | 57396 | 46.4 | 38656 | 46.5 | 27925 | 47.5 | 5055 | 45.7 | 19977 | 46.3 |
| **Range of GERP conservation score** | | | | | | | | | | | | | | | | | | | | | |
| >6 | 4'343 | 1.2 | | 312 | 1.4 | 218 | 1.3 | 369 | 1.3 | 293 | 1.2 | 1494 | 1.2 | 989 | 1.2 | 695 | 1.2 | 125 | 1.1 | 507 | 1.2 |
| 4 - 6 | 96'528 | 26.0 | | 5759 | 25.9 | 4277 | 25.9 | 7545 | 25.9 | 6151 | 26.0 | 32303 | 26.1 | 21652 | 26.1 | 15493 | 26.3 | 2874 | 26.0 | 11148 | 25.8 |
| 2 - 4 | 96'967 | 26.1 | | 6040 | 27.1 | 4598 | 27.8 | 7901 | 27.2 | 6492 | 27.4 | 32525 | 26.3 | 21437 | 25.8 | 14748 | 25.1 | 2814 | 25.4 | 11205 | 26.0 |
| -2 - 2 | 173'015 | 46.7 | | 10144 | 45.6 | 7452 | 45.0 | 13285 | 45.7 | 10720 | 45.3 | 57469 | 46.4 | 39010 | 47.0 | 27914 | 47.4 | 5249 | 47.5 | 20281 | 47.0 |

**Table S5**: Identical to Table S4 except showing indels instead of SNPs.

| **Variable sites within groups/species:** | **All species** | | | *C. ibex* | | *C. ibex* derived | | *C. pyrenaica* | | *C. pyrenaica* derived | | *C. hircus* | | *C. aegagrus* | | *C. sibirica* | | *C. falconeri* | | *C. nubiana* | |
|---|---|---|---|---|---|---|---|---|---|---|---|---|---|---|---|---|---|---|---|---|---|
| | number of indels | % | | number of indels | % | number of indels | % | number of indels | % | number of indels | % | number of indels | % | number of indels | % | number of indels | % | number of indels | % | number of indels | % |
| **Total** | **27'039** | | | **3'618** | | **2'012** | | **2'981** | | **1'960** | | **8'827** | | **5'730** | | **4'746** | | **1'052** | | **3'632** | |
| **SnpEff category** | | | | | | | | | | | | | | | | | | | | | |
| Stop gained | 32 | 0.12 | | 4 | 0.11 | 3 | 0.15 | 2 | 0.07 | 2 | 0.1 | 9 | 0.1 | 8 | 0.14 | 11 | 0.23 | 0 | 0 | 0 | 0 |
| High | 1'890 | 6.99 | | 651 | 17.99 | 280 | 13.92 | 354 | 11.88 | 119 | 6.07 | 825 | 9.35 | 500 | 8.73 | 394 | 8.3 | 186 | 17.68 | 375 | 10.32 |
| Moderate | 1'273 | 4.7 | | 248 | 6.9 | 119 | 5.9 | 178 | 6.0 | 103 | 5.3 | 491 | 5.6 | 265 | 4.6 | 215 | 4.5 | 62 | 5.9 | 185 | 5.1 |
| Low | 95 | 0.4 | | 12 | 0.3 | 5 | 0.2 | 14 | 0.5 | 11 | 0.6 | 25 | 0.3 | 17 | 0.3 | 19 | 0.4 | 3 | 0.3 | 15 | 0.4 |
| Modifier | 23'781 | 88 | | 2'707 | 74.8 | 1608 | 79.9 | 2'435 | 81.7 | 1727 | 88.1 | 7486 | 84.8 | 4'948 | 86.4 | 4'118 | 86.8 | 801 | 76.1 | 3'057 | 84.2 |
| **Range of GERP conservation score** | | | | | | | | | | | | | | | | | | | | | |
| >6 | 15'141 | 56 | | 1'723 | 47.6 | 987 | 49.1 | 1'551 | 52.0 | 1053 | 53.7 | 4835 | 54.8 | 3'246 | 56.6 | 2'706 | 57.0 | 562 | 53.4 | 2'067 | 56.9 |
| 4 - 6 | 7'358 | 27.2 | | 975 | 26.9 | 534 | 26.5 | 821 | 27.5 | 546 | 27.9 | 2427 | 27.5 | 1524 | 26.6 | 1'259 | 26.5 | 268 | 25.5 | 933 | 25.7 |
| 2 - 4 | 4'357 | 16.1 | | 877 | 24.2 | 470 | 23.4 | 587 | 19.7 | 346 | 17.7 | 1508 | 17.1 | 917 | 16.0 | 750 | 15.8 | 212 | 20.2 | 608 | 16.7 |
| -2 - 2 | 183 | 0.7 | | 43 | 1.2 | 21 | 1.0 | 22 | 0.7 | 15 | 0.8 | 57 | 0.6 | 43 | 0.8 | 31 | 0.7 | 10 | 1.0 | 24 | 0.7 |

**Table S6**: Predicted protein functions of genes for which at least one homozygote was found for the derived, high-impact allele.

| Gene name | Predicted protein function | Polymorphism within species | SnpEff stop-gained |
|---|---|---|---|
| **Alpine ibex (*C. ibex*)** | | | |
| **KNG1** | kininogen-1 isoforms X1-3 | Segregating | yes |
| **LOC102181723** | protein FAM188B2 | Fixed | yes |
| **LOC102181167** | 60S ribosomal protein L10a-like | Fixed | yes |
| **DIS3L2** | DIS3-like exonuclease 2 | Segregating | yes |
| **EPHA8** | ephrin type-A receptor 8 | Segregating | yes |
| **LOC102178786** | coiled-coil domain-containing protein 30-like | Segregating | no |
| **LOC102180106** | unknown | Segregating | no |
| **OVCH1** | ovochymase-1 | Fixed | yes |
| **ERAP2** | endoplasmic reticulum aminopeptidase 2 | Segregating | no |
| **TLN2** | talin-2 | Segregating | yes |
| **POLR1A** | DNA-directed RNA polymerase I subunit RPA1 | Segregating | yes |
| **LOC102188952** | unknown | Segregating | yes |
| **OOEP** | oocyte-expressed protein homolog | Segregating | yes |
| **FOXR1** | Forkhead box protein R1 | Segregating | yes |
| **PNMAL1** | PNMA-like protein 1 | Segregating | yes |
| **RAD1** | cell cycle checkpoint protein RAD1 | Fixed | no |
| **KIF9** | kinesin-like protein KIF9 isoforms X1-3 | Fixed | yes |
| **LOC102179713** | C-C chemokine receptor type 1-like | Segregating | yes |
| **CD2BP2** | CD2 antigen cytoplasmic tail-binding protein 2 isoforms X1-2 | Fixed | yes |
| **LOC102189088** | zinc finger protein 205 isoform X1 | Fixed | no |
| **LOC102188472** | synaptotagmin-15 isoforms X1-4 | Segregating | yes |
| **LOC102175191** | solute carrier family 22 member 9-like | Fixed | yes |
| **Iberian Ibex (*C. pyrenaica*)** | | | |
| **LOC102181723** | protein FAM188B2 | Segregating | yes |
| **LOC102181167** | 60S ribosomal protein L10a-like | Segregating | yes |
| **CAPZB** | F-actin-capping protein subunit beta | Segregating | yes |
| **TMBIM6** | bax inhibitor 1 | Segregating | yes |
| **LOC102191021** | protein HP-25 homolog 2 | Segregating | yes |
| **OVCH1** | ovochymase-1 | Fixed | yes |
| **LOC102173270** | C-type lectin domain family 2 member B | Segregating | yes |
| **TLR6** | toll-like receptor 6 | Segregating | yes |
| **TLR6** | toll-like receptor 6 isoform X1 | Segregating | yes |
| **LOC106502316** | zinc finger protein 77-like | Segregating | yes |
| **LOC102191525** | eukaryotic translation initiation factor 3 subunit J | Segregating | yes |
| **TMEM244** | transmembrane protein 244 | Segregating | yes |

**Supplementary File S1**: Script used for individual based simulations.

```
run_mode overwrite
random_seed 30317
logfile logfile.log
root_dir SIMS/sim5000_0.0014_0.01
filename sim

replicates 100
generations 3023
patch_number 12
patch_capacity (@g0 {{1000, 0, 0, 0, 0, 0, 0, 0, 0, 0, 0,0}}, @g3000 {{500, 0, 0, 0, 0, 0, 0, 0, 0, 0, 0,0}}, @g3005 {{80,
    0, 0, 0, 0, 0, 0, 0, 0, 0, 0,0}}, @g3007 {{1000, 0, 0, 0, 0, 0, 0, 0, 0, 0, 0,0}}, @g3010 {{1000, 25, 30, 0, 0, 0, 0, 0, 0,
    0, 20,0}}, @g3012 {{1000, 25, 30, 85, 73, 46, 0, 0, 0, 0, 20,0}}, @g3017 {{1000, 25, 30, 85, 73, 46, 41, 30, 22,
    116, 20, 43}})

ntrl_init 1
breed_selection 2
disperse 3
aging 4
save_stats 5
save_files 6

mating_system 1
mean_fecundity 5
dispersal_connectivity_matrix (@g0 {{1}{2}{3}{4}{5}{6}{7}{8}{9}{10}{11}{12}},@g3010
    {{1,2,3,11}{2}{3}{4}{5}{6}{7}{8}{9}{10}{11}{12}},@g3011
    {{1}{2}{3}{4}{5}{6}{7}{8}{9}{10}{11}{12}},@g3012
    {{1}{2,6,4,5}{3,6,4,5}{4}{5}{6}{7}{8}{9}{10}{11}{12}},@g3013
    {{1}{2}{3}{4}{5}{6}{7}{8}{9}{10}{11}{12}},@g3017
    {{1,5}{2}{3}{4,7}{5,8}{6,9,12,10}{7}{8}{9}{10}{11}{12}},@g3018
    {{1}{2}{3}{4}{5}{6}{7}{8}{9}{10}{11}{12,9}},@g3019 {{1}{2}{3}{4}{5}{6}{7}{8}{9}{10}{11}{12}})
dispersal_reduced_matrix (@g0 {{1}{1}{1}{1}{1}{1}{1}{1}{1}{1}{1}{1}},@g3010
    {{0.854,0.060,0.076,0.010}{1}{1}{1}{1}{1}{1}{1}{1}{1}{1}{1}},@g3011
    {{1}{1}{1}{1}{1}{1}{1}{1}{1}{1}{1}{1}},@g3012
    {{1}{0.035,0.400,0.353,0.212}{0.333,0.49,0.059,0.118}{1}{1}{1}{1}{1}{1}{1}{1}{1}},@g3013
    {{1}{1}{1}{1}{1}{1}{1}{1}{1}{1}{1}{1}},@g3017
    {{0.996,0.004}{1}{1}{0.910,0.090}{0.839,0.161}{0.361,0.051,0.217,0.371}{1}{1}{1}{1}{1}{1}},@g3018
    {{1}{1}{1}{1}{1}{1}{1}{1}{1}{1}{1}{0.754,0.246}},@g3019 {{1}{1}{1}{1}{1}{1}{1}{1}{1}{1}{1}{1}})

## SELECTION TRAITS
selection_trait delet
selection_model direct
selection_fitness_model absolute

## NEUTRAL TRAITS
ntrl_loci 500
ntrl_all 2
ntrl_mutation_rate 5e-05
ntrl_recombination_rate 0.5
ntrl_mutation_model 1
ntrl_init_patch_freq {{0.0014}}

## DELETERIOUS TRAITS
delet_loci 5000
delet_mutation_rate 5e-05
delet_backmutation_rate 5e-07 # default
delet_recombination_rate 0.5
```

delet_mutation_model 1
delet_fitness_model 1

delet_init_freq 0.0014
delet_effects_distribution gamma
delet_effects_mean 0.01
delet_dominance_mean 0.37

delet_effects_dist_param1 0.3

### OUTPUT
stat_dir stats
stat_log_time (@g0 1000, @g3000 1)
stat adlt.demography pop.patch fecundity migrants.patch migrants adlt.delet viability ntrl.freq
stat_output_compact
stat_output_CSV

ntrl_save_genotype FSTAT
ntrl_output_dir ntrl_geno
ntrl_output_logtime {{1, 1000, 2000, 2999, 3000, 3001, 3002, 3003, 3004, 3005, 3006, 3007, 3008, 3009, 3010, 3011, 3012, 3013, 3014, 3015, 3016, 3017, 3018, 3019, 3020, 3021, 3022, 3023}}

delet_save_genotype
delet_genot_dir delet_geno
delet_genot_logtime {{1, 1000, 2000, 2999, 3000, 3001, 3002, 3003, 3004, 3005, 3006, 3007, 3008, 3009, 3010, 3011, 3012, 3013, 3014, 3015, 3016, 3017, 3018, 3019, 3020, 3021, 3022, 3023}}
